# Transcription elongation defects link oncogenic splicing factor mutations to targetable alterations in chromatin landscape

**DOI:** 10.1101/2023.02.25.530019

**Authors:** Prajwal C. Boddu, Abhishek K. Gupta, Rahul Roy, Barbara De La Pena Avalos, Ane Olazabal Herrero, Nils Neuenkirchen, Joshua T. Zimmer, Namrata S. Chandhok, Darren King, Yasuhito Nannya, Seishi Ogawa, Haifan Lin, Matthew D. Simon, Eloise Dray, Gary Kupfer, Amit K. Verma, Karla M. Neugebauer, Manoj M. Pillai

## Abstract

Transcription and splicing of pre-messenger RNA are closely coordinated, but how this functional coupling is disrupted in human disease remains unexplored. Here, we investigated the impact of non-synonymous mutations in SF3B1 and U2AF1, two splicing factors commonly mutated in cancer, on transcription. We find that the mutations impair RNA Polymerase II (RNAPII) transcription elongation along gene bodies leading to transcription-replication conflicts, replication stress and altered chromatin organization. This elongation defect is linked to disrupted pre-spliceosome assembly due to impaired protein-protein interactions of mutant SF3B1. Through an unbiased screen, we identified epigenetic factors in the Sin3/HDAC complex, which, when modulated, normalize transcription defects and their downstream effects. Our findings shed light on the mechanisms by which oncogenic mutant spliceosomes impact chromatin organization through their effects on RNAPII transcription elongation and present a rationale for targeting the Sin3/HDAC complex as a potential therapeutic strategy.

**HIGHLIGHTS:**

– Oncogenic mutations in SF3B1 and U2AF1 cause a gene body RNAPII transcription elongation defect
– The elongation defect is linked to impaired assembly of early spliceosome complexes and leads to replication stress and changes to chromatin landscape
– RNAPII elongation defects in SF3B1^K700E^ are normalized by modulating epigenetic factors but 3’ cryptic splicing events are not reversed
– Targeting the Sin3/HDAC pathway to normalize RNAPII elongation defect is a potential therapeutic approach in these disorders

**GRAPHICAL ABSTRACT:** 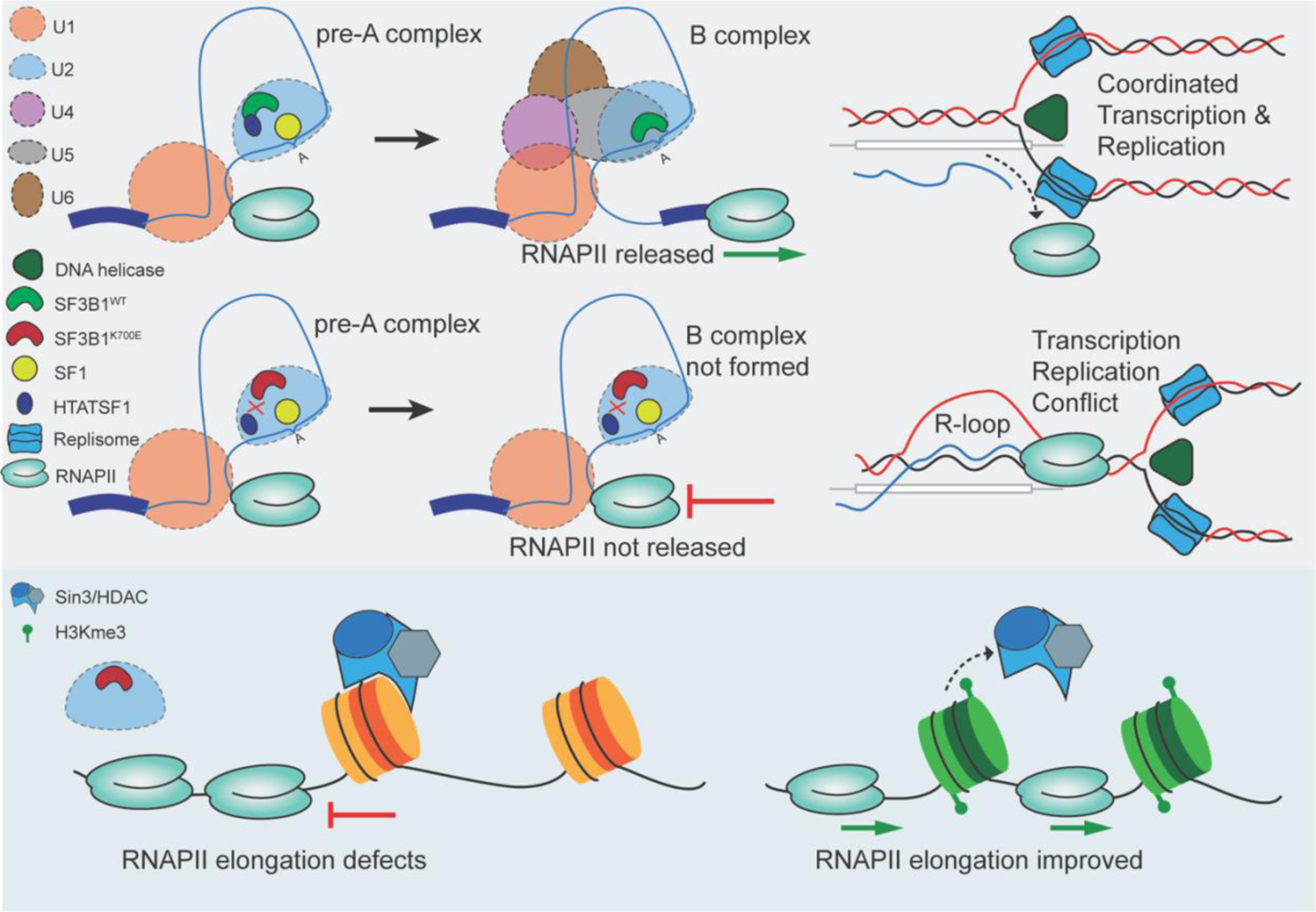

## INTRODUCTION

Although traditionally seen as discrete processes driven by distinct multi-protein complexes, we now recognize that RNA transcription and splicing are closely coordinated^1^, ^2^. Accordingly, the spliceosome assembles and removes introns while the nascent transcript is still growing longer, although not strictly in temporal order^3^, ^4^. Perturbing splicing (through inhibitors such as Pladienolide B (PlaB)) can affect many aspects of transcription and RNA processing^5^, ^6^. Similarly, slowing the transcriptional machinery changes the selection of alternative splice sites^7^. Despite such close functional coordination, disruption of co-transcriptional splicing as a basis of human disease has not been explored.

Recurrent mutations in select RNA splicing factor (SF) genes, such as *SF3B1*, *U2AF1, SRSF2* and *ZRSR2*, are drivers of oncogenesis in many cancers, but are most prevalent in myelodysplastic syndromes (MDS) and acute myeloid leukemia (AML)^8^, ^9^, ^10^. The SF mutations are usually heterozygous, non-synonymous, and largely mutually exclusive^11^ suggesting shared downstream mechanisms.

SF3B1, the most mutated SF, is a component of U2 snRNP that stabilizes the interaction of U2 snRNA with branch point (BP)^12^. The mutations are thought to change its confirmation leading to reduced BP fidelity and selection of cryptic 3’splice sites (SSs)^13^. Altered interactions of mutant SF3B1 with accessory splicing proteins such as SUGP1^14^, DDX42, and DDX46^15^ are thought to underlie these splicing changes. Such alternative splicing (AS) events, demonstrable in genes such as ABCB7^16^, MAP3K7^17^, and BRD9^18^, are speculated to contribute to different aspects of SF3B1-mutant disease, such as clonal evolution and dyserythropoiesis in MDS. Changes to AS, while often demonstrable, are generally modest in degree (low change in isoform ratios or low Percent Spliced-In values), and are not consistent across independent models or patient sample datasets^8^, ^18^, ^19^. Importantly, patterns of AS changes common to different SF mutations have not yet emerged. These, and the observation that SF mutations are largely mutually exclusive, has prompted investigation into alternate mechanisms that do not rely on mutant-specific changes in isoform abundance.

In this study, we explored the mechanisms by which mutant spliceosomes dysregulate transcription kinetics. We show that mutations in SF3B1 and U2AF1 primarily dysregulate RNAPII gene body elongation due to altered assembly of early spliceosomes, leading to replication stress and chromatin re-organization. Altered RNAPII elongation was corrected by modulation of specific epigenetic pathways, underscoring the interplay between dysregulated transcription and chromatin organization, and identifies targets of therapeutic relevance.

## RESULTS

### SF3B1^K700E^ redistributes RNAPII from promoters to gene bodies

To explore the functional and pathologic relevance of disruption of co-transcriptional splicing in SF mutant disease context, we generated isogenic K562 cell lines which express mutant alleles upon induced expression of Cre recombinase ^20^(Fig 1A). SF3B1^K700E^ and U2AF1^S34F^ mutants and corresponding control clones with synonymous mutations were generated. ChIP-seq for RNAPII utilizing a pan-RNAPII antibody (NTD) that recognizes RNAPII irrespective of its activation status (FigS1A) was performed to determine distribution of RNAPII in SF3B1^K700E^. Metagene plot analysis of average RNAPII occupancy in highly expressed genes in K562 cells (determined per methodology in^5^) revealed a redistribution of RNAPII in SF3B1^K700E^ into gene bodies (Fig1B). RNAPII density over the promoter/transcription start site (TSS) was reduced while density across the gene body progressively increased culminating at the transcription termination site (TTS or transcription end site (TES)). Representative density plots of such redistribution are shown in Fig1C. The change was also captured by the traveling ratio (TR)^21^, ^22^, which compares read density in the promoter region to that in gene body (Fig1D). ChIP-seq for elongation-specific RNAPII (RNAPII Ser2P) (FigS1B) also showed a marked redistribution of RNAPII occupancy, suggesting a primary failure of RNAPII elongation in SF3B1^K700E^ cells (Fig1E, FigS1C). To determine gene-specific features that determine these RNAPII elongation defects, we performed unsupervised clustering of delta TR (difference in traveling ratio), which resulted in 3 distinct clusters (Fig1F, FigS1D). We then determined multiple attributes of the genes within these individual clusters. Notably, the clusters differed significantly with respect to introns per gene and total gene length in a consistent trend (genes that were longer or contained more introns had more RNAPII redistribution) (Fig1G). Other parameter such as gene expression levels and distance to neighboring genes did not follow a consistent trend across the clusters (FigS1E). Cluster 3, the one with least difference in TR between SF3B1^K700E^ and SF3B1^WT^was significantly enriched for intronless genes, containing over half of the intronless genes (102/200) in the dataset (Fig1H). Gene length and intron number independently correlated with delta TR, confirming these results (FigS1F, FigS1G). To additionally determine how the transcription elongation behavior changes at the exon-intron boundaries, we performed static binning on the RNAPII NTD ChIP data (instead of dynamic binning used in gene body length normalization in metagene plots). As shown in Fig1I, SF3B1^K700E^ cells showed a relative decrease in RNAPII density downstream of TSS followed by a sharp increase in the immediate gene bodies (note the red-to-blue region after the TSS in the heatmap; Fig1I). This transition zone occurred before the (TSS+0.5kb) region, coinciding with the first exon-intron junction boundary (Fig1J). To determine if RNAPII enrichment was selective to the location of the intron in the gene body, RNAPII ChIP-seq densities for the first, second, and terminal introns were independently analyzed, and were noted to show a similar profile (FigS1H). Together, we conclude that RNAPII in SF3B1^K700E^ cells redistributes from promoters to gene bodies, in a manner dependent on the cumulative distance RNAPII has to traverse (gene length) and the number of spliceosomes assembled along this distance (total introns).

**Figure 1:**
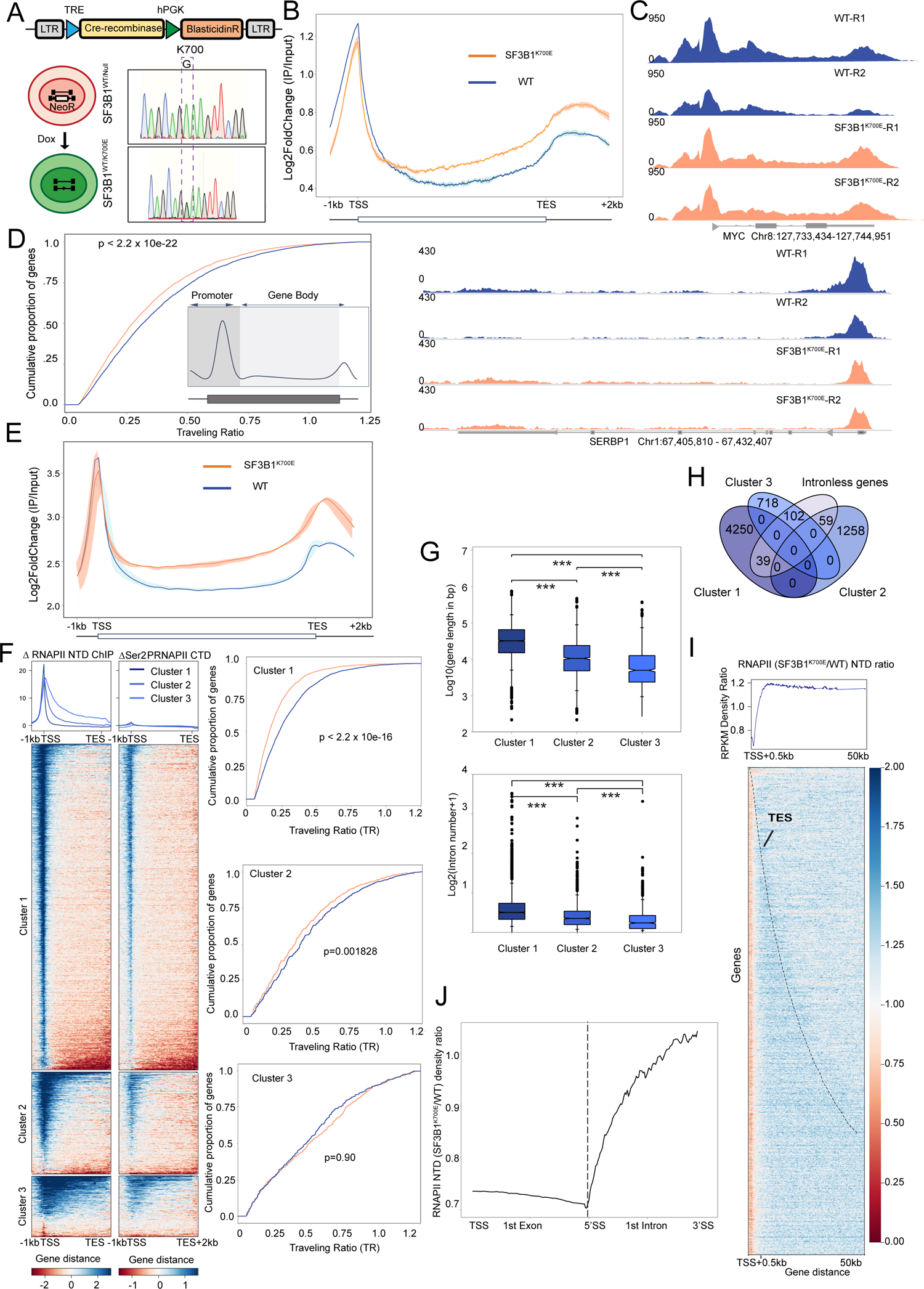
Genome-wide analysis of RNAPII redistribution in SF3B1^K700E^. (A) Inducible *SF3B1* mutant isogenic cell line model system. Upon induction of Cre-recombinase, recombination is near complete at 4 days as confirmed by cDNA sequencing. (B) Metagene plot of ChIP-seq occupancy of RNAPII NTD (log2 fold change IP to input) over 6694 expressed genes in WT and SF3B1^K700E^ cells. Solid lines represent averaged signal, and shaded areas represent the upper and lower signal densities of the individual replicates (n=2 each). Read densities are calculated as RPKM. TSS-transcription start site, TES-transcription end site. (C) Representative genome browser track examples of NTD RNAPII ChIP-seq in WT and SF3B1^K700E^. Y-axis represents read density. (D) Traveling ratio (TR) (calculated as an average of two replicates) in WT and SF3B1^K700E^ cells, p value determined by two-sample K.S. test. Inset: Schematic depicting TR calculation as ratio of promoter (TSS to TSS+300 bp region) to Gene body (TSS+300 bp to TES) read density. (E) Metagene plot of Ser2PRNAPII ChIP-seq (log2 fold change IP to input) as shown in Figure 1B. (n=2 each (F) Heat maps and unsupervised clustering of change in average signal enrichments (Δ) of RNAPII NTD (left) and Ser2PRNAPII CTD (right) up on SF3B1^K700E^ expression in K562 cells. Three Clusters were determined. Heatmap patterns are also reflected in the corresponding ‘TR’ cumulative distribution graphs for each of the 3 clusters (right panel). Cluster 1=4289 genes, Cluster 2=1317 genes, cluster 3=820 genes. (G) Box-whisker plots showing the distribution of gene length (top) and intron number (bottom) for the three gene clusters (two-tailed Mann–Whitney *U*-test. \*\*\**P*<0.0001). (H) Distribution of intronless genes across the 3 gene clusters (total, n=6994 gene dataset; total intronless genes=200). (I) SF3B1^K700E^/WT ratio RNAPII NTD ChIP heatmap with static binning spanning the entire length of all expressed genes up to 50 kb, ordered by increasing gene length. Read densities are in RPKM. (J) Averaged RNAPII enrichment profiles across the first exon-intron junction (total 5118 genes with atleast one intron) in shown as SF3B1^K700E^/WT read density ratio on the Y-axis.

### RNAPII redistribution in SF3B1^K700E^ results from changes in gene body elongation rate

The observed redistribution of RNAPII in SF3B1^K700E^ can arise from two possible scenarios: (a) increased productive elongation resulting from increased transcription initiation and/or premature pause release that reduces pileup of RNAPII at promoters, or (b) defective elongation of RNAPII along the gene body. Since RNAPII ChIP-seq cannot distinguish between the two scenarios, we implemented nascent transcriptomic strategies (including fastGRO-seq and TT-TL-seq) to determine how the redistributed RNAPII transcribes nascent RNA. FastGRO-seq (referred to as GRO-seq from hereon)^23^ is a nuclear run-on assay based on the classical GRO-seq protocol that uses 4-thio ribonucleotides (4sUTPs) to label nascent RNA arising from transcriptionally engaged RNAPII. High correlation between replicates indicates the reproducibility of GRO-seq (FigS4A). Aligned reads were split to sense (+) or antisense (-) and metagene plots were generated. As previously reported, sense read profiles had with a sharp peak at TSS, then tapering through gene body with a second peak at TTS; antisense reads were largely 5’ of the TSS (Fig 2A). Promoter peak densities were similar in SF3B1^K700E^ and SF3B1^WT^, but SF3B1^K700E^ had a higher gene body signal peaking at the TTS (Fig 2A-B).

**Figure 2:**
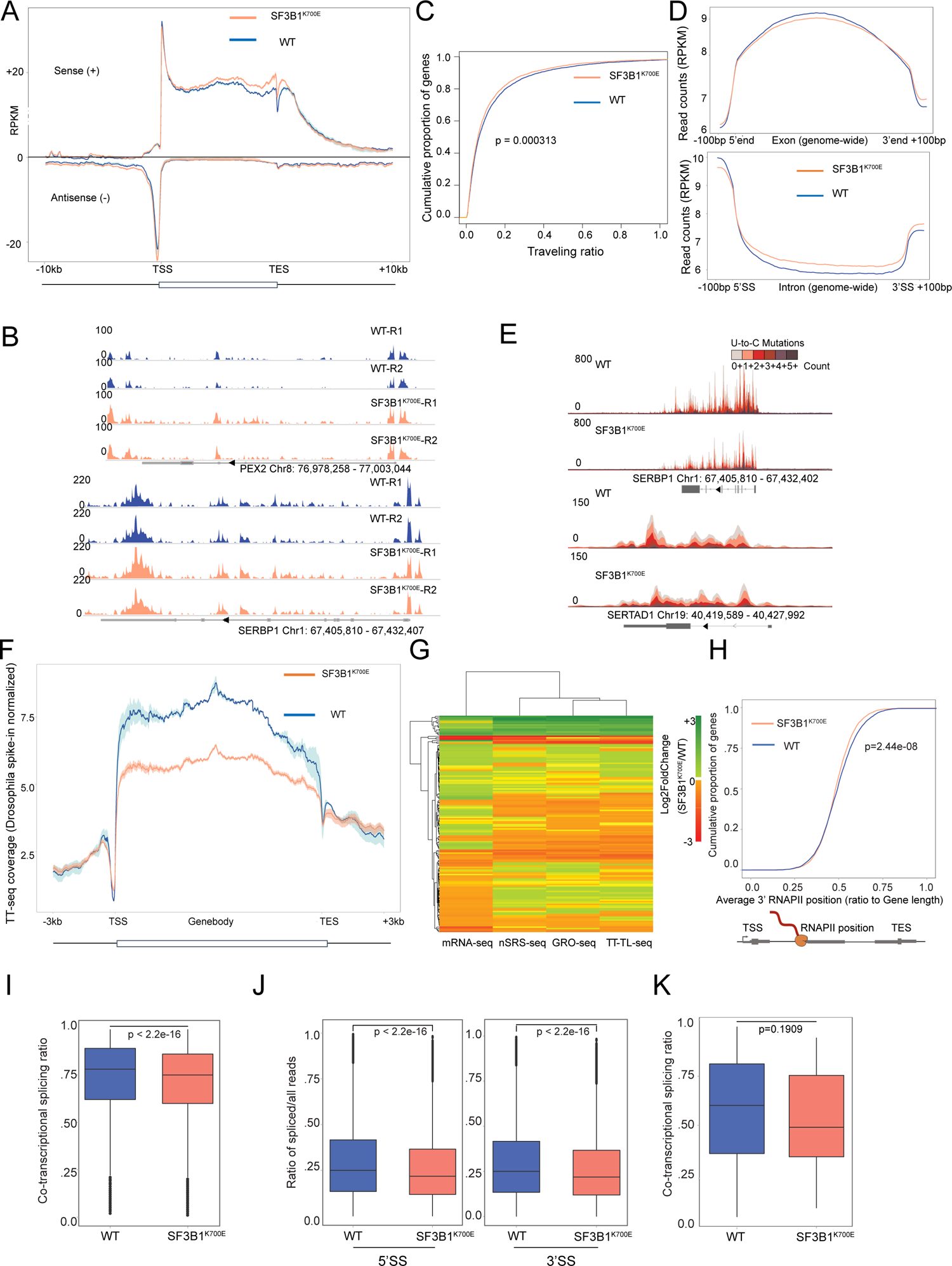
SF3B1^K700E^ induced changes in RNAPII transcription kinetics using nascent transcriptome assays. (A) Metaplot analysis showing GRO-seq densities in WT (n=2) and SF3B1^K700E^-mutant (n=2) K562 cells. (B) Representative genome browser track examples of GRO-seq in K562 WT and SF3B1^K700E^ cells. (C) Traveling Ratio for GRO-seq as performed in 1D. (D) GRO-seq read coverage of introns and exons in WT and SF3B1^K700E^ cells. Coverage normalized to the position −100 bp of the 5’SS (for intron) or to 5’ start (for exons). (E) Genome browser tracks of TT-TL seq data for SERBP1 and SERTAD1 genes after normalization to Drosophila spike-in control. (F) Metaplot of normalized TT-seq densities in WT(n=2) and SF3B1^K700E^-mutant K562 (n=2) cells. (G) Hierarchical clustering of log2 fold changes of read densities along gene bodies of mRNA-seq, nascent RNA-seq, GRO-seq, and TT-TL-seq (H) CDF plot of the median 3’end RNAPII position inferred from 3’read ends of LRS reads (plotted as relative position along transcript length), in WT and SF3B1^K700E^. (I) CoSE for introns (at least 10 reads coverage and 800 bp in length) in SF3B1^K700E^ and WT. P value determined by two-tailed Mann-Whitney-U. (J) Ratio of spliced reads over total unspliced and spliced reads in WT(n=2) and SFB3B1^K700E^(n=2) in GRO-seq. All the introns in the selected 5118 isoforms (from the 6694 gene dataset) were considered. SS-splice sites. (K) CoSE limited to those introns reduced in SF3B1^K700E^ (versus WT) as performed for Figure 2J.

This was also reflected in the TR (Fig 2C). TR calculation analyses of GRO-seq signal were further broken down by cluster 1, 2, and 3 genes (based on Fig 1F), with groups I and II exhibiting a more significant increase of nascent RNA within gene bodies (FigS2B). Relatively higher GRO-seq signal was observed at introns in SF3B1^K700E^, similar to results from RNAPII ChIP-seq (Fig 2D). GRO-seq thus confirms our ChIP-seq results that RNAPII in SF3B1^K700E^ redistributes into the gene body, which when released during run-on transcription results in higher gene body density compared to SF3B1^WT^.

As complementary approaches to GRO-seq, we additionally performed TT-TL-seq^24^ and nascent short-read (nSRS) RNA-seq^25^ in WT and SF3B1^K700E^ (FigS2C-D). TT-TL-seq combines transient transcriptome sequencing (TT-seq, which labels and captures newly synthesized transient RNA) with “time lapse” chemistry (converts 4sU to cytidine analogs, yielding apparent U-to-C mutations that can identify new transcripts in the next generation sequencing datasets (FigS2E). The time lapse component allows precise identification of nascent RNA reads based on the U-to-C conversion (Fig 2E). Metagene analysis showed that the Drosophila-spike-in normalized^26^ TT-seq coverage across the gene body was reduced in SF3B1^K700E^ (Fig 2F), consistent with reduced RNAPII elongation rate and not enhanced/premature pause release. Additionally, differential analysis using DESeq2 showed nearly half of the genes (48.7%) to have reduced coverage across gene body in SF3B1^K700E^ compared to WT; by contrast, only 1.3% of were increased (FigS2F). Notably, the degree of reduction of the TT-seq gene body signal did not correlate with baseline gene expression, a pattern consistent with the lack of a direct interaction effect of SF3B1^K700E^ on the transcribing RNAPII^5^ (FigS2G).

Premature pause release and reduced elongation rate of RNAPII can be distinguished by comparing the relationship between steady state transcript and nascent transcription levels^27^, ^28^. With premature pause release, change to nascent RNA and steady state transcript levels are expected to correlate. Conversely, there will be no correlation between nascent RNA and steady RNA profiles in the latter scenario. Hierarchical clustering of gene expression changes on the four transcriptomic datasets (total RNA-seq, nSRS-seq, GRO-seq and TT-TL-seq) showed the total RNA-seq profile to cluster differently from the 3 nascent RNA-seq measures (Fig 2G). Collectively, our data shows that RNAPII redistribution results from impaired elongation and not premature pause release.

Given the mutual exclusivity of SF mutations, we speculated if a similar RNAPII transcriptional response is demonstrable in U2AF mutant cells. Using an inducible *U2AF1* ^S34F^ isogenic cell line system^20^, we similarly observed redistribution of RNAPII into gene bodies by Ser2P RNAPII ChIP (FigS2H), and a globally reduced gene body signal by TT-TL seq (FigS2I). These data suggest a convergent transcriptional response to distinct SF mutations, which may underlie their mutual exclusivity.

### Long-read sequencing (LRS) reveals reduced co-transcriptional splicing in SF3B1^K700E^

Splicing is co-transcriptional^1^, ^2^ and altered RNAPII elongation rates may affect splice site choice^7^. However, the effect of SF mutations on splicing efficiency itself has not been explored. A common pattern of alternative splicing noted across mutant SF patient samples is a decrease in retained introns (RI)^29^, the molecular basis of which is unclear. Does improved efficiency of co-transcriptional splicing reduce IR, or are the introns removed more efficiently post-transcriptionally? We have previously reported on the use of LRS to determine kinetics of co-transcriptional splicing^1^. LRS provides single molecule-level metrics which Illumina short read sequencing cannot provide. PolyA-depleted, chromatin-associated RNA (nascent RNA) was isolated as previously described^25^ for PacBio library preparation (FigS2J-K). A representative example of aligned LRS reads (ordered by 3’ end position and colored based on splicing status) is shown, in FigS2L. The 3’ end of each LRS read signals the position of RNAPII, and can therefore be used to determine distribution of RNAPII across the gene body. Plotting the median RNAPII position (relative to its normalized gene length as a cumulative distribution function showed a lag (shift towards 5’ position) in median RNAPII positions in SF3B1^K700E^ (Fig 2H). We then divided the genes into the 3 clusters (as determined by RNAPII ChIP-seq data, Fig 1F) and determined the change in 3’ position in each of the clusters. Relative lag of median RNAPII position in SF3B1 ^K700E^ was most pronounced in clusters 1 and 2 (FigS2M), corroborating findings of GRO-seq and RNAPII ChIP-seq.

Co-transcriptional splicing efficiency (CoSE) for individual introns was then determined as a ratio of spliced transcripts to total transcripts (spliced and unspliced) that span the splice junction. Accordingly, a reduced CoSE would indicate a lower rate of co-transcriptional splicing at that junction and vice-versa. Genome-wide comparison of CoSE showed a statistically significant reduction of CoSE in SF3B1^K700E^, compared to WT (Fig 2I). We then determined if the position of introns with relation to transcription start and end sites influenced changes to CoSE. Reductions in CoSE in the middle and final intron subgroups were more pronounced when compared to first introns (FigS2N). We similarly analyzed splicing ratios, separately for the 5’ and 3’ splice sites, using the GRO-seq datasets. Statistically significant reduction of splicing ratios was observed in SF3B1 ^K700E^ compared to WT (when all introns were considered, and when first, middle, and last introns were separately compared) (Fig 2J, FigS2O). Finally, we specifically compared the CoSE of those introns, noted to have reduced IR (identified from RNA-seq using IRFinder^30^), in isogenic SF3B1^K700E^ cells. Interestingly, this group of introns also showed a trend towards reduced CoSE in SF3B1^K700E^ compared to SF3B1^WT^ (Fig 2K). Together, these results show that co-transcriptional splicing is not increased in SF3B1^K700E^. The reduced IR observed in a subset of genes across MDS patient datasets is therefore unlikely to be the result of more efficient co-transcriptional splicing.

### SF3B1^K700E^ disrupts E-to-A spliceosome complex transition

To determine how mutant spliceosomes may affect RNAPII elongation, we considered different possibilities. One would involve a change in expression levels of components of the transcriptional machinery. Analysis of RNA-seq datasets of our isogenic cells, or public databases of patient samples ^31^, ^32^, ^33^ however did not show reproducible changes to gene expression levels or alternative splicing of such transcriptional regulators. Hence, we considered a second possibility – defects in direct interactions between the spliceosome and RNAPII transcription machinery. While such a functional coupling has long been recognized, a direct interaction between RNAPII and the early spliceosome complex (specifically U1 snRNP) was recently reported^2^.

Accordingly, the RNAPII complexes with the early spliceosome and is released only upon formation of the spliceosome B complex. Recent cryoEM studies have also identified the critical initial step in the pre-spliceosome formation (pre-A or E complex) to be the displacement of HTATSF1 from SF3B1 by the helicase DDX46^34, 35^. This ATP-dependent step facilitates displacement of splicing factor-1 (SF1) from the BP, followed by binding of U2AF2 to SF3B1 (UHM to ULM domain interaction), allowing stable binding of U2 snRNP complex to the BP. HEAT domain mutations in SF3B1 (yeast homologue, Hsh155) are known to impair its interaction with DDX46 (yeast homologue, Prp5)^36^, ^15^. We hence determined how the mutation changes SF3B1’s interaction with its partners during early spliceosome assembly. FLAG-SF3B1 (^K700E^ and ^WT^) was transfected in to HEK293T cells, and immunoprecipitated from nuclear lysates after 48 hours. As shown in Fig 3A, FLAG-SF3B1^K700E^ showed reduced binding with HTATSF1, U2AF1, and U2AF2. Since spliceosomes are assembled on chromatin^37^, ^38^, these interactions were also probed on chromatin fractions and the differences were noted to be even more pronounced (Fig 3A). Based on these defective protein-protein interactions early in the spliceosome assembly, we speculated that SF3B1^K700E^ spliceosomes may not transition effectively to later stages (B or later). To test this, we probed nucleoplasmic and chromatin fractions from isogenic K562 cells (SF3B1^K700E^ and SF3B1^WT^) for splicing proteins specific to E (SF3A3, HTATSF1) and B (MFAP1) complex assemblies (Fig 3B). As anticipated, chromatin abundance of HTATSF1 was reduced in SF3B1^K700E^ cells. SF3A3 chromatin levels (E-complex component) were unchanged but MFAP1 (B complex component) was reduced in SF3B1^K700E^ cells, suggestive of a defective transition to B complex. We then determined how spliceosome assembly kinetics of SF3B1^K700E^ differed from SF3B1^WT^ using an *in vitro* splicing assay. Nuclear extracts from HEK293T cells transfected with either SF3B1^WT^ or SF3B1^K700E^ were incubated with p32-labelled RNA-containing splicing reactions for the conditions and timepoints, as indicated in Fig 3C. Transition from E-to A-complex intermediates (as determined by the ratio of A/E complex) was reduced in SF3B1^K700E^ compared to SF3B1^WT^, most evident at the 40 minute and later timepoints (Fig 3C, FigS3A). Given these results, we speculated if disruption of the early pre-spliceosome assembly (through loss of DDX46 or HTATSF1) would result in a transcriptional response similar to that seen with SF3B1^K700E^. We used stable inducible lentiviral shRNA to knock down DDX46 and HTATSF1 in wild-type K562 cells (FigS3B) and performed Ser2P RNAPII ChIP-seq. As shown in Fig 3D, RNAPII was redistributed into the gene bodies upon loss of either DDX46 or HTATSF1, in a pattern similar to that with SF3B1^K700E^. These results were also confirmed using transient siRNA-mediated knockdown of DDX46 and HTATSF1 in K562 cells (FigS3C-D). Finally, we determined if growth arrest of SF3B1^K700E^ cells could be rescued by overexpression of either DDX46 or HTATSF1 (FigS3E). Using doxycycline inducible constructs (FigS3E), we found that overexpression of HTATSF1, but not DDX46, led to partial rescue of SF3B1^K700E^ cells (Fig 3E). Collectively, these findings are consistent with a model of impaired interaction of SF3B1^K700E^ with HTATSF1, which prevents efficient assembly of B complex). Based on the recent cryoEM structures of RNAPII-U1 snRNP interactions, we speculate that this impaired transition prevents release of U1 snRNP and liberation of RNAPII leading to defective RNAPII elongation (Fig 3F).

**Figure 3:**
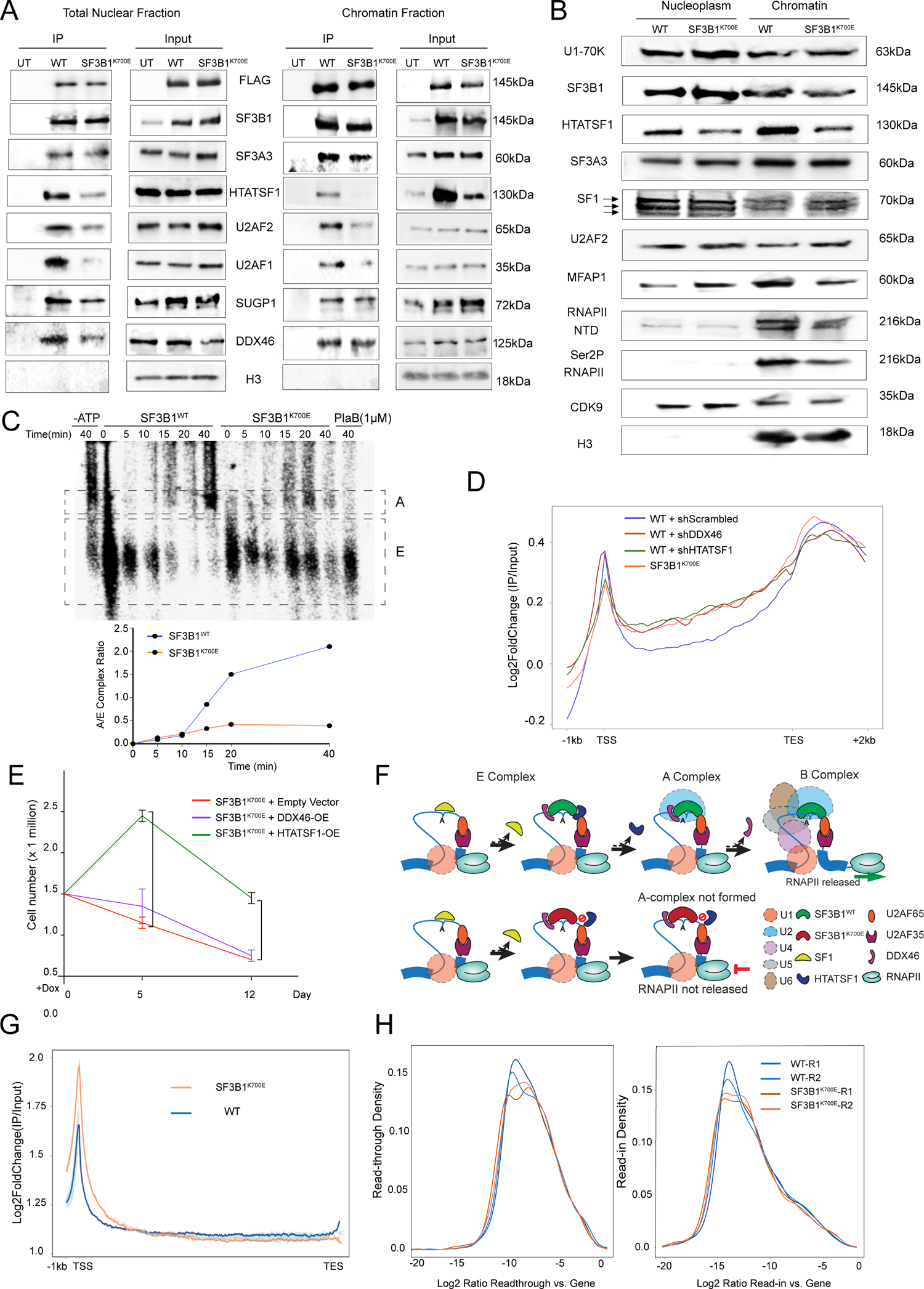
Mechanisms underlying SF3B1^K700E^ induced transcription elongation defect. (A) Immunoblots for proteins in input or IP fractions bound to FLAG-SF3B1 (WT or K700E) along with untransfected controls (UT). Total nuclear extracts are shown on left and chromatin fraction to the right. (B) Immunoblot of indicated proteins in nucleoplasm or chromatin fraction from WT and SF3B1^K700E^ K562 cell lysates. (C) Native gel analyses of spliceosome assembly kinetics on radiolabeled *AdML* pre-mRNA in HEK293T (SF3B1^WT^ and SF3B1^K700E^) nuclear extracts. Expected positions of E and A spliceosome complexes are shown. Relative amounts of A and E complexes were quantified and plotted over time. (D) Meta plots showing log_2_ fold change signal of RNAPII Ser2P ChIP-seq (IP/input) data across the annotated gene transcripts (n=6694) in SF3B1^K700E^ K562 and WT stably integrated with lentiviral constructs expressing inducible shRNA targeting control (shScrambled), DDX46 (shDDX46), and HTATSF1 (shHTATSF1). (E) Growth kinetics of SF3B1^K700E^ cells overexpressing DDX46, HTATSF1 or empty vector control over 12 days (\*\*\**P*<0.0001, \*\**P*<0.001; two-tailed *t*-test). (F) Proposed model for SF3B1^K700E^-induced RNAPII transcriptional defect. Transition of E to A to B complex. Addition of U4/U5/U6 complex releases RNAPII from the pre-catalytic spliceosome. Decreased engagement of HTATSF1 with chromatin-bound SF3B1^K700E^ prevents transition to B complex preventing release of RNAPII from U1snRNP. (G) Metagene analysis of CDK9 ChIP-seq (IP/Input) as in WT (n=2) and SF3B1^K700E^ (n=2) cells. (H) Read-through and read-in transcription of 1000 most expressed genes in GRO-seq of WT and SF3B1^K700E^ cells as determined by ARTDeco.

### Effect of SF3B1^K700E^ on RNAPII kinetics is distinct from SF3B1 inhibition

Based on the nature of mutations (non-synonymous and never loss of function), and selection of novel 3’SSs, SF3B1 mutations are neomorphs and not loss of function variants^29^, ^13^. We sought to determine how SF3B1^K700E^ compares to SF3B1 chemical inhibition, in its effects on transcription. Acute inhibition of SF3B1 with PlaB largely results in RNAPII proximal-promoter (p-p)^5^, ^39^ pausing, a pattern distinct from the RNAPII redistribution in response to SF3B1^K700E^. SF3B1 inhibition also impairs recruitment of elongation complex p-TEFb to promoters, which then facilitates RNAPII from a paused state ^40^, ^41^. ChIP-seq for CDK9 (a component of the p-TEFb complex) expression showed increased CDK9 p-p occupancy in SF3B1^K700E^, suggesting that p-TEFb recruitment is not impaired. PlaB is acutely toxic to cells, and there is suggestion that at least some of PlaB-induced transcriptional alterations are indirect due to cellular stress response^42^. Transcriptional changes in response to cellular stresses include read-through transcription downstream of termination sites (also called Downstream of Genes or DoGs)^39^, ^43^. We utilized data from LRS, GRO-seq and TT-TL-seq to determine extent of readthrough transcription in SF3B1^K700E^ cells. We did not find evidence of increased readthrough past the TTS (Fig 3H, FigS3G-H) with any of the three approaches, arguing against such a perturbation of cleavage/termination in SF3B1^K700E^. Finally, pathway analysis of 4-day RNA-seq datasets from these cells do not show an upregulation of stress-related pathways in SF3B1^K700E^ (FigS3I). Collectively, these findings indicate that SF3B1^K700E^ -induced transcriptional changes are distinct from stress-related transcriptional disruption resulting from chemical inhibition of SF3B1.

### SF3B1^K700E^ induced transcription dysregulation is associated with R-loops, transcription replication conflicts (TRCs) and S-phase arrest

Isogenic SF3B1^K700E^ and U2AF1^SF34F^ cells were noted to have slower growth and fully arrest about 2 weeks after mutant allele expression (FigS4A), consistent with previous reports ^9, 44–46^. SF-mutant cells have been shown to have increased levels of R-loops and DNA damage response^44^, ^47^. Pathologic R-loops typically result from transcriptional dysregulation^48^. Increased R-loop levels were evident in in SF3B1^K700E^ by S9.6 monoclonal antibody immunofluorescence (IF) (Fig 4A). Of note, S9.6 is known to also bind to dsRNA and ribosomal RNA^49^, especially in nucleoli, which can confound R-loop staining. We used nucleolin staining to identify areas of nucleolar signal and subtracted those regions from the nuclei as previously described^50^ (Fig 4A). S9.6 dot-blots on chromatin preps corroborated our microscopy results (FigS4B). Unresolved R-loops pose a major obstacle to replication fork (RF) progression, by colliding with the DNA replisome machinery, leading to TRCs^48^. To determine if transcription and replication machineries are in close proximity, we performed in-situ proximity ligation assay (PLA)^51^ using antibodies against RNAPII and PCNA (a protein component of replication complex). As shown in Fig 4B, SF3B1^K700E^ nuclei had significantly higher foci of co-localized transcription and replication, characteristic of TRCs. Further characterization revealed SF3B1^K700E^ to induce DNA damage, as determined by neutral comet single cell gel electrophoresis assays (FigS4C) and γH2AX Western blotting (FigS4D). Increased replication stress in SF3B1^K700E^ was additionally confirmed using more specific markers, including IF staining for (1) increased ssDNA (BrdU labeling followed by anti-BrdU antibody^52^ IF) (FigS4E), and (2) phosphorylated RPA2 ser33 (pRPA)^53^ (FigS4F). BrdU and pRPA signals, as well as S9.6 staining, were specifically increased in those cells with increased γH2AX staining (FigS4E-G), indicating that R-loop related DDR is associated with increased replication stress. To better define mechanisms by which SF3B1^K700E^ perturbs R-loop homeostasis, we depleted representative factors from disparate pathways regulating R-loop homeostasis with siRNA: UAP56 (prevents nascent RNA back-hybridization^54^), Senataxin (resolves R-loops by promoting transcription termination^55^), and FANCD2 (a DDR factor^56^)^48^ (FigS4H). As anticipated, depletion of each of these factors increased both TRCs and DNA breaks (FigS4I) in the WT cells. In SF3B1^K700E^ cells, depletion of Senataxin alone (but not UAP56 or FANCD2) resulted in a further increase of both TRCs as well as DNA breaks (Fig 4C), compared to control siRNA. Furthermore, overexpressing bacterial RNaseH1 (a R-loop resolving enzyme) in SF3B1^K700E^ (FigS4J), improved cell growth, decreased comet tail moments, γH2AX foci and R-loops, confirming R-loops to be a major contributor to observed DDR (Fig 4A-B,4D; FigS4B-D).

Flow cytometry for DNA content was performed at 4 and 8 days after induction of ^K700E^ locus expression to determine phase of cell cycle arrest. As shown in Fig 4D (and FigS4K), distributions were similar on day 4 (when mutant allele is fully expressing, Fig 1A), but SF3B1^K700E^ accumulated at S-phase at day 8, corresponding to slowing cell growth. Importantly this reduction in G_1_ and increase in S-phase were abrogated by RNaseH1 overexpression (RNaseH1+OE). Co-staining for γH2AX, cleaved caspase 3, and 5-ethynyl-2′-deoxyuridine (EdU) also confirmed that DDR and apoptosis was limited to S-phase arrested cells (FigS4L). S-phase arrest in the absence of toxic or metabolic perturbation suggests replication stress wherein unstable replication forks cannot be resolved^57^, ^58^. Since R-loops arise at sites of transcription^48^, ^59^, and splicing is co-transcriptional, we reasoned that SF3B1^K700E^ spliceosomes are linked to TRCs and DDR through transcriptional dysregulation. To confirm that SF3B1^K700E^ induced DDR requires the presence of active transcription bubbles, G1/S synchronized cells at were released into the S-phase with or without triptolide (a RNAPII inhibitor^60^). The effectiveness of triptolide in inhibiting transcription was confirmed using through 5-ethynyl-2′-uridine (EU) incorporation (FigS4M). DNA damage signaling was assayed at the single cell level, using γH2AX IF and TRCs by PLA, at 6 hours post-release (FigS4M). Triptolide treated cells had significantly less DDR and TRC, confirming the role of transcription in SF3B1^K700E^-associated DNA damage. We finally determined DNA replication kinetics with using two approaches. First, DNA replication fork analysis of combed DNA fibers (using sequential double pulse labeling with two thymidine analogues (IdU and CldU))^61^, showed higher de novo origins of replication (FigS4N), reduced RF velocity (Fig 4E) and reduced DNA fiber track length (FigS4N) in SF3B1^K700E^ versus the SF3B1^WT^. These effects were partially reversed by RNAseH1-OE. Second, incorporation of nucleoside analogue 5EdU (a measure of newly synthesized DNA)^62^ was determined to be lower in SF3B1^K700E^ cells (Fig 4F). This decrease was also partially normalized by RNaseH1-OE. Together, these results indicate that SF3B1^K700E^-associated R-loops impair replication fork progression causing replication stress.

### R-loop accumulate primarily in gene bodies upon SF3B1^K700E^ expression

Reduced rate of transcription elongation rate leads to gene body R-loop accumulation and genome instability^63^, ^64^. Intensity of RNAPII arrest-associated R-loops are further modulated by the orientation of their accompanying TRCs^65^. To map R-loop distribution genome-wide, we immunoprecipitated and sequenced chromatin with S9.6 antibody (DRIP-seq)^66^(FigS4O, P). To avoid being confounded by cell-cycle phases that can affect transcription and R-loop formation. DRIP-seq was performed at 4 days of K700E locus induction when cell-cycle distribution was comparable in SF3B1^K700E^ and WT cells (Fig 4D). DRIP-seq data revealed 4576 peaks more enriched in SF3B1^K700E^, a ∼350% increase compared to SF3B1^WT^ (FigS4Q-R). Gene-metaplot showed progressively increasing signal intensity in gene bodies peaking at termination sites in SF3B1^K700E^ (Fig 4G). Relative enrichment of R-loops at TTSs is a feature of replication stress arising from “head-on” TRCs (HO-TRCs)^65^, ^67^, corroborating the increased TRCs and impaired RF progression. Since replisome movement usually orients in the same direction as transcription, HO-TRCs are particularly enriched in TTSs of gene neighbor pairs with convergently oriented gene transcription (convergent gene pairs) (FigS4S). Such convergent gene pairs are thus distinct from gene pairs which may be proximal but transcribe in the same or divergent directions (non-convergent gene pairs, FigS4S). Accordingly, we classified genes as convergent or non-convergent groups based on orientation of transcription between neighboring genes and constructed separate DRIP-seq metagene plots. As shown in Fig 4H, gene-pairs with convergent transcription had markedly increased enrichment of R-loop signals at TTSs in SF3B1^K700E^ implying the occurrence of HO-TRCs further augmenting the R-loop levels^65^ at these HO-collision hotspots. Furthermore, differential DRIP-signal increase at the TTS regions of converging gene pairs was dependent on distance between TTSs (FigS4U). By contrast, non-convergent genes R-loop enrichment at TTS enrichment in SF3B1^K700E^ was much less pronounced (FigS4T). In an alternative analysis, we intersected DRIP-seq peaks with previously reported genome-wide RF directionality data (Okazaki fragment sequencing or OK-seq) from normal K562 cells^68^ (FigS4V). SF3B1^K700E^-specific genic R-loop peaks were checked for enrichment for intersecting RFs– either HO or co-directional (CD). The absolute number of SF3B1^K700E^ DRIP-seq peaks (Fig 4I) and mean DRIP-seq signal (Fig 4J) were both higher in HO compared to CD conflict zones. The R-loop associated ssDNA accumulation at the stalled RFs likely serve as the substrates for robust ATR-Chk1 DDR pathway activation (FigS4W)^44^, ^47^ in SF mutations). Taken together, the DRIP-seq genome-wide distribution is consistent with a gene body elongation defect in SF3B1^K700E^, with R-loops particularly enriched at convergent gene-pair terminal sites correlating with HO conflict zones.

### SF3B1^K700E^ triggers changes to histone marks and chromatin accessibility

R-loops, chromatin organization and transcription are intricately linked.^69^, ^70^ Widespread epigenetic dysregulation is observed even in SF mutant myeloid disorders^71^, ^72^, and epigenetic therapies are effective in SF-mutant MDS/AML^71^. We therefore determined changes to chromatin states effected by SF3B1^K700E^. We used CUT&RUN sequencing^73^ to map well defined histone marks associated with functional genomic states: H3K4me3 (promoter transcriptional activity), H3K27ac (active promoters and enhancers), H3K27me3 (repressive chromatin), and H3K4me1 (distal regulatory elements including poised enhancers) (FigS5A). Among these, H3K4me3 was prominently reduced at promoters in SF3B1^K700E^ (Fig 5A-C) H3K4me3 marks are now known to be dependent on RNAPII transcriptional activity (specifically p-p release) and efficient splicing at the first exon-intron boundary^74^. Reduced p-p H3K4me3 marks in SF3B1^K700E^ (FigS5B) may therefore arise from defective transcription or splicing at this region. Consistent with this, changes in H3K4me3 signal closely tracked RNAPII ChIP-seq changes upon SF3B1^K700E^ expression (Fig 5E), underscoring the role of RNAPII density in H3K4me3 distribution^75^, ^76^. Among promoter regions, bivalent or “poised” promoters (enriched for both the active H3K4me3 and repressive H3K27me3 marks) showed decreased H3K4me3 and correspondingly increased H3K27me3 marks in SF3B1^K700E^ (Fig 5D). Such a change favors a closed chromatin configuration in these regions^77^. Of note, H3K27ac promoter signal intensities did not differ between SF3B1^WT^ and SF3B1^WT^(FigS5C).

**Figure 4:**
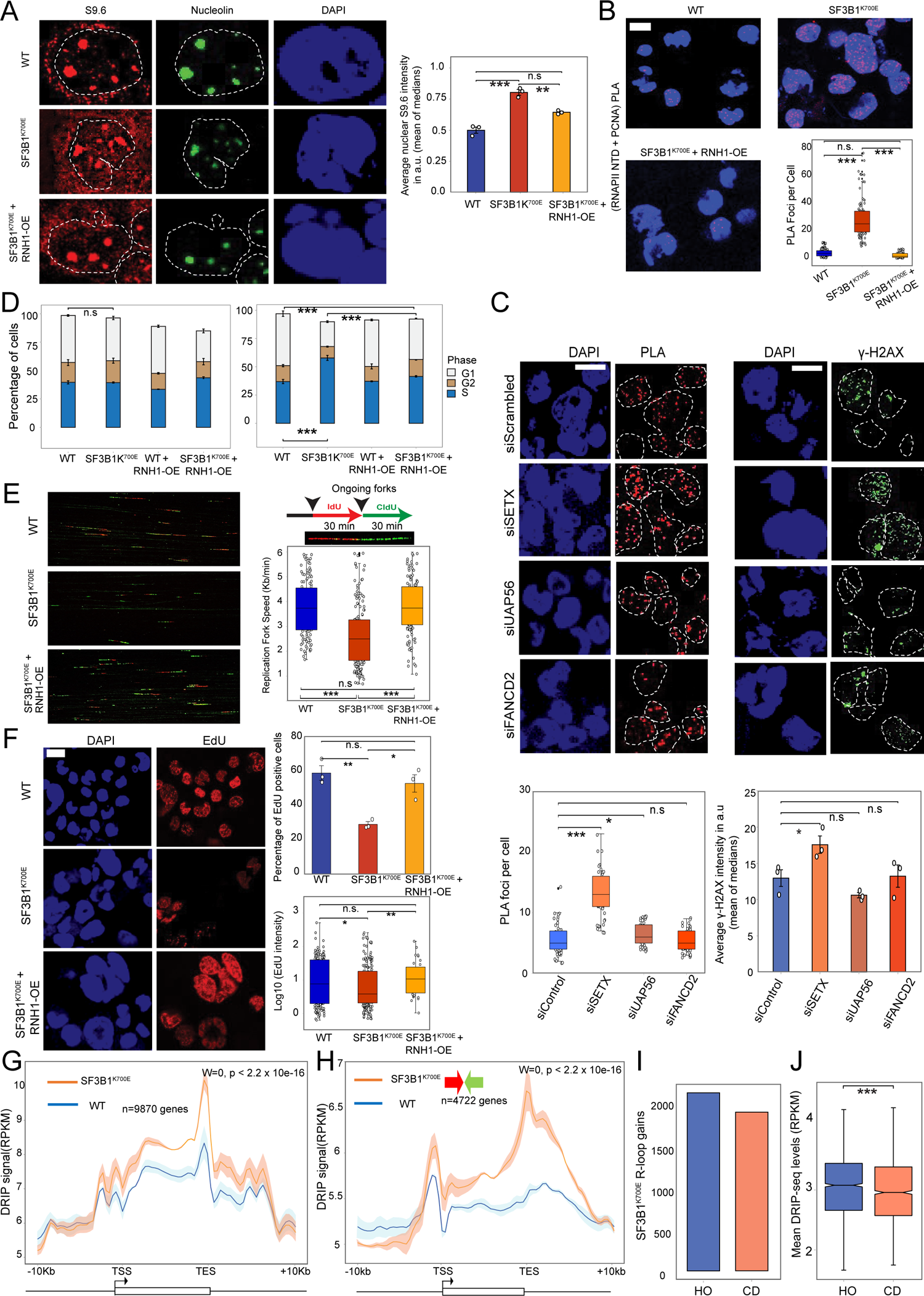
SF3B1^K700E^-induced changes to R-loops, DNA and replication dynamics. (A) Representative images and quantification of nuclear S9.6 intensity in WT, SF3B1^K700E^, and SF3B1^K700E^+RNH1OE (overexpression) cells at 5 days post induction. *n*=3 experiments. (two-tailed, Student’s *t*-test; \*\*\**P*=0.0009, \**P*=0.016, ^ns^*P* >0.05). Scale-bar, 2 μm. a.u: arbitrary units. (B) Representative images and quantification of RNAPII NTD + PCNA PLA foci in cells treated as in **A**. Each dot representing a PLA focus) (Mann-Whitney-U, two-tailed). \*\*\**P*<0.0001, ^ns^*P*>0.05. (C) Epistatic study between SF3B1^K700E^ and known R-loop preventing factors. Left Panel: Quantification of RNAPII N-terminal domain + PCNA PLA in SF3B1^K700E^ cells, transfected with scrambled siRNA (siC), and either siRNA against Senataxin (siSETX), FANCD2 (siFANCD2), or UAP56 (siUAP56). n=3 experiments. Scale-bar, 17 μm. (Mann-Whitney-U, one-tailed, \**P*=0.03, ^n.s^*P*>0.05). Right panel: Quantification of average gamma-H2AX nuclear intensity in SFB1^K700E^ cells, treated as in Left Panel. *n*=3 experiments. (one-tailed, Student’s *t*-test; \**P*=0.025, ^ns^*P*>0.05). Scale-bar, 17 μm. (D) Quantification of cell phase determined by flow cytometry for DNA content in WT, WT+RNH1-OE, SF3B1^K700E^, SF3B1^K700E^+RNH1-OE cells at 4 days and 8 days following doxycycline induction. Watson pragmatic fitting algorithm was used for cell cycle analysis. *n*=3 experiments. (\*\*\**P*<0.0001, ^ns^*P*>0.05; ANOVA followed by Sidak’s test). (E) Representative images and quantification of fork velocity performed in cells as in A, at 6 days post-doxycycline induction. (\*\*\**P*<0.0001, ^ns^*P*>0.05; one-tailed Mann-Whitney-U). (F) Representative images and quantification of percentage of cells incorporating EdU (Upper panel) and nuclear EdU intensity (lower panel) in cells, as in E. Scale bar, 17 μm. Upper panel: (*n*=3 experiments; two-tailed, Student’s *t*-test; \*\**P*=0.0028, \**P*=0.019, ^ns^*P*>0.05). Bottom panel: (*n*=3 experiments). (one-tailed Mann– Whitney-U. \*\**P*=0.0037, \**P*=0.039, ^ns^*P* >0.05). (G) Metaplot of the distribution of S9.6 signals (DRIP-seq mean coverage) along gene bodies and flanking regions (±10 kb) of all genes expressed in K562 in WT (n=2) and SF3B1^K700E^ (n=2) K562 cells. Differences in signal intensity at TTS±2 kb calculated with Wilcoxon rank-sum test with continuity correction. (H) Metaplot of subset of convergent gene/gene pairs as in (G). (I) Absolute numbers of head-on (HO; blue bar) vs co-directional (HO; orange bar) transcription-replication conflict zones intersecting with the SF3B1^K700E^-specific R-loop peaks. (J) Average DRIP-seq coverage at SF3B1^K700E^-specific R-loop peaks reached by the replication fork in CD or HO orientation during DNA replication. (Mann-Whitney-U, two-tailed, \*\*\**P*<0.0001).

To determine changes to chromatin accessibility and nucleosome organization, we performed ATAC-seq. An overall reduction in chromatin accessibility was evident in SF3B1^K700E^ compared to SF3B1^WT^ (Fig 5F). Of the ATAC peaks downregulated in SF3B1^K700E^, 44% were at promoters and 13% at enhancers. R-loops may increase or decrease chromatin accessibility, depending on the context^78^. Given the preponderance of excess R-loops at gene bodies, we posited that changes to chromatin accessibility and R-loops may be co-localized. Colocalization analysis^79^ of downregulated ATAC-seq peaks and differential DRIP-seq peaks in SF3B1^K700E^ showed enrichment of upregulated promoter and gene body DRIP-seq at closed chromatin regions (Fig 5G, FigS5D). This suggests that excess R-loops are closely related to reduced chromatin accessibility in SF3B1^K700E^. We next implemented NucleoATAC^80^, an algorithm to infer nucleosome occupancy from ATAC-seq datasets to determine nucleosome occupancy at TSS. TSS of actively transcribing genes are devoid of nucleosomes; thus, increased nucleosome occupancy denotes transcriptional repression^81^, ^82^. As shown in Fig 5H, SF3B1^K700E^ had increased nucleosome occupancy at TSS, further confirming transcriptional repression accompanying RNAPII elongation defects. GRO-seq RNAPII density at promoter and enhancers is a surrogate measure for their ability to activate transcription of their target genes^83^. We found loss of transcribing RNAPII activity extending from pausing sites (highest GRO-seq signal within the +20 to +300 bp window downstream of TSS) to downstream regions for the sense transcription, at both promoters (upstream region of TSS) and putative enhancers (distal H3K27ac peaks) (Fig 5I, FigS5E-F). Taken together, our data shows changes to histone marks and chromatin accessibility in SF3B1^K700E^, in patterns consistent with reduced RNAPII transcription elongation.

**Figure 5:**
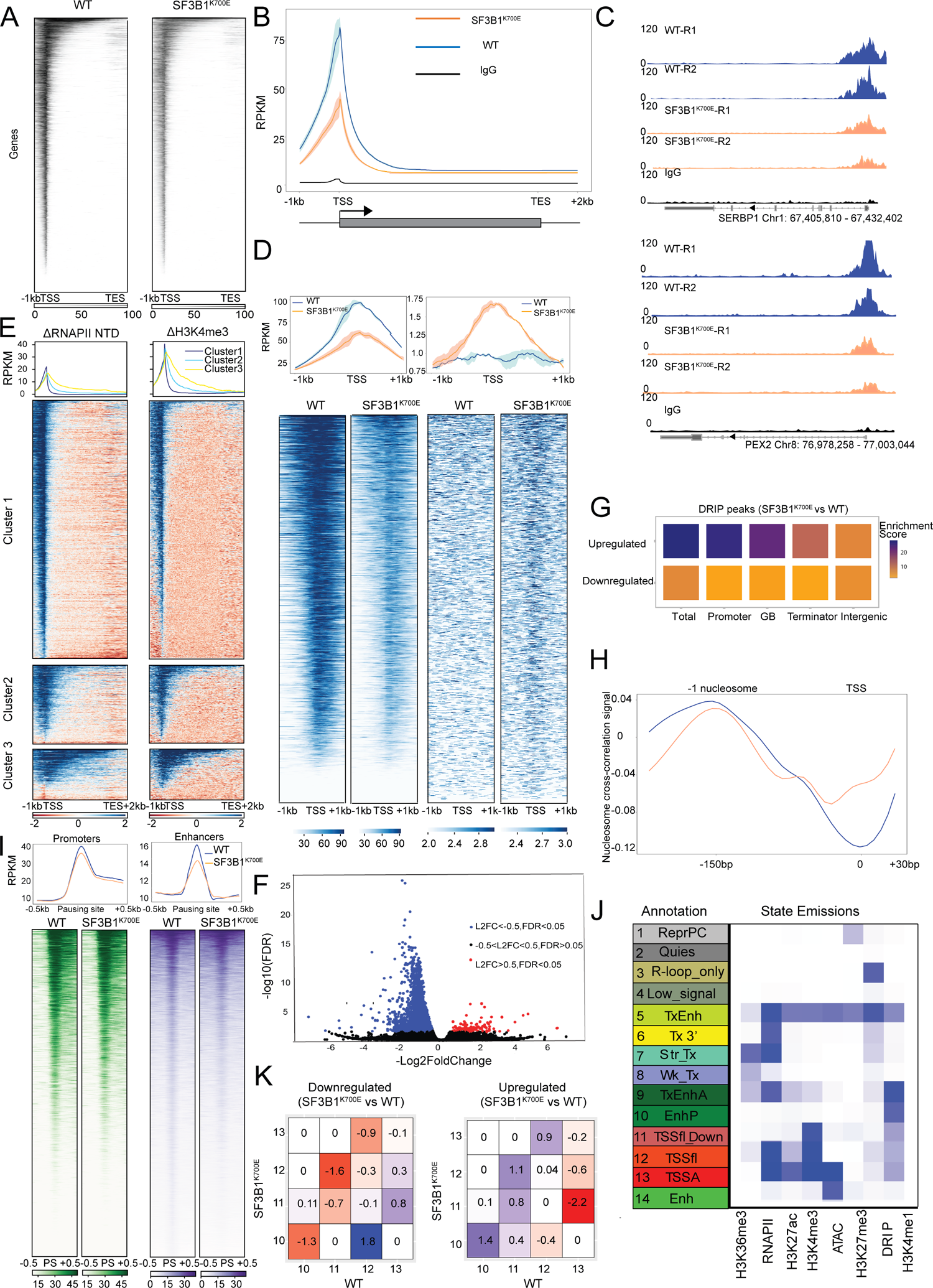
Transcription-coupled changes in chromatin accessibility and histone marks due to SF3B1^K700E^. (A) Heatmap plots depict H3K4me3 signal in WT (left) and SF3B1^K700E^ (right) K562 cells, at 4 days post doxycycline induction. Genes ranked by decreasing H3K4me3 occupancy. (B) Metagene plot showing the normalized H3K4me3 CutnRun levels over 6694 expressed genes in WT (n=2) and SF3B1^K700E^ (n=2). IgG included as control (in black). (C) Representative genome-browser track examples of H3K4me3 CutnRun-seq in WT versus SF3B1^K700E^ for the indicated genes. (D) Metagene and heatmap plots depict H3K4me3 (left) and H3K27me3 (right) signals in WT and SF3B1^K700E^ K562 cells at bivalently marked gene-promoters (n=2168 genes). Signals mapped from −1kb to +1kb of TSS. (E) Heatmaps depict changes in RNAPIINTD ChIP and H3K4me3 CutnRun average signal enrichments (WT–SF3B1^K700E^) in K562 cells. Genes broken down by groups, based on Fig 1F. (F) Volcano plot of ATAC-seq differential analysis between SFB1^K700E^ versus WT (n=2, each). FDR-false discovery rate. (G) RegioneR-based colocalization analysis of selectively downregulated ATAC peaks in SF3B1^K700E^ and DRIP-seq-based differential R-loop regions (SF3B1^K700E^ vs WT). (H) Metaplot of WT and SF3B1^K700E^ ATAC-seq signal profiling changes in the nucleosome occupancy among high expressing genes (FPKM>1000, *n* = 274 from dataset of 6694 genes). (I) Left: Metaplots and heatmaps of RPKM-normalized GRO-seq signal for sense transcription at promoters (left) and putative enhancers (right), ranked by decreasing occupancy. (J) Fourteen chromatin states learned jointly for WT and SF3B1^K700E^ cells by ChromHMM. The heatmap shows the state emissions learned on the basis of combinatorial analysis, where each column corresponds to a different epigenetic signature and each row corresponds to a different chromatin state. Candidate state annotations for each state are listed. Blue shading indicates intensity. Putative state annotations: 1.ReprPC-repressed polycomb, 2.Quies-quiescent, 3.Rloop_only: R-loop-only state, 4.Low_signal: low-signal state, 5.TxEnh-transcribing enhancers, 6.Tx 3’-transcribing gene 3’end, 7.Str_Tx-Strongly transcribing genic region, 8.Wk_Tx-Weakly transcribing genic region, 9.TxEnhA-active transcribing enhancers, 10.EnhP-poised enhancers, 11.TSSfl_Down-Region flanking downstream of TSS, 12. TSSfl-Region flanking TSS, 13.TSSA-active promoters, 14.Enh-Enhancer. (K) Relative enrichment of chromatin state transitions at promoters of downregulated genes (left panel) or upregulated genes (right panel) in SF3B1^K700E^ (vs WT), for chromatin state transition pairs involving states 10, 11, 12, and 13. Red shows enrichment, whereas blue shows depletion. Log-fold change values are listed in each box. For further details, see extended supplemental figure 5 data.

We next applied a Hidden Markov Modeling (ChromHMM)^84^, defining a set of chromatin states based on the histone modifications, DRIP-seq, and ATAC-seq, to understand how chromatin landscape alterations correlate with RNA-seq gene expression changes in SF3B1^K700E^ cells. A final model containing 14 states, annotated by intersecting each with known genome annotation features (FigS5G-H), was adopted for downstream analysis (Fig 5J). Within each of these groupings, enrichment of specific genomic structures was as expected (FigS7G-H), supporting the biologic relevance of the assigned state annotations. Calculation of coverage changes in the genome-wide occupancies for each state, from SF3B1^K700E^ to WT, are shown in FigS5I. To specifically understand relationships between chromatin states and mRNA-seq gene expression, we calculated the relative enrichment of all possible chromatin state transitions at the promoters (+0.5 kb and −0.5 kb from TSS) of genes that were differentially expressed, with an expectation that genes downregulated in SF3B1^K700E^ cells would show a switch from the most ‘active’ promoter state (state 13) to ‘less active’ states (states 10 - 12) on their promoters and vice-versa. As highlighted in Fig 5K, (see also supplemental, Sourcedata_ChromHMM file for details), we observed a reciprocal pattern of chromatin state transition enrichments consistent with ‘active’ to ‘less active’ and ‘less active’ to ‘active’ in the SF3B1^K700E^ downregulated and upregulated genes, respectively. These data indicate the combinatorial effects of transcription kinetics, R-loop distribution and changes to chromatin landscape on the steady state transcriptome in SF3B1^K700E^ cells.

### Components of the Histone Deacetylase/H3K4me pathways modulate sensitivity to SF3B1^K700E^

Based on the above results, we speculated if modulating epigenetic regulators may improve RNAPII elongation defects and rescue cells from downstream effects including growth arrest. We hence performed a forward genetic screen with a short hairpin RNA (shRNA) library targeting epigenetic regulator and chromatin modifier protein classes (∼2500 shRNAs targeting 357 genes, experimental details in Fig 6A). We focused our analyses on those shRNAs that promoted survival of genes progressively in SF3B1^K700E^ (^“^survival candidates”) since knockdown of the candidate genes would improve cell survival and growth (Fig 6B; TableS1). The top 20 survival candidates were noted to be highly enriched in the histone deacetylase (HDAC) pathway by GO analysis (FigS6B). Analysis for protein-protein interactions using the StringDB package^85^ showed a similar enrichment of Sin3/HDAC complex (Fig 6C). Specifically, the top hits (HDAC2, ING2, and PHF21A (or BHC80)) are part of the extended LSD1-Sin3-HDAC transcriptional corepressor complex which negatively modulates H3K4me activity^86^, ^87^, ^88^. Additionally, WDR5, a key writer component of H3K4me3 methyltransferase complex^89^, was depleted in the screen (i.e., WDR5 loss led to accelerated cell death in SF3B1^K700E^ cells) (TableS2).

**Figure 6:**
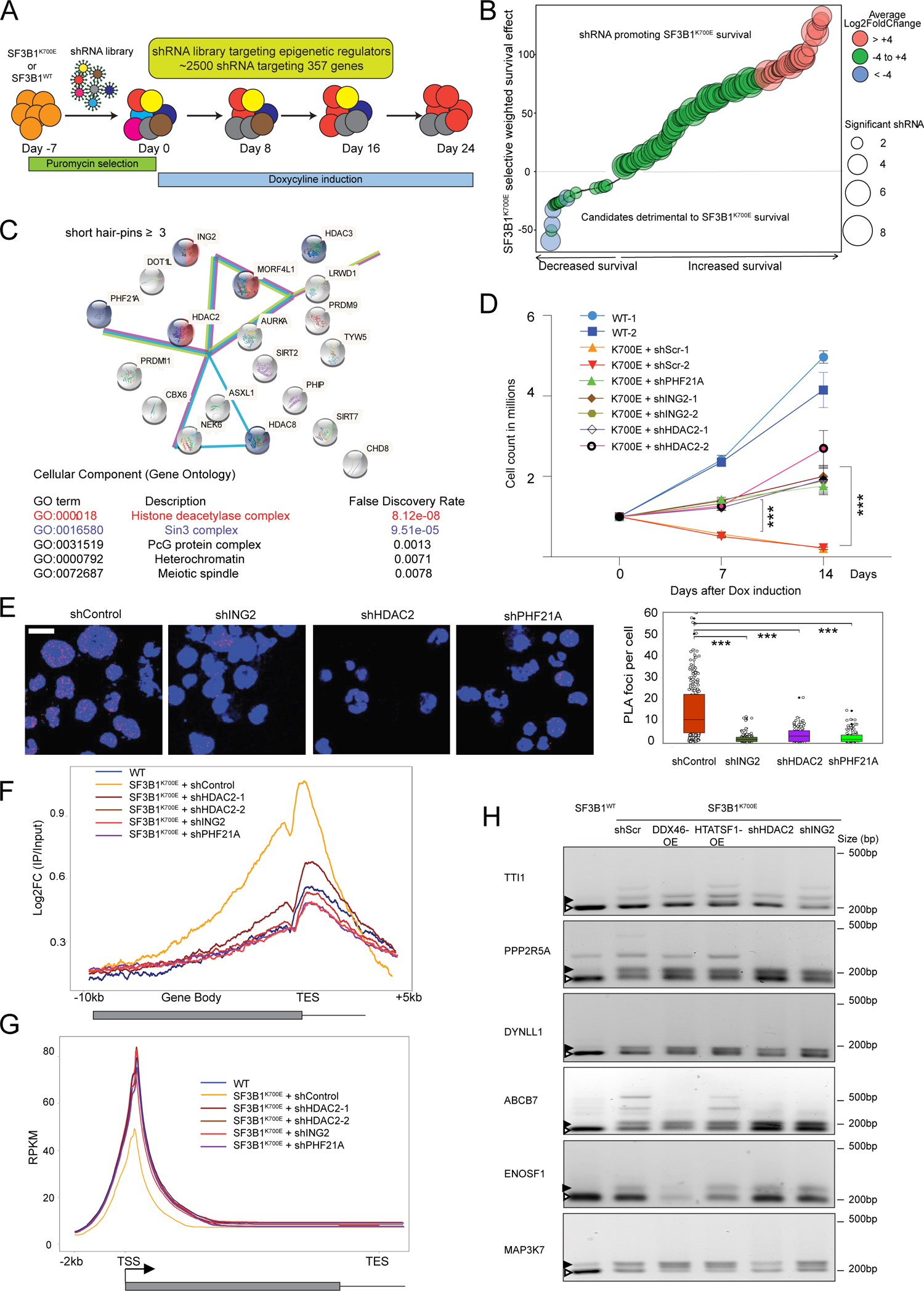
An epigenetic shRNA rescue screen identifying key pathways functionally important to SF3B1^K700E^-related cell survival. (A) shRNA screen for epigenetic/chromatin modifier genes that modulate sensitivity to SF3B1^K700E^. (B) Waterfall bubble plot shows combined SF3B1^K700E^-selective survival effect of each gene, calculated as a weighted effect of knockdown by multiple shRNAs. (C) Sin3 (marked as violet) and HDAC (marked as red) complex components are enriched among genes that promote cell survival in SF3B1^K700E^. StringDB functional enrichment analysis was performed on the top 20 survival candidates (Log2 fold enrichment>4 and shRNAs=>3 against a particular target). Top 5 functionally enriched cellular component pathways with FDR<0.001 are shown. (D) WT, SF3B1^K700E^+shScrambled, SF3B1^K700E^+shING2, SF3B1^K700E^+shHDAC2, SF3B1^K700E^+shPHF21A cells were induced with doxycycline for 4 days (Day 0 at 96 hours post-induction time point), and cell growth plotted over a course of 14 days. (*n*=3 experiments). (\*\*\**P*<0.0001_;_ two-tailed *t* test). (E) Representative images and quantification of RNAPII NTD+PCNA PLA foci in SF3B1^K700E^+shScrambled (shScr), SF3B1^K700E^+shHDAC2-1, SF3B1^K700E^+shING2-1, SF3B1^K700E^+shPHF21A at 4 days post induction. (Mann-Whitney-U, two-tailed). ***P<0.0001. Scale-bar: 15 μm. (F) Metagene plot analysis showing the log_2_-fold change (L2FC) of anti-Ser2PRNAPII ChIP-seq (IP/Input) across the annotated gene transcripts genes (n = 6694) in cells as in D, at 4 days post-doxycycline induction. (G) Metagene plot analysis showing RPKM normalized H3K4me3 CutnRun levels over 6694 expressed genes in cells as in F. (H) Total RNA from the indicated cell lines was extracted at 5 days post- doxycycline induction, followed by cDNA RT-PCR of the cryptic 3′SSs (black arrowheads) and canonical 3’SSs (blue arrowheads) of six select genes.

Secondary validation experiments with two shRNA each for HDAC2, ING2 and PHF21A confirmed improved cell survival (Fig 6D), upon knockdown of these genes (FigS6C). Next, we determined that the improved cell survival was associated with reductions in R-loops (FigS6D), DDR (FigS6E-F), TRCs (Fig 6E), and normalization of cell-cycle phase (FigS6G) and DNA replication dynamics (FigS6H-I). Furthermore, a reduction in Ser2PRNAPII gene body densities (Fig 6F) accompanied by a significant increase in promoter H3K4m3 (Fig 6G) was observed in the knockdown SF3B1^K700E^ cells, confirming that loss of these modulators affected H3K4me3 mark deposition in the promoter region either directly or through improvement in RNAPII elongation kinetics. Taken together, our results suggest that transcription changes and downstream effects induced by SF3B1^K700E^ can be mitigated by modulating the H3K4me3 functional pathway^90^.

RNAPII kinetics can potentially influence 3’splice site selection^7^. To determine how RNAPII elongation may influence selection of cryptic 3’SSs in SF3B1^K700E^, we first identified cryptic 3’SSs (from both our K562 isogenic cells and analysis of patient samples^14^) and determined RNAPII distribution in those introns containing such cryptic 3’SSs. RNAPII distribution changes in this subset of introns was no different from the remaining unselected introns (FigS6J-K). We then checked if reversing RNAPII kinetics (through over-expression of HTATSF1, or knockdown of HDAC2/ING2) changed cryptic 3’SS usage by RT PCR. As shown in Fig 6H, change in RNAPII elongation did not appear to have an effect on selection of cryptic 3’SS. These findings suggest that altered transcription and selection of cryptic 3’SS in SF3B1^K700E^ is likely driven by independent mechanisms.

### WDR5 inhibition reduces colony formation in SF3B1-mutant MDS

We analyzed CD34+ cells isolated from up to seven SF3B1-mutant MDS patient samples (and 5 CD34+ controls from normal age-matched donors) to confirm key findings from the isogenic cell line system (Fig 7A & FigS7A). MDS patient-derived CD34+ cells showed significantly higher TRCs (FigS7B) and DDR (FigS7C) by gamma-H2AX and PLA microscopy, respectively. Given that ChIP-seq is difficult to perform with low cell numbers, we optimized CUT&RUN for Ser2P RNAPII to map RNAPII distribution. Metagene analyses of Ser2P RNAPII occupancies revealed a redistribution of RNAPII with reduced promoter/TSS density and increased gene body levels (Fig 7B-C) in MDS samples, similar to isogenic cells. CUT&RUN for H3K4me3 demonstrated a significant reduction in promoter H3K4me3 densities in SF3B1-mutant MDS (Fig 7D-E). Nucleosome mapping ATAC-seq datasets from patient samples also showed changes similar to those in isogenic cell systems (reduction in nucleosome occupancy at the −1 and +1 positions accompanied by higher occupancy at the nucleosome-depleted region) (Fig 7F). Our patient sample data thus confirms changes to transcription kinetics and chromatin organization in primary patient samples as predicted by isogenic SF3B1^K700E^ cells.

**Figure 7:**
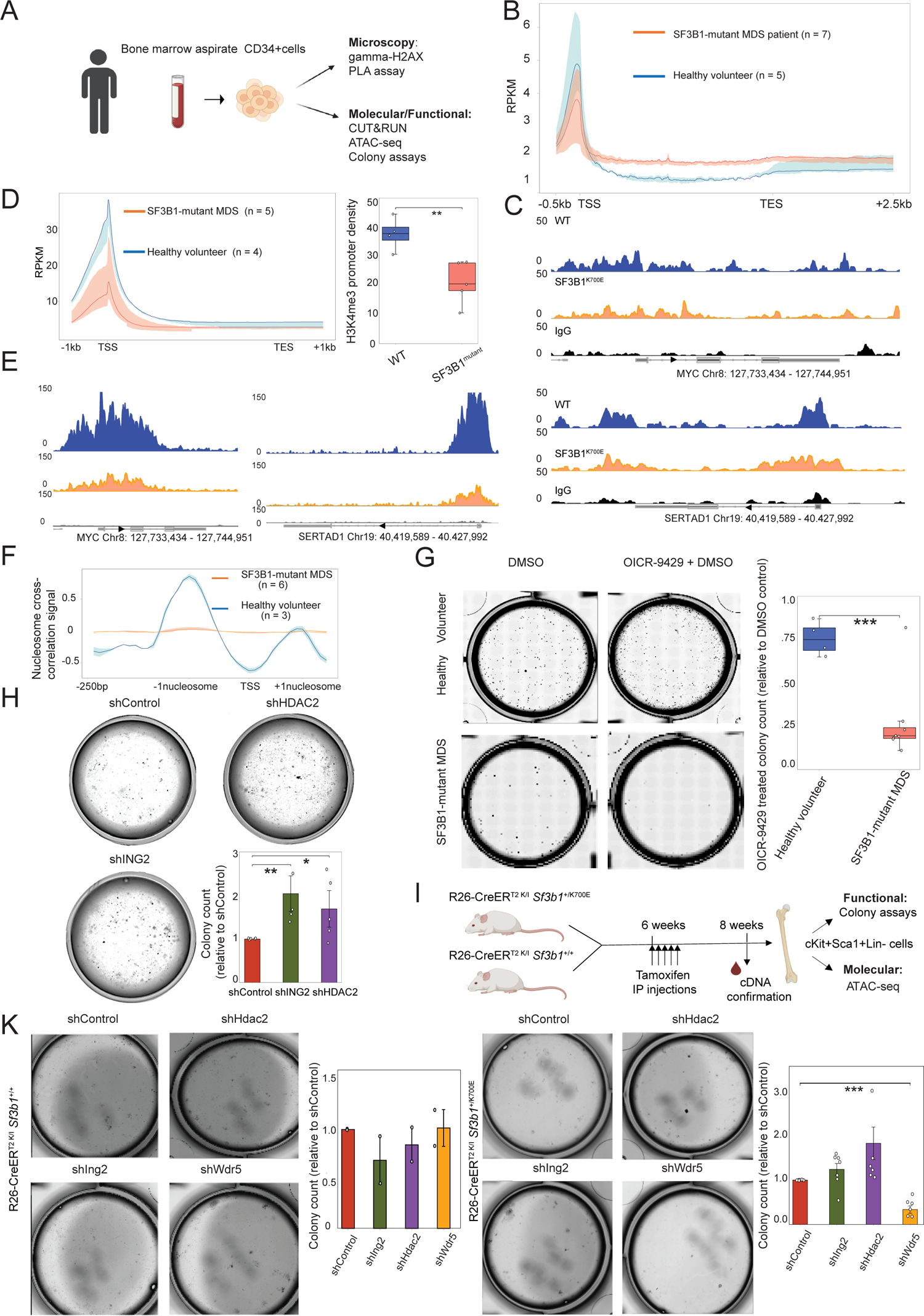
Targeting H3K4me pathway in SF3B1-mutant MDS. (A) Workflow for isolation/characterization of SF3B1-mutant MDS patient marrow aspirate-derived CD34+ cells. (B) Metagene plot analysis showing RPKM-normalized CUT&RUN levels of Ser2PRNAPII over 6694 expressed genes in WT and SF3B1-mutant CD34+ cells. Solid lines represent averaged signal, and upper and lower borders of the shaded area represent the highest and lowest densities among the replicates, combined (n=5, WT; n=7, SF3B1-mutant MDS). (C) Representative genome browser track examples of Ser2PRNAPII CUT&RUN in CD34+ cells: WT versus SF3B1^K700E^, for the indicated genes. Corresponding IgG tracks are in black. (D) Left: Metagene plot showing RPKM-normalized CUT&RUN levels of H3K4me3 over 6694 expressed genes in WT and SF3B1-mutant MDS CD34+ cells. (n=4, WT; n=5, SF3B1-mutant MDS). Right: Box-whisker plots of H3K4me3 promoter density in WT and SF3B1-mutant (student t-test, two-tailed, \*\**P=*0.0072). (E) Representative genome browser track examples of H3K4me3 CUT&RUN in CD34+ cells: WT versus SF3B1-mutant, for the indicated genes. (F) Metaplot of WT and SF3B1-mutant ATAC-seq signal profiling changes in the nucleosome occupancy across 6694 expressing genes in WT (n=3) and SF3B1-mutant (n=6) CD34+ cells. (G) Methylcellulose colony-forming (MCF) unit assay of mononuclear cells isolated from healthy volunteer and MDS patient-derived peripheral blood/bone marrow aspirates. Isolated cells were treated with either 0.05% DMSO or 5μM OICR-9429 (in 0.05% DMSO) on day 0 after isolation and incubated in methylcellulose medium for 14 days before colony counting and imaging. Colony formation analysis in WT (n=4) and SF3B1-mutant (n=8 experiments) treated with OICR-9429 (relative to DMSO controls). \*\*\**P*=0.0003 (two-tailed t-test). (H) MCF assay of mononuclear cells isolated from MDS patient derived peripheral blood/bone marrow aspirates. Representative images and colony count formation analysis in shScrambled, shHDAC2, shING2 (all relative to shScrambled). \**P_shHDAC2vsScrambled_=0.01, *P_shING2vsScrambled_*=0.05 (one-tailed t-test). (I) Experimental scheme for murine experiments. LSK population (cKit+Sca-1+Lin-cells) were isolated from marrow cells by FACS and divided for CFU or ATAC-seq. (J) Methylcellulose colony forming unit assay of LSK cells isolated from murine *Sf3b1*^+/+^ and *Sf3b1*^+/K700E^ bone marrow aspirates. Representative images and colony formation analysis in shScrambled, shHdac2, shIng2, shWdr5 (all relative to shScrambled) shown for *Sf3b1*^+/+^ (left) and *Sf3b1*^+/K700E^ (right^)^. \*\**P_shWdr5vsshScrambled_<0.01*(one-tailed t-test).

We next sought to determine effects of epigenetic targets identified by the shRNA screen on primary MDS patient samples. Based on the shRNA screen results (Fig 6) that showed increased cell death upon knockdown of WDR5, we used OICR-9429, a small molecule inhibitor with known anti-tumor effects^91^, ^92^ to perform methyl cellulose colony forming assays. As shown in Fig 7G, treatment with OICR-9429 at the previously established IC_50_ of 5μM^92^, ^91^ had a modest effect on colony growth in normal controls, but significantly decreased in colony growth of CD34+ hematopoietic progenitors of SF3B1-mutant MDS samples. Additionally, loss of both HDAC2 and ING2 knockdowns by shRNA significantly increased colony formation in these cells (more pronounced with shING2, Fig 7H, FigS7D). Sequencing of mutation alleles from colonies confirmed preservation of mutant allele upon growth, and not improvement of unmutated clones (FigS7G). Flow cytometry evaluation of colonies at 14 days show that both mature erythroid (CD71) and myeloid colonies (CD11b) arose upon expansion compared to control transfected with scrambled shRNA (FigS7E-F). Thus, knocking down ING2 or HDAC2 promoted lineage commitment and differentiation in primary SF3B1-mutant MDS samples.

We finally utilized a previously established conditional murine model of Sf3b1^K700E^, ^93^ to independently confirm its functional interplay with Sin3/HDAC pathway in primary cells. The Sf3b1^K700E^ mice were crossed with Tamoxifen-Cre mice, and double mutant mice (*R26-*CreER^T2 KI/+^ *Sf3b1*^+/K700E^) were induced by intraperitoneal injections with tamoxifen. LSK progenitor population (cKit+Sca-1+Lin-) were isolated from bone marrow (controls were *R26*-CreER^T2 KI/+^ *Sf3b1*^+/+^)) (Fig 7I, FigS7G) and transduced with lentiviruses expressing shRNAs targeting Hdac2, Ing2, and Wdr5, or scrambled control. The ability of cells to form colonies was assayed by standard methyl-cellulose CFU assays at 14 days of cultures. Consistent with observations from human patient sample data, we observed an increase in colony growth with loss of Hdac2 and Ing2 (Fig 7J, FigS7H). Conversely, Wdr5 knock-down inhibited colony growth selectively in the mutant cells (Fig 7J). We also performed ATAC-seq on LSK cells to determine changes to nucleosome occupancy. As shown in FigS7I, NucleoATAC analysis showed a similar pattern of nucleosome occupancy in mutant murine cells as was observed for both patient samples and isogenic cells. These findings in the murine model provide additional validation to the role of Sin3/HDAC/H3K4me3 pathway in SF3B1-mutant diseases, representing a targetable functional vulnerability.

## DISCUSSION

Studies investigating molecular mechanisms by which mutant SFs serve as oncoproteins have largely focused on changes in the isoform ratio of well-defined tumor suppressors or oncogenes through AS^94^, ^95^. We independently analyzed 172 MDS/AML patient samples with SF mutations and 45 wild-type controls, curated from three different studies^31^, ^33^, ^32^ to determine patterns of AS associated with mutational subtypes (FigS8). Consistent with previous reports, we found evidence of widespread AS events that clustered only loosely along the mutational subtypes. However, there were few AS events common across the different SF mutations that could explain the common MDS phenotype. This prompted the current study to identify AS-independent mechanisms. One such is the increased presence of R-loops and DDR observed across the various SF-mutation subtypes^44^, ^47^. For SRSF2, a reduced ability of the mutant protein to extract pTEFb from the inhibitory complex 7SK was proposed to underlie the impaired p-p release of RNAPII^44^. This interaction being specific to SRSF2 is not applicable to SF3B1 or U2AF1.

Our genome-wide analysis that linked RNAPII elongation defects in SF-mutations to increased R-loops, replication stress, and chromatin re-organization prompted us to ask what mechanisms underlie these transcription defects, and how these maybe therapeutically targeted:

### SF3B1 mutations are associated with distinct patterns of AS; hence how are splicing and RNAPII elongation defects related?

SF3B1 mutations are associated with selection of “cryptic” 3’SS, and also associated with a decrease in retained introns^29^, ^13^. Hence, our analysis focused on how RNAPII occupancy is related to the selection of cryptic 3’SSs and retained introns. We found that introns containing such cryptic 3’SSs did not differ from unselected introns in their RNAPII distribution. Additionally, correction of RNAPII elongation defects did not reverse cryptic 3’SS. Next, we studied CoSE, a metric to determine the efficiency of intron splicing in nascent RNA. If reduction of RI is related to improved co-transcriptional splicing efficiency, an increased CoSE is expected. SF3B1 mutations were associated with a reduction in CoSE, suggesting that the reduced RI does not arise from the improved efficiency of co-transcriptional splicing. Taken together, our data suggest that changes in alternative splicing and RNAPII transcription in response to SF3B1^K700E^are independent. While the use of cryptic 3’SS appears to be related to use of non-canonical BP (REF), the basis of reduced RI is unclear, but likely results from improved processing of nuclear-detained RI, which requires further study.

### What are the molecular mechanisms by which mutant SFs impact RNAPII elongation?

Recent cryo-EM studies have shed light on the organization of early spliceosome complexes^34^, ^35^. Accordingly, a critical step in the assembly of the pre-spliceosome A complex is the eviction of HTATSF1 bound to SF3B1 by DDX46^34^, ^35^. RNAPII is released from the spliceosome only upon the transition to the pre-B complex^2^. Our results show that protein-protein interactions of SF3B1^K700E^ with HTATSF1, U2AF1 and U2AF2 are impaired, and this results in reduced transition of early spliceosome complexes to the pre-catalytic B complex. Based on our study findings, we propose a model in which impaired interaction of SF3B1^K700E^ with components of the early spliceosome including HTATSF1^96^, ^97^ prevents transition from the E-to-A complex impairing the release of RNAPII from its bound configuration to U1 snRNP (Fig 3F). We speculate that this impaired release underlies the RNAPII elongation defect along gene bodies.

### Given the interplay between transcription and chromatin states, how do RNAPII elongation defects affect chromatin organization and vice versa?

Interdependence of transcription on chromatin organization has been well described^70^, ^98^, ^99^. Furthermore, we found that SF3B1^K700E^ associated transcription changes were accompanied by changes in histone marks (predominantly H3K4me3) and chromatin accessibility. The results of our shRNA library screen strongly implicate the Sin3/HDAC pathway in regulating RNAPII elongation and its downstream effects. These results also hold true in primary patient samples and a mouse model of Sf3b1^K700E^. Our results linking transcription to chromatin organization are significant for many reasons. First, MDS is broadly susceptible to epigenetic therapies regardless of the type of driver mutation. Our results suggest that even with SF-mutant disease without genetic alterations to epigenetic factors, mechanisms may converge upon epigenetic mechanisms. Second, these results may explain why SF mutations are largely exclusive to some epigenetic mutations (such as those in the PRC2 complex)^100, 101^. One possibility is that PRC2 and SF mutations may have similar effects on chromatin organization rendering dual mutant cells unviable. The role of 3D genome organization in clonal myeloid disorders has recently been explored^102^, and our results underscore the need for mechanistic studies that link oncogenic drivers with altered chromatic organization.

### Can the modulation of RNAPII elongation defects be exploited therapeutically?

WDR5 is a key “writer” in the COMPASS H3K4me3 methyltransferase complex, which was depleted in our shRNA rescue screen, suggesting a lethality due to its loss in SF3B1^K700E^ cells. Indeed, chemical inhibition of WDR5 resulted in profound inhibition of colony growth in patient samples as well as in mutant mouse cells without affecting wild-type controls. WDR5 inhibitors are currently explored for their anti-tumor effects broadly^103^, ^104^, ^91^, ^105^. Our results suggest they may be particularly effective in situations where RNAPII transcription elongation is disrupted.

In summary, our results define how RNAPII elongation is disrupted in response to oncogenic mutations in SF3B1 and U2AF1. Modulation of epigenetic regulators improves RNAPII transcription kinetics and point to potential therapeutic targets. An important inference from our model is that the RNAPII elongation defects arise from the defective assembly of early spliceosomes. Interestingly, most SFs mutated in disease, including SF3B1, U2AF1, SRSF2, U1 snRNA, have established roles in early spliceosome assembly^106^, suggesting a possible shared defective RNAPII transcriptional response to these mutations. Exploring RNAPII kinetics and early spliceosome assembly in these additional mutational contexts will be informative for defining potential therapeutic targets similar to the Sin3/HDAC pathway in *SF3B1* mutations.

### Limitations of the study

This study primarily utilized an ENCODE tier 1 cell line, K562, for genome-wide studies and screens. We acknowledge that specific genes or pathways relevant to oncogenesis cannot be inferred from transformed cell lines. However, the fundamental aspects of transcriptional regulation are likely preserved across cell systems. Critical aspects of our work were confirmed in human samples and mouse models, an approach used in many previous studies. Although our sequencing data were collected at the earliest optimal time point of 4 days following doxycycline induction, we cannot entirely rule out indirect or secondary effects. LRS studies are currently limited by the low total read depth on the PacBio platform. The currently available murine models of *Sf3b1*^K700E^, ^107^, ^93^ do not confer clonal advantage in repopulation assays, making *in vivo* studies of clonal expansion to study the effect of modulating specific proteins (such as WDR5) challenging. Finally, confirmation for the working model we propose may require additional approaches may benefit from studies utilizing single molecule microscopy or molecular structure.

## MATERIALS AND METHODS

### Antibodies

Antibodies used in the study are listed in Table S8

### Cell lines

All parent cell lines were obtained from ATCC unless otherwise stated, authenticated by ATCC STR profiling, and tested for mycoplasma contamination at regular intervals using Mycoplasma PCR detection kit (Boca Scientific). K562 were grown in RPMI1640 medium supplemented by 10% fetal bovine serum (FBS) and 1% penicillin-streptomycin. HEK293T cells were grown in DMEM supplemented by 10% FBS and 1% penicillin-streptomycin. *Drosophila melanogaster* S2 (Schneider-2) cells (used for TT-TL seq and ChIP-seq spike-ins) were cultured in Schneider’s modified *Drosophila* medium (Lonza) supplemented with 10% FBS at 27°C without CO_2_. Doxycycline induction was performed using Doxycycline (Sigma) at 1 ug/ml concentration respectively. Puromycin, blasticidin, hygromycin, and neomycin antibiotic resistance selections and growth were performed at 1 ug/mL, 10 ug/mL, 250 ug/mL, and 200 ug/mL concentrations, respectively. Isogenic SF3B1^K700(E/K)^ and U2AF1^S34(F/S)^ endogenous knock-in K562 cells were generated using a combinatory AAV intron trap-CRISPR approach (sgRNA/CRISPR mediated DSB and AAV-intron trap-vector donor template) as previously described^20^. In case of SF3B1^K700E^, the mutation is ‘A’ to ‘G’ which results in addition of Glutamic acid in place of Lysine at 700 amino acid position in *SF3B1*. In case of U2AF1^S34F^, the mutation is ‘C’ to ‘T’ which results in addition of Phenylalanine in place of Serine at 34 amino acid position in *U2AF1*. Wild-type K562 cells are subject to the exact same genome manipulation except that the wild-type cells have a synonymous change (A-to-A in SF3B1 and S-to-S in U2AF1).

### siRNA and shRNA mediated knock-down

For single siRNA knockdown K562, cells were electroporated with 50 nM siRNA DharmaFECT 1 (Dharmacon) per manufacturer’s instructions (Nucleofector V kit, T-106 program). Pooled siRNAs (ON-TARGET SMARTpool siRNAs from Dharmacon) were used for knockdown of UAP56, SETX, FANCD2, DDX46, and HTATSF1 were used to achieve protein depletion. ON-TARGETplus Non-targeting Control Pool was used as the control (siC). A detailed list of siRNAs is in Table S3. Gene expression changes were assessed by qRT-PCR using gene-specific primers (Table S7). For the epistasis analysis experiment in K562 cells, siRNA transfections against Senataxin (siSETX), FANCD2 (siFANCD2), or siRNA against UAP56 (siUAP56), were performed at 72 after doxycycline induction (and repeated at 96 hours), and microscopy analysis was performed at 120 hours post-doxycycline induction.

Lentiviral gene-specific shRNAs were prepared in pLKO.1-puro backbone (Addgene # 8453) (identified from verified sequences at https://www.sigmaaldrich.com/US/en/product/sigma/shrna; TableS4). Lentiviral doxycycline-inducible shRNAs (targeting DDX46 and HTATSF1) were prepared in the Tet-pLKO-puro backbone (Addgene # 21915) by cloning the gene-targeting annealed oligos also identified from the above url into the AgeI - EcoR1 sites (details in TableS5). Lentivirus supernatants were prepared by co-transfecting the Tet-pLKO-puro plasmid along with helper plasmids pVSV-G (Addgene #138479) and p8.92 (Addgene #132929) into 293T cells as previously described. Supernatants were used as is or concentrated by PEG-8000 if required. Cells were spinfected (1800 RPM for 45 minutes at room temperature) in growth media supplemented with Polybrene (Yeasen), incubated at 37 C for 48 hours before selecting in 1 μg/ml puromycin for 5-7 days.

### Inducible lentiviral overexpression

Doxycycline-inducible gene overexpression vectors for DDX46 and hTATSF1 were prepared by cloning C-terminally HA-tagged DDX46 (Genscript, OHu29752) and hTATSF1 (amplified from K562 cDNA prepared using oligo(dT) primers) ORFs respectively into pLIX_403 (Addgene # 41395). Viral preps and selection were done as in previous section. Gene expression changes were assessed by qRT-PCR using transcript-specific primers (TableS7).

### Quantification of mRNA

RNA purification was performed using the RNeasy Mini kit (QIAGEN) according to the manufacturer’s conditions. cDNA synthesis was achieved using the M-MuLV Reverse Transcription kit (NEB), according to manufacturer’s guidelines. Finally, qPCR was performed on a 7500 FAST Real-Time PCR system (Thermo Fisher Scientific) and messenger RNA expression values calculated using the ΔΔ*C*t method and *Beta-Actin* housekeeping gene as a control. A detailed list with primers used in the present study is provided in Supplementary TableS7.

### Affinity purification of SF3B1-associated proteins

For FLAG-tagged expression and purification in HEK293T cells, 1.5 million cells per plate were seeded in 10-cm plates. Next day, 5 ug of expression plasmid (expressing either Flag-tagged SF3B1^WT^ or FLAG-SF3B1^K700E^) were transfected using TransIT-X2 (MirusBio) and harvested after 48 hours. One-third was used for total nuclear fraction and two thirds for chromatin fraction by affinity purification. Protease inhibitor cocktail (Millipore Sigma, 11836170001) and phenylmethylsulfonyl fluoride (PMSF) were added to all the buffers to prevent protein degradation. For total nuclear fraction, cells were first lysed in cell lysis buffer (lysed in 250 μL cytoplasmic lysis buffer (10 mM Tris-HCl pH 7.5, 0.05% NP40, 150 mM NaCl) by gently resuspending then incubating on ice for 5 minutes. Lysate was then layered on top of a 500 μL cushion of 24% sucrose in cytoplasmic lysis buffer and spun at 2,000 rpm for 10 min at 4°C. The supernatant (cytoplasm fraction) was removed, and the pellet (nuclei) were rinsed once with 500 μL PBS/1 mM EDTA. Nuclei were resuspended in a ‘NET’ resuspension buffer (40mM Tris pH7.5, 150mM NaCl, 0.05% NP-40, 5mM MgCl2). Chromatin-bound SF3B1-protein fraction was isolated using a previously published protocol^108^, with minor modifications. Briefly, the cell pellet was first lysed with 5 volumes of ice-cold E1 buffer for 1 volume of cell pellet and then centrifuged at 1,100 *× g* at 4 °C for 2 min. The supernatant (cytoplasmic fraction) was discarded, and the pellet resuspended with the same volume of E1 buffer, for a total of 2 washes, followed by incubation in E1 buffer for 10 min on ice and centrifugation at 1,100 *× g* at 4 °C for 2 min. The pellet was then resuspended in 2 volumes of ice-cold E2 buffer, centrifuged at 1,100 *× g* at 4 °C for 2 min, and the supernatant (nucleoplasmic fraction) discarded. The pellet was again resuspended with the same volume of E2 buffer used for the previous step, for a total of 2 washes, followed by incubation in E2 buffer for 10 min on ice and centrifugation at 1,100 *× g* at 4 °C for 2 min. The chromatin pellet was resuspended in a resuspension buffer (40mM Tris pH7.5, 150mM NaCl, 0.05% NP-40, 5mM MgCl2). Both the nuclear and chromatin pellet fractions were sonicated 3 times (5 seconds on, 20 seconds off; 30% amplitude; Sonic Dismembranator Model 500 (Fisher Scientific)) on ice. For co-immunoprecipitation of FLAG-tagged SF3B1, nuclear extracts were incubated with ANTI-FLAG® M2 affinity gel (Sigma-Aldrich) for 8 hours at 4°C with gentle shaking, beads washed five times with pre-chilled NET resuspension buffer and eluted under native conditions by competition using 3x FLAG peptide (Millipore Sigma, F4799). Samples were boiled in the Laemmli SDS sample buffer (Thermofisher) at 95 C for 5 minutes.

### Isolation and analyses of chromatin associated proteins

All steps were performed on ice, and all buffers contained PMSF and 1x Roche protease inhibitor mix. Briefly, 40 million cells per sample were rinsed once with PBS/1 mM EDTA, then lysed in 250 μL cytoplasmic lysis buffer (10 mM Tris-HCl pH 7.5, 0.05% NP40, 150 mM NaCl) by gently resuspension, and incubated on ice for 5 minutes. Lysate was then layered on top of a 500 μL cushion of 24% sucrose in cytoplasmic lysis buffer and spun at 2,000 rpm for 10 min at 4**°**C. The supernatant (cytoplasm fraction) was removed, and the pellet (nuclei) were rinsed once with 500 μL PBS/1 mM EDTA. Nuclei were resuspended in 100 μL nuclear resuspension buffer (20 mM Tris-HCl pH 8.0, 75 mM NaCl, 0.5 mM EDTA, 0.85 mM DTT, 50% glycerol) by gentle flicking, then lysed by the addition of 100 μL nuclear lysis buffer (20 mM <u>HEPES</u> pH 7.5, 1 mM DTT, 7.5 mM MgCl_2_, 0.2 mM EDTA, 0.3 M NaCl, 1 M Urea, 1% NP-40), vortexed for twice for 2 seconds each, and then incubated on ice for 3 min. Chromatin was pelleted by spinning at 14,000 rpm for 2 min at 4**°**C. The supernatant (nucleoplasm fraction) was removed, and the chromatin was rinsed once with PBS/1 mM EDTA. Chromatin was immediately dissolved in 100 ul of resuspension buffer (40mM Tris pH7.5, 150mM NaCl, 0.05% NP-40, 5mM MgCl2). Both nucleoplasmic and chromatin fractions were sonicated 3 times (5 seconds on, 20 seconds off; 30% amplitude; ultrasonic probe sonicator) on ice. Protein concentrations were determined using BCA assay and sample volumes adjusted. Samples were boiled in Laemmli SDS sample buffer (Thermofisher) at 95 C for 5 minutes.

### In vitro splicing assay

For SF3B1 overexpression in HEK293T cells, 1.5 million cells per plate were seeded in 10-cm plates for a total of 4 plates per condition. Next day, 5 ug of expression plasmid, per plate, (expressing either SF3B1^WT^ or SF3B1^K700E^) were transfected using TransIT-X2 (MirusBio) and cells were harvested after 48 hours. Small-scale nuclear extracts (NEs) were prepared using a previously published protocol^109^. Protein concentrations in the dialyzed NEs were confirmed to be similar between the two conditions. The pSP72-AdML plasmid (Addgene#11242) was linearized using BamH1 enzyme digestion and the linearized template was purified by running in an agarose gel. [^32^P]-labelled AdML pre-mRNA was synthesized in vitro using the T7 HiScribe Transcription Kit (NEB) using [α-^32^P] UTP (Perkin Elmer, BLU007H250UC), and purified using ethanol precipitation. The NEs were mixed with either DMSO (both SF3B1^WT^ or SF3B1^K700E^) or PlaB (Cayman Chemical, Cay16538-100) in DMSO and pre-incubated at 30°C for 10 minutes. Splicing reactions contained 15 ul of NEs diluted in dialysis buffer (20mM HEPES-KOH (pH8), 100mM KCl, 0.2mM EDTA, 20% glycerol) supplemented with 3mM MgCl_2_, 1 mM ATP, 10 mM creatine phosphate, 2.6% PVA (added at the last), 1ul of Murine RNAase Inhibitor (40U/uL), and 10ng of [^32^P]-labelled pre-mRNA in a total of 25 ul, per reaction. Splicing reactions were then incubated at 30 °C for the indicated time points. For the kinetic analysis of the spliceosome complexes, 5 ul of each reaction was mixed with 2.5ul of 10x Heparin loading dye (6.5mg/ml Heparin, 40%w/v glycerol, 0.5% bromophenol blue, 0.5% xylene cyanol) and incubated on ice, and the reactions were resolved on 1.5% agarose gel in 0.5x TBE by native gel electrophoresis for 3.5 hours at 70V. The agarose was subsequently fixed in a fixation solution (10% acetic acid, 10% methanol in water). Gels were dried in vacuum gel dryer at 80°C for up to 2 hours. The gels were scanned using a Typhoon phosphorimager (Molecular Dynamics).

### Cell Synchronization and analysis

K562 cells were synchronized as previously reported. Briefly, for G_2_/M, cells were cultured in 40 ng/ml nocodazole (Sigma-Aldrich) for 20 h. For G_1_/S arrest, double thymidine blockade was used (2mM Thymidine for 20 h, washed and recultured for 9 hours, and blocked a second time for 20 h). Cell cycle status was verified by FACS flow analysis.

### Flow cytometry and cell cycle studies

CytoPhase Violet (Biolegend) was used for DNA content evaluation by FACS by staining for 90 min per manufacturer’s instructions. For immunophenotyping, cells washed in PBS/EDTA/BSA were typically incubated for 30minutes at 4 degrees and washed in same buffer 3 times. Antibodies used for FACS analysis and flow sorting are listed in TableS8. At least 20,000 events were recorded on a FACS LSR II HTS-1 flow cytometer (BD Biosciences) and analyzed using FlowJo software.

### Cell growth kinetics

To determine cell growth kinetics, equal number of cells (1 million per condition) were plated in 3 replicates. Cell counts were performed using a Nexcelom cellometer cell counter every 7 days. Cell viability was independently determined by quantifying cellular ATP, using Cell Titre Glo 2.0 Cell Viability Assay (Promega) per manufacturer’s instructions.

### DNA fiber combing assay

DNA combing was performed as previously described^61^. Briefly, exponentially growing K562 cells (3.0 x 10^5^ per condition) were labeled with consecutive pulses of thymidine analogues (IdU and CldU) for 30-min. After labeling, cells were harvested, embedded in agarose and DNA was prepared then combed onto silanized coverslips using the FiberComb Molecular Combing System (Genomic Vision). Following combing, IdU and CldU were detected using mouse anti-BrdU (BD Biosciences 347580) and rat anti-BrdU (Abcam ab6326) antibodies, respectively. Sheep anti-mouse Cy3 (Sigma C2181) and goat anti-rat Alexa Fluor 488 (Invitrogen A11006) were used as secondary antibodies. Single-stranded DNA was counterstained with anti-ssDNA mouse antibody (DSHB University of Iowa), followed by anti-mouse BV480 (Jackson ImmunoResearch 115– 685-166) as secondary antibody. DNA fiber images were acquired using an Olympus FV3000 confocal microscope. Track lengths were measured with ImageJ. To calculate replication fork velocity (kb/min), the following equation was used to convert fork length from mm to kb/min: length mm’ 2/labeling time in min = fork speed kb/min (conversion factor of 2 kb/mm specific for DNA combing method). For quantification of fork velocity, a minimum of 150 fibers were analyzed per dataset.

### Proximity Ligation assay

PLA was performed as previously described using Duolink PLA Technology (Merck)^51^. Briefly, cytospin slides were pre-extracted, fixed, permeabilized and incubated with primary antibodies as described above for IF. Secondary antibody binding, ligation and amplification reactions were then performed according to the Duolink instructions. Duolink in situ PLA probe anti-rabbit PLUS (Merck) and Duolink in situ PLA probe anti-mouse MINUS (Merck), and Duolink Detection Reagents Red or Green (Merck) were used to perform the PLA reaction. Finally, nuclei were stained with DAPI and mounted in ProLong Gold AntiFade reagent (Cell Signaling Technology, 8961S). For PLA reactions requiring mouse FANCD2, mouse PCNA, rabbit PCNA, or rabbit RNAPII NTD antibodies, 1:200 dilution of these antibodies was used. Finally, images were acquired with a SP8 STED microscope equipped with a DFC390 camera (Leica) at 60× magnification and LAS AX image acquisition software (Leica). A FIJI image processing package^110^ was used for image analysis and quantification. PLA foci numbers per cell were quantified for all conditions.

### EdU and EU incorporation and detection

EdU (Catalog No. C10337) and EU (Catalog No. C10329) labeling were performed using Click-IT chemistry kits from Invitrogen. First, cells were cultured in complete medium supplemented with 10 μM EdU for 30 min. Then, samples were fixed, permeabilized and Click-iT reaction performed according to the manufacturer’s guidelines. Finally, nuclei were stained with DAPI and mounted in ProLong Gold AntiFade reagent (Invitrogen). Images were acquired with a Leica DM6000 microscope equipped with a DFC390 camera (Leica) at 63× magnification and LAS X image acquisition software (Leica). The FIJI image processing package was used for image analysis and quantification. The EdU entire population nuclear intensity and percentage of cells incorporating EdU were determined. EdU intensity was also determined only for those cells that incorporated EdU.

### Alkaline single-cell electrophoresis

Alkaline single-cell electrophoresis or comet assay was performed as previously described^111^. Comet slides were stained with SYBRsafe, and images were captured with a Leica DM6000 microscope equipped with a DFC390 camera (Leica). Analysis of comet assay images was performed using FIJI. Tail moment was measured for each cell in each condition.

### Immunofluorescence (IF)

K562 cells were cytospun at 400 RPM x 3 minutes onto glass coverslips coated with Poly-L-lysine (Gibco) in a Thermoshandon Cytospin 3 centrifuge. S9.6 IF was performed essentially as previously described^50^. Briefly, cells were fixed with 100% ice-cold methanol x 15 minutes, blocked with phosphate-buffered saline–bovine serum albumin (PBS–BSA) 2% overnight at 4 °C and incubated with S9.6 (1:1,000) and anti-nucleolin (1:2,000) antibodies overnight at 4 °C. Then, coverslips were washed three times in PBS 1× and incubated with secondary antibodies (1:1,000) for 1 hr at room temperature (RT). Finally, cells were washed again, stained with DAPI (1:1000, Thermofisher) and mounted in ProLong Gold AntiFade reagent (Cell Signaling Technology, 8961S). IF images were acquired with a Leica DM6000 microscope equipped with a DFC390 camera at 63× magnification and LAS X image acquisition software (Leica). A FIJI (ImageJ) image processing package was used for IF analysis. The nuclear mean gray value for S9.6, after subtraction of nucleolar signal, was measured for each condition.

DNA-damage assessment by γH2AX immunostaining was performed mainly as previously described^112^ with minor modifications. All cells were fixed in paraformaldehyde for 20 min, washed twice in PBS, permeabilized with PBS containing 0.1% Triton-X100, and blocked in PBS + 2% bovine serum albumin. Immunofluorescence was performed using anti-γH2AX (1:1000) primary antibody overnight at 4 °C. Coverslips were washed three times with PBS and incubated with Alexafluor secondary antibodies (1:1000) and counterstained with DAPI (1:1000, Thermofisher) for 1 h at room temperature. Coverslips were mounted on glass slides using ProLong Gold anti-Fade mount (Cell Signaling Technology, 8961S) and cured overnight. IF images were acquired with a Leica DM6000 microscope equipped with a DFC390 camera at 63× magnification and LAS AX image acquisition software (Leica). A FIJI (ImageJ) image processing package was used for IF analysis.

### Dot-blot assay

Cells were lysed in 0.5% SDS/TE, pH 8.0 containing Proteinase K overnight at 37 °C. Total chromatin was isolated with phenol/chloroform/isoamyl alcohol extraction followed by standard ethanol precipitation. The eluted chromatin was digested for one hour in RNaseIII (NEB, M0245), purified with MinElute PCR purification kit (Qiagen, 28004), and quantified using Nanodrop. A negative control treated for 4 hours at 37 °C with RNaseH1 (New England Biolabs) was included for each condition. 300 and 600 nanograms of total DNA was loaded in duplicate onto a Nylon membrane using dot blot apparatus, UV crosslinked wit at 0.12 Joules and stained with methylene blue to determine uniform loading. Membranes were then blocked with 5% skim milk in PBST (PBS; 0.1% Tween-20) for 1 hour and RNA:DNA hybrids were detected by immunoblotting with S9.6 antibody.

### ChIP-seq

ChIP-seq was performed as previously described^113^ with 1-5 × 10^7^ cells from two biological replicates. Briefly, formaldehyde (16% concentrate stock methanol-free, Cell Signaling Technology, 12606) was directly added to the media to a final concentration of 1% and incubated for 15 min. 125 mM Glycine (final concentration) was added to quench the reaction for 5 min. Cells were spun down and pellet was washed twice with PBS at 4°C. Protease (Millipore Sigma, 11836170001) inhibitors were added to all the buffers. The cell pellet was lysed on ice in cell lysis buffer (5 mM Pipes pH 8, 85 mM KCl, 0.5% NP-40). Pellet was centrifuged for 5 min at 1,700 g at 4°C. Pellet was resuspended with 0.5 mL of MNase digestion buffer (10 mm Tris pH 7.4, 15 mM NaCl, 60 mM KCl, 0.15mM Spermine, 0.5mM Spermidine) containing MNase (NEB, M0247S) to final concentration of final concentration of 20 U/µL and incubated at 37°C for 15 min. MNase was inactivated by the addition of EDTA (final concentration of 50mM) and brief incubation on ice. Pellet was centrifuged for 1 min at 16000g at 4°C. Pellet was resuspended in 0.5mL of nuclear lysis buffer (50mM Tris pH 8, 10mM EDTA, 0.5% SDS).

Sonication was performed with a sonic Dismembranator Model 500 (Fisher Scientific) with the following parameters: amplitude 30%, 5 seconds on, 15 seconds off, 16 cycles. Sheared chromatin was centrifuged at 10,000 g for 15 min at 4°C. 15 μL of samples were de-crosslinked overnight at 65°C, size distribution was checked in 1% agarose gel and chromatin concentration measured using Nanodrop^TM^. 5-10 μg of chromatin was used for each IP for RNAPII NTD and pSer2 RNAPII ChIP assay, and 100 μg of chromatin was used for each IP for CDK9 ChIP assay. ChIP spike in normalization for the RNAPII ChIP assays was performed using Drosophila-specific chromatin and *Drosophila*-specific histone variant (H2Av) antibody^114^. 50 ng of Drosophila S2 sheared crosslinked-chromatin (Active Motif, 53083) was added to 10 μg of chromatin as spike-in control. Chromatin was diluted with IP buffer (16.7mM Tris pH 8, 167 mM NaCl, 1.2 mM EDTA, 0.01% SDS, 1.1% TritonX-100) to obtain a 0.1% final concentration of SDS. 5% of diluted chromatin was kept as input at 4°C. 10 μg of RNAPII NTD (D8L4Y) antibody (Cell Signaling Technology, 14958), 10 μg of <u>pSer2</u> RNAPII (E1Z3G) antibody (Cell Signaling, 13499)), 10μg of CDK9 (C12F7) antibody (Cell Signaling Technology, 2316), or 1 μg of <u>histone</u>H2Av (Active Motif, 39715) was mixed with diluted chromatin and subjected to IP performed on a rotating wheel at 4°C overnight. Pierce Protein A/G Magnetic beads (ThermoScientific, 78609) were then added to the chromatin-antibody mixture and 4°C rotation continued for 3 more hours. Beads were washed once with wash buffer 1 (0.1% SDS, 1% TritonX-100, 2mM EDTA, 20 mM Tris pH 8, 150 mM NaCl), four times with wash buffer 2 (0.1% SDS, 1% TritonX-100, 2mM EDTA, 20 mM Tris pH 8, 500mM NaCl), once with wash buffer 3 (0.25M LiCl, 1% NP-40, 1 mM EDTA, 10 mM Tris pH 8), once with wash buffer 4 (10 mM Tris pH 8, 1 mM EDTA). Immuno-bound chromatin was eluted at 55°C for 1 hour with elution buffer (50 mM Tris pH 8, 10 mM EDTA, 1%SDS, proteinase K 7 μg/mL) and de-crosslinked overnight at 65°C. DNA was extracted with one volume phenol:chloroform:isoamyl alcohol 25:24:1 (Sigma-Aldrich, P2069) and precipitated for 60 min at −20°C with sodium acetate and ethanol. Pellet was washed with 70% ethanol and resuspended in TE buffer. DNA quality and size distribution were checked on Fragment Analyzer. 10 ng of DNA was used for library preparation according to NEBNext® Ultra II DNA Library Prep Kit (NEB, E7645S). Purity and size distribution of the libraries were estimated using Fragment Analyzer. Size-selected libraries were sequenced on Illumina NovaSeq S4 (paired-end 2×150) to average depth of 50 million reads.

### GRO-seq

GRO-seq was performed as previously described^23^ with minor modifications. 20 million cells from two biological replicates were used for each GRO-seq assay. Briefly, cells were washed twice with ice-cold PBS before adding swelling buffer (10 mM Tris-HCL pH 7.5, 2mM MgCl2, 3 mM CaCl2, 2U/ml Superbase-in (Invitrogen)). Cells were swelled for 5 min on ice, washed with swelling buffer + 10% glycerol and then lysed in lysis buffer (10 mM Tris-HCL pH 7.5, 2 mM MgCl2, 3 mM CaCl2, 10% glycerol, 1%l Igepal (NP-40), 2 U/ml Superasein) to isolate nuclei. Nuclei were washed twice with lysis buffer and resuspended in freezing buffer (40% glycerol, 5 mM MgCl2, 0.1 mM 0.5M EDTA, 50 mM Tris-HCL pH 8.3) to a concentration of 2×10^7^ nuclei per 100 mL. Nuclei were then frozen in dry ice and stored at −80°C until further use. Nuclei were thawed on ice just prior to the nuclear run-on. An equal volume of pre-warmed nuclear run-on reaction buffer (10 mM Tris-HCl pH 8, 5 mM MgCl2, 300 mM KCl, 1 mM DTT, 500 mM ATP, 500 mM GTP, 500 mM 4-thio-UTP, 2 mM CTP, 200 U/ml Superasein, 1% Sarkosyl (N-Laurylsarcosine sodium salt solution) was added and incubated for 7 min at 30°C for the nuclear run-on. Nuclear run-on RNA was extracted with TRIzol LS reagent (Invitrogen) following the manufacturer’s instructions and ethanol precipitated. NRO-RNA was resuspended in water and concentration was determined with Nanodrop^TM^. Up to 150 mg of RNA was transfer to a new tube and 5% (w/w) of Drosophila S2 spike-in RNA was added. RNA was then chemically fragmented by mixing with 2X RNA fragmentation buffer (150 mM Tris pH 7.4, 225 mM KCl, 9 mM MgCl_2_) and incubated at 94°C for 3 min 30 sec, before placing it back on ice and adding EDTA (to a final concentration of 50 mM) and incubating on ice for a further 2 minutes. Fragmentation efficiency was analyzed by running fragmented and unfragmented RNA on Agilent 2200 TapeStation using High Sensitivity RNA ScreenTapes following manufacturer’s instructions. Fragmented RNA was incubated in Biotinylation Solution (20 mM Tris pH 7.5, 2 mM EDTA pH 8.0, 40% dimethylformamide, 200 mg/ml EZ-link HPDP Biotin (Thermo Scientific, 21341)) for 2h in the dark at 25°C, 800 rpm. After ethanol precipitation, the biotinylated RNA was resuspended in water and biotinylated-RNA was separated with M280 Streptavidin Dynabeads (Invitrogen). 100 ul/sample of beads were washed twice with 2 volumes of freshly prepared wash buffer (100 mM Tris pH 7.5, 10 mM EDTA pH 8.0, 1M NaCl, 0.1% (v/v) Tween-20) and resuspended in 1 volume of wash buffer and added to the biotinylated-RNA. After 15 min in rotation at 4°C, beads were washed three times with wash buffer pre-warmed at 65°C and three times with a room temperature wash buffer. 4-S-UTP containing RNA was eluted in 100 mM DTT buffer and purified with RNA Clean and Purification kit (Zymo Research, R1013) with in-column DNAse reaction to eliminate traces of genomic DNA. The eluted RNA was quantified with Qubit High Sensitivity Assay kit (Invitrogen) and used to produce barcoded cDNA sequencing libraries using the SMARTer® Universal Low Input RNA Kit (Takara, 634938). Libraries were sequenced on Illumina NovaSeq S4 (paired-end 2×150) to average depth of 50 million reads per sample.

### TT-TL seq

TT-TL-seq was performed as described^24^, with a few modifications (primarily using HDPD biotinylation instead of MTSEA-biotin). About 15 million cells per sample was labeled with s^4^U (by adding it to culture media at final concentration of 500 μM) for 5 minutes. Cells were harvested and washed in ice cold PBS and RNA extracted by Trizol and chloroform extraction, ethanol precipitation (1 mM DTT added to prevent oxidation of the s^4^U RNA). Total RNA was resuspended and treated with TURBO DNase, then extracted with acidic phenol:chloroform:isoamyl alcohol and again ethanol precipitated. RNA was resuspended in water and concentration was determined with Nanodrop^TM^. Up to 150 mg of RNA was transferred to a new tube and 5% (w/w) of Drosophila S2 spike-in RNA was added. RNA was then chemically fragmented by mixing with 2x RNA fragmentation buffer (150 mM Tris pH 7.4, 225 mM KCl, 9 mM MgCl_2_) and heating to 94°C for 3 min 30 sec, before placing it back on ice and adding EDTA (to a final concentration of 50 mM) and incubating on ice for a further 2 minutes. Fragmentation efficiency was analyzed by running fragmented and unfragmented RNA on Agilent 2200 TapeStation using High Sensitivity RNA ScreenTapes following manufacturer’s instructions. Fragmented RNA was incubated in Biotinylation Solution (20 mM Tris pH 7.5, 2 mM EDTA pH 8.0, 40% dimethylformamide, 200 mg/ml EZ-link HPDP Biotin (Thermo Scientific, 21341)) for 2h in the dark at 25°C, 800 rpm. After chloroform: isoamyl alcohol (24:1) extraction followed by purification (Qiagen RNeasy Mini Kit, 74104), the biotinylated RNA was resuspended in water and biotinylated-RNA was isolated using Dynabeads MyOne Streptavidin C1 magnetic beads (Invitrogen), as previously described^24^. s^4^U-biotinylated RNA was eluted from the beads by cleaving the HPDP Biotin in elution buffer (100mM DTT, 20 mM DEPC, pH 7.4, 1 mM EDTA, 100 mM NaCl, 0.05% Tween-20) and cleaned up using 1x RNAcleanXP beads (Beckman Coulter, A63987). For time lapse chemistry, 20 ul of isolated s^4^U-enriched RNA was added to a mixture of 0.835 μl 3M sodium acetate pH 5.2, 0.2 μl 500 mM EDTA, 1.365 μl RNAse free water, and 1.3 μl trifluoroethylamine. 1.3 μl of NaIO_4_ (200 mM) was then added dropwise, and the reaction mixture was incubated for 1 h at 45 °C. 1x volume of RNA clean beads were then added and purified as described above. 2 μl of reducing master mix (58 μl of water, 10 μl 1M DTT, 10 μl 1M Tris pH 7.4, 2 μl 0.5mM EDTA, 20 μl 5M NaCl) was added to each sample for a final volume of 20 μl, mixed, and incubated in thermocycler at 37C for 30 min. The mixture was then mixed with 1x RNAclean beads and RNA purified, as described above. The eluted RNA was quantified with Qubit High Sensitivity Assay kit (Invitrogen) and used to produce barcoded cDNA sequencing libraries using the SMARTer® Universal Low Input RNA Kit (Takara, 634938). Validation of TT-TL procedure was performed by qPCR (TableS7 for primer sequences). Libraries were sequenced on Illumina NovaSeq S4 (paired-end 2×150) to average depth of 50 million reads per sample.

### CUT&RUN seq

CUT&RUN was performed as previously described^115^. Briefly, 0.5 million K562 cells or 25,000 human CD34+ cells were harvested, washed, and bound to activated Concanavalin A-coated magnetic beads (activated in 20 mM HEPES, pH 7.9, 10 mM KCl,1 mM CaCl2,1mM MnCl2), then permeabilized with Wash buffer (20 mM HEPES, pH7.5, 150 mM NaCl, 0.5 mM spermidine and Roche protease inhibitor tablet) containing 0.05% Digitonin (Dig-Wash). The bead-cell slurry was incubated with 0.5 μl antibody (H3K4me3 (Cell Signaling Technology, 9751), H3K4me3 (Cell Signaling Technology, 5326), H3K27me3 (Cell Signaling Technology, 9733), H3K4me3 (Cell Signaling Technology, 9751), H3K27ac (Cell Signaling Technology, 8173), pSer2 RNAPII (Cell Signaling Technology, 13499), Rabbit IgG (Cell Signaling Technology, 2729)) in a 50 µL antibody buffer ((20 mM HEPES, pH 7.5, 150 mM NaCl, 0.5 mM Spermidine, 0.01% digitonin, 2 mM EDTA, protease inhibitor) at 4°C overnight on a nutator. After 3 washes in 1 ml Dig-wash, beads were resuspended in 50 µl digitonin wash, mixed with 2.5 µl 20x pAG/MNase (EpiCypher, 151016) and incubated for 15 min at room temperature. After two washes in Dig-wash, beads were resuspended in 150 µl digitonin wash. Tubes were chilled to 0°C and 1 µl 100 mM CaCl2 was added while gently vortexing. Tubes were nutated at 4°C for 2 hours followed by addition of 33 µl EGTA-STOP buffer (340 mM NaCl, 20 mM EDTA, 4 mM EGTA, 50 μg/mL RNase A, 50 μg/mL Glycogen with E.coli spike in control (Epicypher, 18-1401)).

Beads were incubated at 37°C for 30 min, replaced on a magnet stand and the liquid was removed to a fresh tube and DNA was extracted. DNA quality and size distribution were checked on Fragment Analyzer. 10 ng of DNA was used for library preparation according to NEBNext® Ultra II <u>DNA Library</u> Prep Kit (NEB, E7645S). Purity and size distribution of the libraries were estimated using Fragment Analyzer. Size-selected libraries were sequenced on Illumina NovaSeq S4 (paired-end 2×150). Size-selected libraries were sequenced on Illumina NovaSeq S4 (paired-end 2×150) to an average depth of 10 million reads per sample.

### ATAC-seq

ATAC seq was performed using the Omni-ATAC protocol^116^, optimized for low cell numbers. 50,000 K562 cells or 5000 CD34+ cells (flow sorted) were harvested, washed, and resuspended in cold PBS. Washed cells were resuspended in 1 ml of cold ATAC-seq resuspension buffer (RSB; 10 mM Tris-HCl pH 7.4, 10 mM NaCl, and 3 mM MgCl_2_ in water) and centrifuged at 500g for 5 min in a 4 °C fixed-angle centrifuge. Cell pellets were then resuspended in 50 μl of RSB containing 0.1% NP40, 0.1% Tween-20, and 0.01% digitonin by pipetting up and down three times. This cell lysis reaction was incubated on ice for 3 min. After lysis, 1 ml of RSB containing 0.1% Tween-20 (without NP-40 or digitonin) was added, and the tubes were inverted to mix. Nuclei were then centrifuged for 10 min at 500g at 4 °C. Supernatant was carefully removed and nuclei were resuspended in 50 μl of transposition mix (25 μl 2× TD buffer (20 mM Tris pH 7.6, 10 mM MgCl_2_, 20% Dimethyl Formamide) 2.5 μl transposase (100 nM final), 16.5 μl PBS, 0.5 μl 1% digitonin, 0.5 μl 10% Tween-20, and 5 μl water) by pipetting up and down six times. Transposition reactions were incubated at 37 °C for 30 min in a thermomixer with shaking at 1,000 rpm. Reactions were cleaned up with the MinElute PCR purification kit (Qiagen, 28004). DNA quality and size distribution were checked on Fragment Analyzer. 10 ng of DNA was used for library preparation according to NEBNext® Ultra II <u>DNA Library</u> Prep Kit (NEB, E7645S). Purity and size distribution of the libraries were estimated using Fragment Analyzer. Size-selected libraries were sequenced on Illumina NovaSeq S4 (paired-end 2×150).

### Nascent RNA-seq (long and short read)

Nascent RNA-seq for short- and long-read sequencing was performed on K562 cells (∼60 million per biological replicate), as described in previously published protocols^251^, with minor modifications. To minimize RNA degradation, all steps were performed on ice, and all buffers contained 25 uM α-amanitin, 40 U/ml RNase inhibitor (Invitrogen), and 1x Roche protease inhibitor mix. Briefly, 20 million cells were rinsed once with PBS/1 mM EDTA, then lysed in 250 μL cytoplasmic lysis buffer (10 mM Tris-HCl pH 7.5, 0.05% NP40, 150 mM NaCl) by gently resuspending then incubating on ice for 5 minutes. Lysate was then layered on top of a 500 μL cushion of 24% sucrose in cytoplasmic lysis buffer and spun at 2,000 rpm for 10 min at 4**°**C. The supernatant (cytoplasm fraction) was removed, and the pellet (nuclei) were rinsed once with 500 μL PBS/1 mM EDTA. Nuclei were resuspended in 100 μL nuclear resuspension buffer (20 mM Tris-HCl pH 8.0, 75 mM NaCl, 0.5 mM EDTA, 0.85 mM DTT, 50% glycerol) by gentle flicking, then lysed by the addition of 100 μL nuclear lysis buffer (20 mM <u>HEPES</u> pH 7.5, 1 mM DTT, 7.5 mM MgCl_2_, 0.2 mM EDTA, 0.3 M NaCl, 1 M Urea, 1% NP-40), vortexed for twice for 2 seconds each, and then incubated on ice for 3 min. Chromatin was pelleted by spinning at 14,000 rpm for 2 min at 4**°**C. The supernatant (nucleoplasm fraction) was removed, and the chromatin was rinsed once with PBS/1 mM EDTA. Chromatin was immediately dissolved in 100 μL PBS and 300 μL TRIzol Reagent (ThermoFisher). RNA was purified from chromatin pellets in TRIzol Reagent (ThermoFisher) using the RNeasy Mini kit (QIAGEN) according to the manufacturer’s protocol, including the on-column DNase I digestion. For genome-wide nascent RNA-seq, samples were depleted three times of polyA(+) RNA using the Magnetic mRNA isolation kit (New England Biolabs), each time keeping the supernatant, then depleted of ribosomal RNA using oligo probe-based rRNA depletion^117^. For short read sequencing, the isolated polyA, rRNA depleted RNA was then fragmented with a Bioruptor UCD-200 for 1-5 cycles of 30 s ON / 30 s OFF, high settings. Fragmentation efficiency was analyzed by running fragmented and unfragmented RNA on Agilent 2200 TapeStation using High Sensitivity RNA ScreenTapes following manufacturer’s instructions. The fragmented RNA was used to produce barcoded cDNA sequencing libraries using the SMARTer® Universal Low Input RNA Kit (Takara, 634938). Short-read libraries were sequenced on Illumina NovaSeq S4 (paired-end 2×150). For long read sequencing, a DNA adapter (TableS6) was ligated to 3′ ends of nascent RNA using the T4 RNA ligase kit (NEB) by mixing 50 pmol adapter with 300-600 ng nascent RNA. cDNA was generated from the adapter ligated RNA using the SMARTer PCR cDNA Synthesis Kit (Clontech), replacing the CDS Primer IIA with a custom primer complementary to the 3′ end adapter for first strand synthesis (TableS6). cDNA was amplified by 15 cycles of PCR using the Advantage 2 PCR Kit (Clontech) with template switching custom oligo (TableS6), cleaned up using a 1X volume of AMPure beads (Agencourt), then PacBio library preparation was performed by the Yale Center for Genome Analysis using the SMRTbell Template Prep Kit 1.0 (Pacific Biosciences). The library was sequenced on four RSII flowcells and four Sequel 1 flowcells.

### DRIP-seq

DRIP-seq was performed as described^66^. K562 cells (5 × 10^6^) were lysed in 0.5% SDS/TE, pH 8.0 containing Proteinase K overnight at 37 °C. Total DNA was isolated with phenol/chloroform/isoamylalcohol extraction followed by standard ethanol precipitation. One-third of total DNA was fragmented by a cocktail of restriction enzymes (EcoRI, HindIII, BsrgI, SspI, XbaI) overnight at 37 °C. A negative control treated overnight with RNaseH1 (New England Biolabs) was included. Digested DNA was purified by phenol/chloroform/isoamyl alcohol extraction, ethanol precipitation and quantified by Nanodrop^TM^. 4 μg of digested DNA were diluted in binding buffer (10 mM NaPO_4_, pH 7.0; 0.14 M NaCl; 0.05% Triton X-100) and incubated with 10 μg of S9.6 antibody overnight at 4 °C on a rotator. DNA/antibody complexes were added for 2 hours at 4 °C to Agarose Protein-A/G beads (ThermoScientific, 20421) prewashed with a binding buffer. Immunoprecipitated DNA was eluted by incubating with an elution buffer (50 mM Tris pH 8.0; 10 mM EDTA; 0.5% SDS) containing proteinase K at 55 °C for 45 min on a rotator. The eluent was precipitated by phenol/chloroform/isoamylalcohol extraction and ethanol precipitation. Validation of the DRIP procedure was performed by qPCR (TableS7 for primer sequences). The pulled down material and input DNA were then sonicated, size-selected, and ligated to Illumina barcoded adaptors, using TruSeq ChIP Sample Preparation Kit (Illumina) or ThruPLEX® DNA-seq Kit (Rubicon Genomics) for next-generation sequencing (NGS) on Illumina HiSeq 2500 platform. NEBNext® Ultra II Prep Kit (NEB, E7645S). Purity and size distribution of the libraries were estimated using Fragment Analyzer. Size-selected libraries were sequenced on Illumina NovaSeq S4 (paired-end 2×150) to an average depth of 50 million reads per sample.

### RNA-seq

RNA was purified, fragmented and size checked on a 2100 Bioanalyzer (Agilent). Libraries were built using TruSeq Stranded Total RNA Library Prep (Illumina) after rRNA depletion according to the manufacturer’s protocol. RNA-seq libraries were prepared using the Illumina TruSeq Stranded mRNA Library Prep Kit. Paired-end RNA-seq were performed with Illumina NovaSeq S4 (paired-end 2×150) to an average depth of 100 million reads per sample.

### Bioinformatic analysis

For all next generation sequencing datasets, the quality of the paired-end sequencing data was assessed with FastQC^118^. ATAC-, ChIP-, DRIP-, GRO-seq, TT-TL-seq data and Drosophila S2 spike in (for RNAPII ChIP) were trimmed with Cutadapt v4.1^119^ to remove adaptor sequences. The trimmed reads were aligned to the canonical version of human reference genome hg38 and Drosophila genome (UCSC, dm6) using specific aligners as indicated for each assay.

ChIP-seq: Bowtie2^120^

GRO-seq: Bowtie2 and STAR^121^ (for splicing analysis)

TT-TL-seq: Bowtie2 and STAR (for splicing analysis) nSRS-seq: Bowtie2

CUT&RUN: Bowtie2 RNA-seq: STAR

Mapping quality in the Bam files was assessed with SAMtools^122^. Then, PCR duplicates were removed, and the resulting reads were extended to 150 bases toward the interior of the sequenced fragment and were normalized to total reads aligned (reads per kilobase per million, RPKM). In cases when stranded libraries were built (GRO-seq, RNA-seq), reads assigned to Watson and Crick strands using SAMtools. For assessing correlation between biological replicates to determine reproducibility, the paired-end sequencing reads were mapped to human genome using Bowtie2. Average read coverages were calculated by feeding resulting bam files into ‘multiBamSummary’ program (deepTools Version 3.5.1) which was subsequently plotted using ‘PlotCorrelation’ program (deepTools Version 3.5.1). Average coverages, RPKM-normalized bigwig files, and metaplot images were generated using deepTools 2.0^123^. Tracks were visualized using IGV^124^and generated using Gviz^125^. Further analyses were done in R (http://www.R-project.org), with Bioconductor packages and ggplot2^126^ for graphic representation. With regard to RNA-seq, counts were assigned to every gene as mentioned and RPKM normalized for each replicate. After entire population analysis, those with an RPKM average from both replicates >0.001 were considered as expressed genes. rMATS^127^ package was used to identify alternative splicing events from RNA-seq data. For comparative purposes in nascent SRS-seq, GRO-seq and RNA-seq datasets, counts per gene were calculated using FeatureCounts^128^ for each condition and replicate. The raw count values were then subjected to default library-size normalization and differential expression analysis using DESeq2^129^.

Heat maps showing the changes in RNAPII NTD, Ser2P RNAPII CTD, and H3K4me3 CUT&RUN enrichments upon SF3B1^K700E^ expression in K562 cells were constructed by subtracting the averaged signal enrichments (n=2) in SF3B1^K700E^ from the corresponding averaged signal enrichments (n=2) in WT using the ‘-- operation subtract’ option in the deepTools ‘bamCompare’ program. Unsupervised K clustering was performed on the RNAPII NTD and RNAPII PS2 changes, in the −1 kb upstream of TSS to 2 kb downstream of the TES region, to separate active genes into three distinct clusters (number of clusters was determined using R-NbClust package).

For DRIP-seq and ATAC-seq data, Epic2^130^ package was used to call differentially enriched domains in WT and SF3B1^K700E^, using default parameters. Differentially DRIP-seq and ATAC peaks were determined as those with a |log_2_ fold change(L_2_FC)| > 0.58 and adjusted p value < 0.05, calculated with DESeq2. Genome and gene annotation of peaks was performed using the HOMER v4.11 package^131^ and genes retrieved from Gencode version 29 release (2018). Volcano plots were generated using ggplot2 on L_2_FC and adjusted p values. Convergent gene-pairs were identified and retrieved from Gencode version 29 release (2018) using an in-house python script.

The analyses of replication fork directionality and replication initiation zones used the published OK-seq data^132^ in K562 cells from the ENCODE database. Bamscale^68^ was used to create RFD scores from the OK-seq data and scaled coverage tracks for visualization in genome browsers. RFD values were measured for each R-loop peak to establish R-loop orientation with respect to the replication fork (by intersecting R loop peaks with replication forks using BEDTools^133^). For purposes of RFD collision analysis, assays involving RFD were performed only on those sites (R-loops) where |RFD| > 0.5, ensuring a high chance of collision in a specific orientation.

### Long read sequencing analysis

Long Read Sequencing analysis was performed as previously reported^1^, utilizing Porechop v0.2.4 (https://github.com/rrwick/Porechop) to remove external adapters, Cutadapt for removing 3’ adapters and Prinseq-lite^134^ to remove PCR duplicates. Minimap2^135^ was used to align to hg38 build. Custom scripts in Python and R were used to generate Co-SE and cumulative distribution of 3’ RNAPII position.

### CUT&RUN seq analysis

Paired-end reads were aligned to the human reference genome hg38 using Bowtie2. Mapping quality in the Bam files was assessed with SAMtools. PCR duplicates were removed, and the resulting reads were normalized to total reads aligned (RPKM). Average coverages, RPKM-normalized bigwig files, and metaplot images were generated using deepTools 2.0. Tracks were generated using Gviz. Further analyses were done in R (http://www.R-project.org), with Bioconductor packages and ggplot2 for graphic representation. Peak calling was performed using SEACR package^136^ using control IgG CUT&RUN bedgraph files to generate empiric thresholds.

### TT-TL-seq analysis

TT-TL-seq reads from SF3B1^WT^, SF3B1^K700E^ and U2AF1^WT^, U2AF1^S34F^ cells containing spiked-in RNA from *Drosophila* S2 cells were trimmed with Cutadapt to remove adaptor sequences. Trimmed reads were mapped to a combined hg38 and dm6 genome using Bowtie2. Bam files for uniquely mapped reads (SAM flag 83/163 or 99/147) were created, sorted and indexed using Samtools. To calculate normalization factors, reads aligning to the dm6 genome were filtered and counted with Samtools. RPKM-normalized, Drosophila-scaled bigwig files and metaplot images were generated using deepTools 2.0. To assess the TL-component in the TT-TL seq, unfiltered, trimmed reads were mapped to a combined hg38 and dm6 genome using HISAT-3N^137^ with default parameters and U-to-C mutation calls. Normalized coverage tracks were generated using Gviz.

For accurate differential expression analysis using DESeq2, normalization factors for the sample libraries were calculated with FeatureCounts, using readcounts mapped to the dm6 genome in addition to library size normalization. Differential expression analysis of normalized read counts was performed in gene bodies (TSS+300bp to TES).

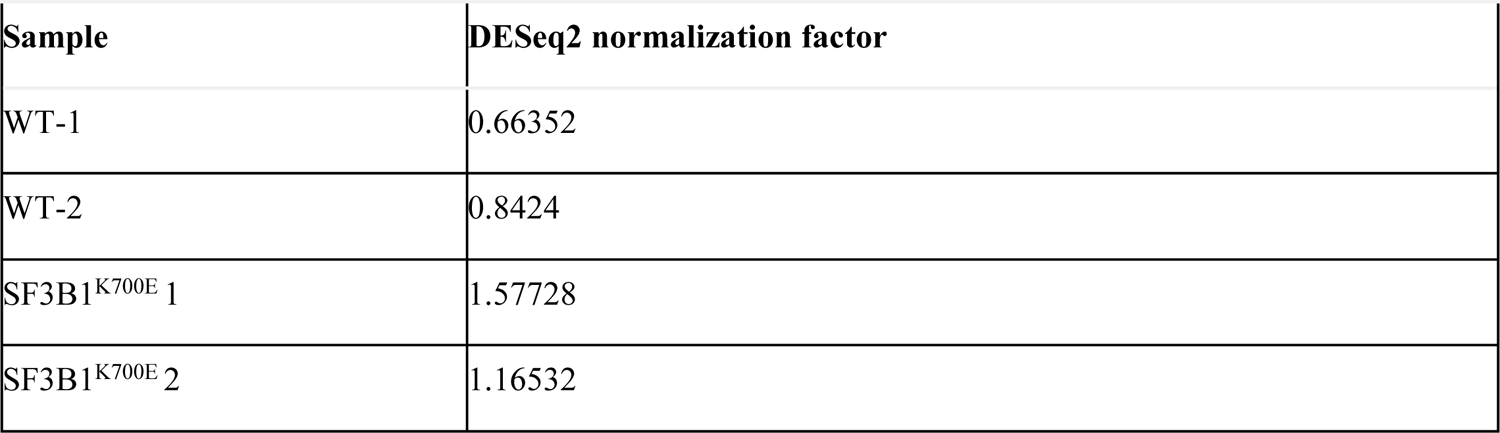

Clusterprofiler was used for gene ontology enrichment analysis of biological enrichments. Read-in and read-through values were generated for TT-TL-seq (reads with 1 mutation or more) as well as GRO-seq using ARTDeco^138^.

### Identification of promoter and enhancer analysis

Genome annotation and reference genome sequences were downloaded from the UCSC Genome Browser. Only those RefSeq-validated transcripts with a GRO-seq signal at the region between TSS – 10 bp and TSS + 300 bp were selected. Transcripts that are ≤ 1 kb long and ≤ 1 kb from the nearest genes were removed. TSS of antisense transcription was defined by the highest GRO-seq signal in the antisense strand within 500 bp upstream and downstream of the sense TSS. The highest GRO-seq signal within the region from +20 to +300 bp downstream of TSS was defined as the pausing site. The promoter was defined as the region between TSS – 100 bp and TSS + 300 bp.

For putative enhancer identification, distal H3K27ac peaks (determined from K562 CUT&RUN data), without overlap of the region from 2kb upstream and downstream of the protein coding genes, were defined as enhancers and the strand having higher GRO signal was defined as the sense direction. The pausing site of the enhancer was defined as described above, for promoters.

### Chromatin state analysis

ChromHMM^84^ was used with default parameters to derive genome-wide chromatin state maps for all cell types. Input ChIP-seq, ATAC-seq, DRIP-seq, and CUT&RUN data were binarized with ChromHMM’s BinarizeBed method using a p-value cutoff of 10^−4^. Chromatin state models were learned jointly on all the marks from SF3B1^WT^ and SF3B1^K700E^ from 10 to 15 states. A model with 14 states was chosen for detailed analysis. Chromatin state annotations of SF3B1^WT^ and SF3B1^K700E^ were produced subsequently by applying this model to the original binarized chromatin data from these cell types. To determine important chromatin state changes between WT and SF3B1^K700E^, the chromatin state annotations were intersected. In each case, the number of 200bp bins that is occupied by each of the 14 by 14 possible chromatin state transitions. To calculate enrichment scores, this number was divided by the expected number of such bins assuming a null model that treats the two chromatin states involved in each transition as being independently distributed.

For promoter state analysis, promoters were defined as regions −1Kb to +1Kb proximal to TSS. Every RefSeq gene was assigned to one chromatin state based on the state call on the gene’s TSS in WT and SF3B1^K700E^ cell types by using the 14 state ChromHMM genome annotation output.

### Unsupervised clustering of Alternative Splicing (AS) events

RNA-seq files were aligned to GRCh8 by STAR and bam files were used to determine percentage spliced in (PSI) for each sample using rMATS. Splicing events were aggregated and those with at least PSI of 0.1 in 10% samples (22 of 217 total) were used for unsupervised clustering using ComplexHeatmap package^139^ in R. To optimize performance, 10,000 random events (from total 137,435 that met PSI criteria) were chosen for clustering. Default parameters (Pearson distance method) were used to cluster by rows (AS events). Annotations for disease status (MDS/AML or normal) and splicing status of SF3B1, U2AF1, SRSF2 and ZRSR2 were provided.

### shRNA screen

∼8 million K562 cells, per replicate, were transduced with a HIV-based lentiviral RNAi library^140^ containing a total of 2343 shRNAs (targeting 357 genes) at a MOI of 0.2 and a target representation of 3000 cellular integrations for each shRNA. After transduction, cells were selected for 4 days in puromycin, then grown in doxycycline, and cultured for ∼12 population doublings (24 days). During this time, cells were passaged every 4 days. Cells were collected immediately after puromycin selection (day 0) and subsequently on days 8, 16 and 24 after doxycycline induction (2 million cells each). The QIAamp DNA Blood Midi kit (Qiagen) was used to extract genomic DNA from cell pellets followed by ethanol precipitation to clean and concentrate. Half-hairpin sequences were amplified from the genomic DNA (common forward primers 5′- AATGATACGGCGACCACCGAGATCTACACTCTTTCCCTACACGACGCTCTTCCGATCTTCTTGTGGAAAGGACGA-3′ and reverse primer 5′-CAAGCAGAAGACGGCATACGAGATNNNNNNNNGTGACTGGAGTTCAGACGTGTGCTCTTCCGATCTTCTACTATTCTTTCCCCTGCACTGT-3′) using Q5 High-Fidelity 2x Master Mix (NEB). PCR reactions were cleaned up using Agencourt AMPure XP beads (Beckman Coulter) followed by measurement of the concentration of the target amplicon using the Agilent 4200 TapeStation. DNA samples were indexed using the Nextera XT kit (Illumina). Indexed and size-selected samples were quantified using the KAPA Illumina Library Quantification kit (KAPA Biosystems) and pooled at equal concentration prior to sequencing using the Illumina NovaSeq S4 (paired-end 2×150).

### shRNA data processing and analysis

A custom reference library of the shRNA library was created for Bowtie2. Adaptor sequences were removed from the reads using Cutadapt, followed by alignment to the custom reference library using Bowtie 2. The number of reads mapping to each shRNA in each sample was then extracted from the SAM files and Mageck2^141^ was used to determine the fold-change in abundance of each shRNA between the D0 time point and later time points (D8, D17, D24) in both WT and SF3B1^K700E^. shRNA-level fold-change estimates were combined to gene-level estimates by averaging the sum of individual fold-changes of all significant shRNAs (adjusted p value < 0.05 and |FC| > 0.5) mapped to a given gene. Gene-level p values were calculated by combining the individual p values of all significant shRNAs mapping to a given gene using Fisher’s method. Gene-level weighted growth effects were calculated by multiplying the average gene-level fold change by the -log_10_ (gene-level p value). Clusterprofiler R package^142^ was used for gene ontology enrichment analysis of target gene enrichments. Analysis of over-represented gene pathways was performed using the clusterprofiler R package with the following parameters: OrgDb = ‘org.Hs.eg.db’, ont = “BP”, pvalueCutoff = 0.05, qvalueCutoff = 0.10. Pathway over-representation for screen candidates was performed using STRINGdb v11.5^85^.

### Methylcellulose colony forming unit assay

Mononuclear cells were isolated by Ficoll-Hypaque density gradient separation (Sigma Aldrich, Histopaque-1077) and cultured in methylcellulose media (Methocult GF#H4435, Stem Cell Technologies) per manufacturer’s instructions. Similarly, FACS-sorted cKit+Sca-1+Lin-cells isolated from murine bone marrow aspirates were cultured in methylcellulose median (Methocult GF#M3434, Stem Cell Technologies). The following independent experiments were performed in technical duplicates for each setting: 0.05% DMSO and 5 uM OICR-9429 (SelleckChem, S7833) in 0.05%DMSO). After 14 days of incubation, erythroid or myeloid colonies were counted, representative pictures were taken, and cultures aspirated, washed in PBS and stained with antibodies for flow cytometry. To knockdown human HDAC2 and ING2 (and Hdac2, Ing2, Wdr5 in mouse), the FACS-isolated cells (were transduced with pLKO.1-based GFP-expressing scrambled, HDAC2, ING2 (scrambled, Hdac2, Ing2, Wdr5 in mouse) -targeting lentiviral vectors by spinfection at 1800 rpm for 90 minutes at 37°C in 8 μg/mL polybrene. After 6 hours of incubation in IMDM + 10% FBS at 37°C, stably transduced cells were cultured in methylcellulose media as detailed above, and then incubated in methylcellulose medium for 14 days before colony counting and imaging. Colony counting was performed manually using STEMgrid-6 (StemCell technologies, #27000). Images of the well plates were taken using Keyence BZX-800 inverted fluorescence microscope. For the human samples, the same sequences previously validated in our shRNA rescue screen validation experiments were used to knockdown HDAC2 and ING2. Knockdown efficiency of the respective proteins was confirmed using Western Blots. The shRNA sequences used to knockdown HDAC2, ING2 in the human samples and Hdac2, Ing2, and Wdr5 in the murine LT-HSC cells are included in TableS4.

### Statistics and reproducibility

Statistical parameters including the number of biological replicates (*n*), standard error of mean (s.e.m), and statistical significance are reported in the figure legends. All results presented in this article were obtained from a minimum of three independent biological replicates, except for genome-wide analyses and DNA combing. CUT&RUN-seq for Ser2P RNAPII and H3K4me3 in human CD34+ cells was performed in a minimum of 5 biological replicates, per condition. DNA combing, K562 DRIP-seq, K562 ATAC-seq, K562 nascent SRS-seq, K562 LRS, K562 ChIP-seq, K562 GRO-seq, K562 TT-TL seq, K562 RNA-seq, K562 H3K4me3-CUT&RUN-seq, K562 H3K27me3-CUT&RUN-seq, and murine LT-HSC-CUT&RUN-seq data resulted from two independent biological replicates for each condition. U2AF1 (WT and S34F) Ser2P RNAPII ChIP-seq, H3K4me1-CUT&RUN-seq, and H3K27ac-CUT&RUN-seq, shRNA and siRNA knockdown Ser2P RNAPII ChIP-seq data came from one biological replicate per condition. K562 GRO-seq (1 replicate), H3K36me3 ChIP-seq (1 replicate) and OK-seq (three replicates) data were retrieved from external database repositories.

For Student’s *t*-test comparisons, unpaired *t*-tests were used unless otherwise specified. Data in bar graphs was presented as mean ± s.e.m. When data was presented as box-whisker plots, the Mann-Whitney U test was performed. Two sample K.S test was used to determine significance between samples in the cumulative frequency distribution plots. Test details are indicated in the figure legends. For box plots, boxes and whiskers indicate the 25th to the 75th and the 5th to the 95th percentiles, respectively, and medians are indicated. Pearson’s correlation was calculated for correlation analysis. In Venn diagrams, numbers represent genes or regions co-occurring between conditions. In screenshots from genome-wide experiments, scales are adjusted so that the background signal is low for better visualization of the results.

For IF (S9.6, γH2AX) and co-IF (γH2AX + BrdU, γH2AX + p-RPA, γH2AX + S9.6) and EdU experiments, >100 cells per replicate were measured. A minimum of 50 cells was also measured in each comet assay replicate. For DNA-combing analysis, at least 150 tracks per replicate were measured to determine fork velocity. In RNAPII NTD + PCNA PLA and PCNA + FANCD2 PLA plots, >100 total cells were represented from at least 30 cells per replicate.

Graphs were generated with the ggplot2 package of R. *P* values are indicated in the figures, and statistical tests applied are described in the figure legends. Analyzed samples were randomly chosen and data acquisition automatically performed by analysis software to ensure unbiased results.

## Supporting information

Figure5_ChromHMMdatafile

K562_MUTvsWT_4days_RNAseq

Supplementalfigures_with_legends_1-8

## Data Accession

All sequencing files used in this study are available from GEO with accession number GSE226003

## Author Contributions

PCB and MMP conceived and planned the study. PCB, AKG performed the microscopy and NGS experiments. BD and ED performed the DNA fiber combing experiments. RR, NN, HL contributed to the in vitro splicing assay experiment. BD, ED performed the DNA combing studies. AO, GK contributed to the DNA replication studies experiments. JZ, MS contributed to and supervised the bioinformatic analysis for the TT-TL-seq experiment. NC, DK, AKV provided patient samples for the study. YN and SO provided RNA-seq files of some patient samples. PCB and MMP performed bioinformatic analysis and wrote the manuscript. KMN provided input on experimental design and analysis of long read sequencing. KMN and AKV edited the manuscript. MMP provided overall supervision for the project. All the authors discussed and approved the final version of the manuscript.

## Acknowledgements

This work was funded in part by National Institutes of Health (R01 HL 133406 to MMP and KMN, R21 HL150642 to MMP and KMN, T32 CA233414 to PCB, NIDDK CCEH pilot to PCB), pilot awards from the Frederick A. DeLuca Foundation (to MMP and KMN), EvansMDS Young Investigator Award from the Edward P. Evans Foundation (to PCB). We thank the Yale Center for Genome Analysis (YCGA), Yale Center for Research Computing (YCRC), Yale Animal Resource Center (YARC), Yale Core Center of Excellence in Hematology (YCCEH, supported by NIDDK U54 DK106857) for the core resources used. Some of the patient samples used in the study were provided by The Yale Hematology Tissue bank. We thank Dr. Carl Walkley (St. Vincent’s, Melbourne, Australia) and Dr. Alex C. Minella for critical insights and discussions.

