## Supplementaltables_1-9 for "Transcription elongation defects link oncogenic splicing factor mutations to targetable alterations in chromatin landscape"

**Table S1: Top 20 positive survival candidates related to Figure 6**

| Gene | Gene p-value | Number of significant hairpins | Cumulative log2foldchange |
|---|---|---|---|
| ASXL1 | 1.00E-20 | 3 | 4.490266667 |
| AURKA | 1.00E-20 | 3 | 4.0331 |
| C2orf60 | 1.00E-20 | 3 | 4.641966667 |
| CBX6 | 1.00E-20 | 3 | 4.5981 |
| CHD8 | 1.00E-20 | 3 | 4.567633333 |
| DOT1L | 1.00E-20 | 3 | 6.452466667 |
| HDAC2 | 1.00E-20 | 3 | 4.6348 |
| HDAC3 | 1.00E-20 | 3 | 4.3838 |
| HDAC8 | 1.00E-20 | 3 | 4.384133333 |
| ING2 | 1.00E-20 | 3 | 6.3909 |
| LRWD1 | 1.00E-20 | 3 | 4.7215 |
| MORF4L1 | 1.00E-20 | 3 | 4.091766667 |
| NEK6 | 1.00E-20 | 8 | 4.063475 |
| PHF21A | 1.00E-20 | 3 | 6.336566667 |
| PHIP | 1.00E-20 | 3 | 4.903933333 |
| PRDM11 | 1.00E-20 | 3 | 4.1284 |
| PRDM9 | 1.00E-20 | 3 | 4.483566667 |
| SIRT2 | 1.00E-20 | 3 | 4.5436 |
| SIRT7 | 1.00E-20 | 3 | 4.771 |

**Table S2:Top 4 negative survival candidates related to Figure 6**

| Gene | Gene p-value | Number of significant hairpins | cumulative log2foldchange |
|---|---|---|---|
| ESCO2 | 3.25E-09 | 4 | -6.97375 |
| WDR5 | 3.77E-08 | 3 | -6.1079 |
| PYGO1 | 8.69E-06 | 3 | -4.4694 |
| KDM4D | 9.90E-07 | 3 | -5.4435 |

**Table S3: siRNAs used for protein knockdown**

| siRNA reagent | Source | Identifier |
|---|---|---|
| ON-TARGET plus Non-targeting Control | Dharmacon | D-001810-10 |
| ON-TARGET plus UAP56 | Dharmacon | L-003805-00-0005 |
| ON-TARGET plus SETX | Dharmacon | L-021420-00-0005 |
| ON-TARGET plus FANCD2 | Dharmacon | L-016376-00-0005 |
| ON-TARGET plus DDX46 | Dharmacon | L-021234-01-0005 |
| ON-TARGET plus HTATSF1 | Dharmacon | L-016645-00-0005 |

**Table S4: shRNAs used for protein knockdown**

| shRNA target | Target species | Hair-pin sequence |
|---|---|---|
| HDAC2 | Human | Replicate 1: 5’-GCAAATACTATGCTGTCAATT-3’ |
| | | Replicate 2: 5’-GCTGTGAAGTTAAACCGACAA-3’ |
| ING2 | Human | Replicate 1: 5’-CCTGTTGAGTTTGCAATAGAT-3’ |
| | | Replicate 2: 5’-CATGTGTTTCACTTACCTATA-3’ |
| PHF21A | Human | Replicate 1: 5’-CCAGAGACCTACCATTGCTAT-3’ |
| | | Replicate 2: 5’-CCTGGAACTTTAGCAATTGTT-3’ |
| Hdac2 | Mouse | 5’-CCCAATGAGTTGCCATATAAT-3’ |
| Ing2 | Mouse | 5’-CCAAGAAACGTTAAAGGAAAT-3’ |
| Wdr5 | Mouse | 5’-GCCGTTCATTTCAACCGTGAT-3’ |
| Scrambled/Control | Non-mammalian | 5’-CAACAAGATGAAGAGCACCAA-3’ |

**Table S5: Oligos used for inducible-shRNA protein knockdown**

| Target gene | Replicate | Primer orientation | Oligo sequence |
|---|---|---|---|
| HTATSF1 | 1 | Forward | 5’CCGGCGCATCTAGTTCTACCGCAAACTCGAGTTTGCGGTAGAACTAGATGCGTTTTTG-3’ |
| | | Reverse | 5’AATTCAAAAACGCATCTAGTTCTACCGCAAACTCGAGTTTGCGGTAGAACTAGATGCG-3’ |
| | 2 | Forward | 5’CCGGGCAACTGGAATGGCGTTTGAACTCGAGTTCAAACGCCATTCCAGTTGCTTTTTG-3’ |
| | | Reverse | 5’AATTCAAAAAGCAACTGGAATGGCGTTTGAACTCGAGTTCAAACGCCATTCCAGTTGC-3’ |
| DDX46 | 1 | Forward | 5’CCGGGCAGAAATCACCAGGCTCATACTCGAGTATGAGCCTGGTGATTTCTGCTTTTTG-3’ |
| | | Reverse | 5’AATTCAAAAAGCAGAAATCACCAGGCTCATACTCGAGTATGAGCCTGGTGATTTCTGC-3’ |
| | 2 | Forward | 5’CCGGGAAGAAGTGAAAGAGGAAGTACTCGAGTACTTCCTCTTTCACTTCTTCTTTTTG-3’ |
| | | Reverse | 5’AATTCAAAAAGAAGAAGTGAAAGAGGAAGTACTCGAGTACTTCCTCTTTCACTTCTTC-3’ |

**Table S6: Primers used in nascent long-read sequencing**

| Primer | Sequence |
|---|---|
| DNA adapter | 5’-/5rApp/NNNNNCTGTAGGCACCATCAAT/3ddC/-3’ |
| Custom Reverse transcriptase primer (WT) | 5’AAGCAGTGGTATCAACGCAGAGTACCACATATCAGAGTGCGGATTGATGGTGCCTACAG-3’ |

| | |
|---|---|
| Custom Reverse transcriptase primer (SF3B1K700E) | 5’AAGCAGTGGTATCAACGCAGAGTACACACACAGACTGTGAGGATTGATGGTGCCTACAG-3’ |
| Template switching oligo | 5’-AAGCAGTGGTATCAACGCAGAGTACrGrG+G-3’ |

**Table S7: Primers used for qRT-PCRs in gene expression, DRIP-seq, and TT-TL-seq analyses**

| **DNA primer** | |
|---|---|
| **Gene expression** | |
| DDX46#1 Fwd | 5’-CCAAGCTCAATTATGTGCCGT-3’ |
| DDX46#1 Rev | 5’-AGCTTCCTTAGAGGTAACTTTCCA-3’ |
| DDX46#2 Fwd | 5’-AAAGGAAAAGGTTGCCCCAAAC-3’ |
| DDX46#2 Rev | 5’-GATGGGCGTGGGCTTTTCATA-3’ |
| hTATSF1#1 Fwd | 5’-ATGGTTTCATGTTGAAGAAGACAG-3’ |
| hTATSF1#1 Rev | 5’-TGCCAAACTTGGACATAAGTTG-3’ |
| hTATSF1#2 Fwd | 5’-CATGTTGAGGTGGCAAAGTTTCA-3’ |
| hTATSF1#2 Rev | 5’-CCTCTCAGGTCTCCAATCCAA-3’ |
| hTATSF1#3 Fwd | 5’-TTGGTGGCCGTCAAATCACT-3’ |
| hTATSF1#3 Rev | 5’-AGGAAAGCCTCCCATCCTCT-3’ |
| B-Actin Fwd | 5’-ATCCACGAAACTACCTTCAACTC-3’ |
| B-Actin Rev | 5’-GAGGAGCAATGATCTTGATCTTC-3’ |
| **DRIP-seq** | |
| EGR1 Fwd | 5’-GAACGTTCAGCCTCGTTCTC-3’ |
| EGR1 Rev | 5’-GGAAGGTGGAAGGAAACACA-3’ |
| snRPN Fwd | 5’-GCCAAATGAGTGAGGATGGT-3’ |
| snRPN Rev | 5’-TCCTCTCTGCCTGACTCCAT-3’ |
| TFPT Fwd | 5’-TCTGGGAGTCCAAGCAGACT-3’ |
| TFPT Rev | 5’-AAGGAGCCACTGAAGGGTTT-3’ |
| RPL13A Fwd | 5’-AGGTGCCTTGCTCACAGAGT-3’ |
| RPL13A Rev | 5’-GGTTGCATTGCCCTCATTAC-3’ |
| **TT-TL seq** | |
| MYC_premRNA Fwd | 5’-TGTAACCTTGCTAAAGGAGTGAT-3’ |
| MYC_premRNA Rev | 5’-GAGCCTGCCTCTTTTCCACA-3’ |
| MYC_matmRNA Fwd | 5’-CTCGACGGAGTCCTCCCC-3’ |
| MYC_matmRNA Rev | 5’-GAGCCTGCCTCTTTTCCACA-3’ |
| MIR17HG_premRNA Fwd | 5’-CGGTCGTAGTAAAGCGCAGG-3’ |
| MIR17HG_premRNA Rev | 5’-ACAACAGGTTTCCCTCCGTC-3’ |
| MIR17HG_matmRNA Fwd | 5’-CGGTCGTAGTAAAGCGCAGG-3’ |
| MIR17HG_matmRNA Rev | 5’-TTCTCAGAAAAGCTACTGGTGC-3’ |
| CDKN1B_premRNA Fwd | 5’-GTCTTAGGTGTTCAGTGCTACC-3’ |
| CDKN1B_premRNA Rev | 5’-GAGGGACCGCGATGTATTAAG-3’ |
| CDKN1B_matmRNA Fwd | 5’-GGCAAAAATCCGAGGTGCTT-3’ |
| CDKN1B_matmRNA Rev | 5’-TGTGTTTACACAGCCCGAAG-3’ |
| JUN Fwd | 5’-CCTTGAAAGCTCAGAACTCGGAG-3’ |
| JUN Rev | 5’-TGCTGCGTTAGCATGAGTTGGC-3’ |

**Table S8: Antibodies used in the study**

| Antibody | Species | Company | Catalogue |
|---|---|---|---|

| | | | |
|---|---|---|---|
| Anti-phospho-Histone H2AX (Ser139), clone JBW301 | Mouse | EMB millipore | 05-636 |
| Anti-phospho-Histone H2AX (Ser139) (20E3) polyclonal antibody | Rabbit | Cell Signaling Technologies | 9718S |
| Anti-Histone H2AX (D17A3) polyclonal antibody | Rabbit | Cell Signaling Technologies | 7361S |
| Monoclonal S9.6 antibody | Mouse | Hybridoma HB-8730 | n/a |
| Polyclonal anti-Nucleolin antibody | Rabbit | Abcam | ab50279 |
| Polyclonal anti-phospho RPA32 (S33) antibody | Rabbit | Abcam | Ab211877 |
| Monoclonal anti-BrdU antibody | Mouse | Santa Cruz Biotechnology | sc-70411 |
| Polyclonal anti-Senataxin antibody | Rabbit | Abcam | ab220827 |
| Polyclonal FANCD2 antibody | Rabbit | Bethyl | A302-174A |
| Monoclonal FANCD2 antibody | Mouse | Santa Cruz | sc-20022 |
| Polyclonal UAP56/DDX39B antibody | Rabbit | Protein tech | 14798-1-AP |
| Polyclonal anti-ATR antibody | Rabbit | Bethyl | A300-137A |
| Polyclonal anti-phospho-ATR antibody | Rabbit | Cell signaling technologies | 2853T |
| Polyclonal anti-Chk1 antibody | Rabbit | Bethyl | A300-298A |
| Polyclonal anti-Chk2 antibody | Rabbit | Bethyl | A300-618A |
| Polyclonal anti-phospho-Chk1 antibody (Ser345) | Rabbit | Cell signaling technology | 2348 |
| Polyclonal anti-phospho-Chk2 antibody (Thr68) | Rabbit | Cell signaling technology | 2661 |
| Polyclonal anti-GAPDH (14C10) antibody | Rabbit | Cell signaling technology | 2118 |
| Polyclonal anti-PCNA antibody | Mouse | Cell signaling technology | 2586 |
| Polyclonal anti-Beta-Actin antibody | Rabbit | Cell signaling technology | 4967 |
| Monoclonal anti-BrdU (B44) | Mouse | BD Biosciences | 347580 |
| Monoclonal anti-BrdU [BU1/75 (ICR1)] | Rat | Abcam | ab6326 |
| Polyclonal anti-Rpb1 NTD (D8L4Y) antibody | Rabbit | Cell signaling technology | 14958S |
| Polyclonal anti-Phospho-Rpb1 CTD (Ser2) (E1Z3G) antibody | Rabbit | Cell signaling technology | 13499S |
| Polyclonal anti-CDK9 (C12F7) antibody | Rabbit | Cell signaling technology | 2316 |
| Polyclonal anti-Drosophila Histone H2Av antibody | Rabbit | Active Motif | 39715 |
| Polyclonal anti-Tri-methyl-Histone 3 (Lys4) (C42D8) antibody | Rabbit | Cell signaling technology | 9731 |
| Polyclonal anti-Mono-methyl-Histone 3 (Lys4) (D1A9) antibody | Rabbit | Cell signaling technology | 5326 |

| | | | |
|---|---|---|---|
| Polyclonal anti- Tri-methyl-Histone 3 (Lys4) (C36B11) antibody | Rabbit | Cell signaling technology | 9733 |
| Polyclonal anti-Acetyl-Histone 3 (Lys27) (D5E4) antibody | Rabbit | Cell signaling technology | 8173 |
| Polyclonal Normal IgG antibody | Rabbit | Cell signaling technology | 2729 |
| Polyclonal anti-HDAC2 (D6S5P) antibody | Rabbit | Cell signaling technology | 57156 |
| Polyclonal anti-PHF21A antibody | Rabbit | Bethyl | A303-603 |
| Polyclonal anti-ING2 antibody | Rabbit | Protein tech | 11560-1-AP |
| Polyclonal anti-WDR5 antibody | Rabbit | Bethyl | A302-430A |
| Monoclonal anti-DYKDDDDK (FLAG) (D6W5B) antibody | Rabbit | Cell signaling technology | 14793S |
| Polyclonal anti-Histone H3 antibody | Rabbit | Cell signaling technology | 9715S |
| Monoclonal anti-HA-tag (C29F4) antibody | Rabbit | Cell signaling technology | 3724S |
| Polyclonal anti-HTATSF1 antibody | Rabbit | Protein tech | 20805-1-AP |
| Polyclonal anti-SF3A3 antibody | Rabbit | Bethyl | A302-506A |
| Polyclonal anti-U2AF1 (D6S3Q) antibody | Rabbit | Cell signaling technology | 13705S |
| Polyclonal anti-MFAP1 antibody | Rabbit | Bethyl | A304-647A |
| Polyclonal anti-SF1 antibody | Rabbit | Bethyl | A303-214A |
| Polyclonal anti-SNRP70/U1-70K antibody | Rabbit | Abcam | ab83306 |
| Monoclonal anti-SF3B1 [EPR11987(B)] antibody | Rabbit | Abcam | ab170854 |
| Polyclonal anti-U2AF2 antibody | Rabbit | Bethyl | A303-666A |
| Polyclonal anti-SUGP1 antibody | Rabbit | Bethyl | A304-675A |
| Polyclonal anti-DDX42 antibody | Rabbit | Bethyl | A303-354A |
| Polyclonal anti-DDX46 antibody | Rabbit | Bethyl | A301-052A |
| Flow antibodies: | | | |
| PE/Cynanine7 anti-mouse/human CD11b antibody, clone M1/70 | Rat | Biolegend | 101215 |
| APC anti-mouse TER-119/Erythroid cells, clone TER-119 | Rat | Biolegend | 116211 |
| APC anti-mouse Lineage cocktail | | BD Biosciences | 558074 |
| PE/Cyanine7 anti-mouse CD117 (c-kit) antibody, clone 2B8 | Rat | Biolegend | 105813 |
| FITC anti-mouse Ly-6A/E (Sca1) antibody | Rat | Biolegend | 108105 |
| PE/Cyanine7 anti-human CD71 antibody, clone CY1G4 | Mouse | Biolegend | 334111 |
| APC anti-human CD11b antibody, clone CBRM1/5 | Mouse | Biolegend | 301409 |
| FITC anti-human CD34 antibody, clone 581 | Mouse | Biolegend | 343503 |
| Secondary antibodies: | | | |

| | | | |
|---|---|---|---|
| anti-rat Alexa Fluor 488 | Goat | Invitrogen | A11006 |
| Anti-mouse Cy3 | Sheep | Sigma | C2181 |
| anti-rat Alexa Fluor Plus 488 | Goat | Invitrogen | A48262TR |
| Anti-mouse Cy3 | Goat | Abcam | Ab97035 |
| Anti-rabbit IgG, HRP-linked antibody | Goat | Cell signaling technologies | 7074S |
| Anti-mouse IgG, HRP-linked antibody | Horse | Cell signaling technologies | 7076S |

**Table S9: Reagents used in the study**

| **Chemical name** | **Company** | **Catalogue number** |
|---|---|---|
| RPMI1640 | Gibco | 11875-093 |
| Fetal Bovine Serum | Gibco | A3840001 |
| Penicillin Streptomycin Solution 50x | Corning | 30-001-CI |
| DMEM | Gibco | 11965-092 |
| Schneider's modified *Drosophila* medium | Lonza | 04-351Q |
| Doxycycline | Sigma | D9891-1G |
| Puromycin | Invitrogen | A1113802 |
| Blasticidin | Invitrogen | A1113903 |
| Hygromycin B | Invitrogen | 10687010 |
| Neomycin Trisulfate | Sigma | N1876-25G |
| PEG-8000 | Sigma | 1546605 |
| Polybrene | Yeasen | 40804ES76 |
| TransIT-X2 | Mirus Bio | MIR 6003 |
| Protease inhbitor coctail | Millipore Sigma | 11836170001 |
| phenylmethylsulfonyl fluoride | Sigma | P-7626 |
| Tris-HCl pH 7.4 | American Bioanalytical | AB14044-01000 |
| NP40 | Calbio Chem | 492015 |
| Sodium Choride Solution, 5M | Sigma-Aldrich | S5150-1L |
| Sucrose, Crystal | J.T.Baker | 4072-01 |
| PBS, pH 7.4 | American Bioanalytical | AB11072-01000 |
| 0.5M EDTA | LIFE TECHNOLOGIES | 15575-038 |
| MgCl2 | Sigma | M-8266 |
| ANTI-FLAG® M2 affinity gel | Sigma-Aldrich | |
| FLAG peptide | Millipore Sigma | F4799 |
| Laemmli SDS sample buffer | Thermofisher | J60015.AD |
| Protease inhibitor cocktail tablets | Sigma | 11873580001 |

| | | |
|---|---|---|
| Phosphatase inhibitor cocktail tablets | Roche | 04906837001 |
| Dithiothreitol, DTT | SIGMA | D-5545 |
| Glycerol | J.T.Baker | 2136-01 |
| HEPES 1M | American Bioanalytical | AB06021 |
| Nocodazole | Sigma-Aldrich | M1404-2MG |
| 5-Ethynyl 2'-deoxyuridine | Thermofisher Scientific | A10044 |
| 5-Ethynyl Uridine | Thermofisher Scientific | E10345 |
| Thymidine | Sigma | T-1895 |
| CytoPhase Violet | Biolegend | 425701 |
| Bovine Serum Albumin | Sigma-Aldrich | A9647-100G |
| ProLong Gold AntiFade reagent | Cell Signaling Technology | 8961S |
| SYBRsafe | Invitrogen | S33102 |
| Glass coverslips coated with Poly-L-lysine | Millipore Sigma | P0425-72EA |
| Paraformaldehyde | Sigma | P-6148 |
| Triton-X100 | MP Biomedicals | 194854 |
| Phenol solution | Sigma-Aldrich | P4557-100ML |
| Chloroform:isoamyl alcohol 24:1 | Sigma | C0549-1PT |
| Short cut RNaseIII | NEB | M0245S |
| RNaseH1 | NEB | M0297S |
| Methylene blue | MCIB | 742341 |
| Tween-20 | Bio-rad | 1706531 |
| Nylon membrane | Amersham Biosciences | RPN203B |
| Formaldehyde 16% concentrate stock methanol-free | Cell Signaling Technology | 12606P |
| Glycine | Sigma | G-7403 |
| Protease inhibitor | Millipore Sigma | 11836170001 |
| Pipes pH 8 | Fisher | BP304 |
| KCl | Sigma | P-4504 |
| Spermine | Sigma | S3256 |
| Spermidine | Sigma-Aldrich | 85558 |
| MNase | NEB | M0247S |
| 20% SDS Solution | American bio | AB01922-00500 |
| Pierce Protein A/G Magnetic beads | ThermoScientific | 78609 |
| Pierce Protein A/G Agarose beads | ThermoScientific | 20421 |
| LiCl | Sigma | L7026-500ML |
| Proteinase K | Roche | 12570000 |
| phenol:chloroform:isoamyl alcohol 25:24:1 | Sigma-Aldrich | P2069 |

| | | |
|---|---|---|
| Sodium Acetate, 3M, pH 5.2 | American bio | AB13168-01000 |
| Ethyl alcohol | Sigma-Aldrich | E7023-500ML |
| CaCl2 | Sigma | C-4901 |
| Superase-in | Invitrogen | AM2694 |
| IGEPAL CA-630 | Sigma | I8896-50ML |
| ATP | NEB | N0450S |
| GTP | NEB | N0450S |
| 4-thio-UTP | Trilink Biotechnologies | N-1025-1 |
| CTP | NEB | N0450S |
| Sarkosyl (N-Laurylsarcosine sodium salt solution) | Sigma | 61747 |
| TRIzol Reagent | ambion | 15596018 |
| Dimethylformamide | Sigma | D-4254 |
| EZ-link HPDP Biotin | ThermoFisher Scientific | 21341 |
| 4-thio-uridine | Sigma | T4509-25MG |
| M280 Streptavidin Dynabeads | Invitrogen | 11205D |
| TURBO DNase | ThermoFisher Scientific | AM2238 |
| Dynabeads MyOne Streptavidin C1 magnetic beads | ThermoFisher Scientific | 65001 |
| Diethylpyrocarboate, DEPC | American bio | AB00472-00025 |
| RNAcleanXP beads | Beckman Coulter | A63987 |
| 2,2,2-Trifluoroethylamine | Sigma-Aldrich | 269042-1G |
| DEPC-treated water | Invitrogen | 750023 |
| Concanavalin A-coated magnetic beads | Cell signaling technologies | 93569 |
| Manganese Chloride | Fisher | M87-500 |
| Digitonin | Dig-Wash | MP215948082 |
| pAG/MNase | EpiCypher | 151016 |
| EGTA | Sigma | E-3889 |
| RNase A | Thermofisher Scientific | R1253 |
| Glycogen | Thermofisher Scientific | R0551 |
| E.coli spike in control | Epicypher | 18-1401 |
| α-amanitin | Cayman Chemicals | 17898 |
| Urea | American Bioanalytical | AB02100 |
| AMPure XP beads | Beckman Coulter | NC9933872 |
| $NaPO_4$ | Sigma | S-0751 |
| Agarose Protein-A/G beads | ThermoScientific | 20421 |
| Q5 High-Fidelity 2x Master Mix | NEB | |

| | | |
|---|---|---|
| Agencourt AMPure XP beads | Beckman Coulter | |
| Ficoll-Hypaque density gradient separation | Sigma Aldrich | Histopaque-1077 |
| Methylcellulose media (Human) | Stem Cell Technologies | Methocult GF#H4435 |
| Methylcellulose media (Mouse) | Stem Cell Technologies | Methoccult GF#M3434 |
| 5 ìM OICR-9429 | SelleckChem | S7833 |
| IMDM | Gibco | 12440053 |
| STEMgrid-6 | StemCell technologies | 27000 |
| TRIzol LS reagent | Invitrogen | 10296028 |
| MinElute PCR purification kit | Qiagen | 28004 |
| NEBNext® Ultra II DNA Library Prep Kit | NEB | E7645S |
| RNA Clean and Purification kit | Zymo Research | R1013 |
| SMARTer® Universal Low Input RNA Kit | Takara | 634938 |
| Qiagen RNeasy Mini Kit | Qiagen | 74104 |
| SMARTer PCR cDNA Synthesis Kit | Clontech | 634928 |
| Advantage 2 PCR Kit | Clontech | 639207 |
| Magnetic mRNA isolation Kit | NEB | S1550S |
| Click IT chemistry Kit | Thermofisher Scientific | C10640 |
| Duolink PLA technology | Merck | DUO92007-30RXN DUO92014-30RXN |
