## Supplementalfigures_with_legends_1-8 for "Transcription elongation defects link oncogenic splicing factor mutations to targetable alterations in chromatin landscape"

### SUPPLEMENTAL FIGURE 1

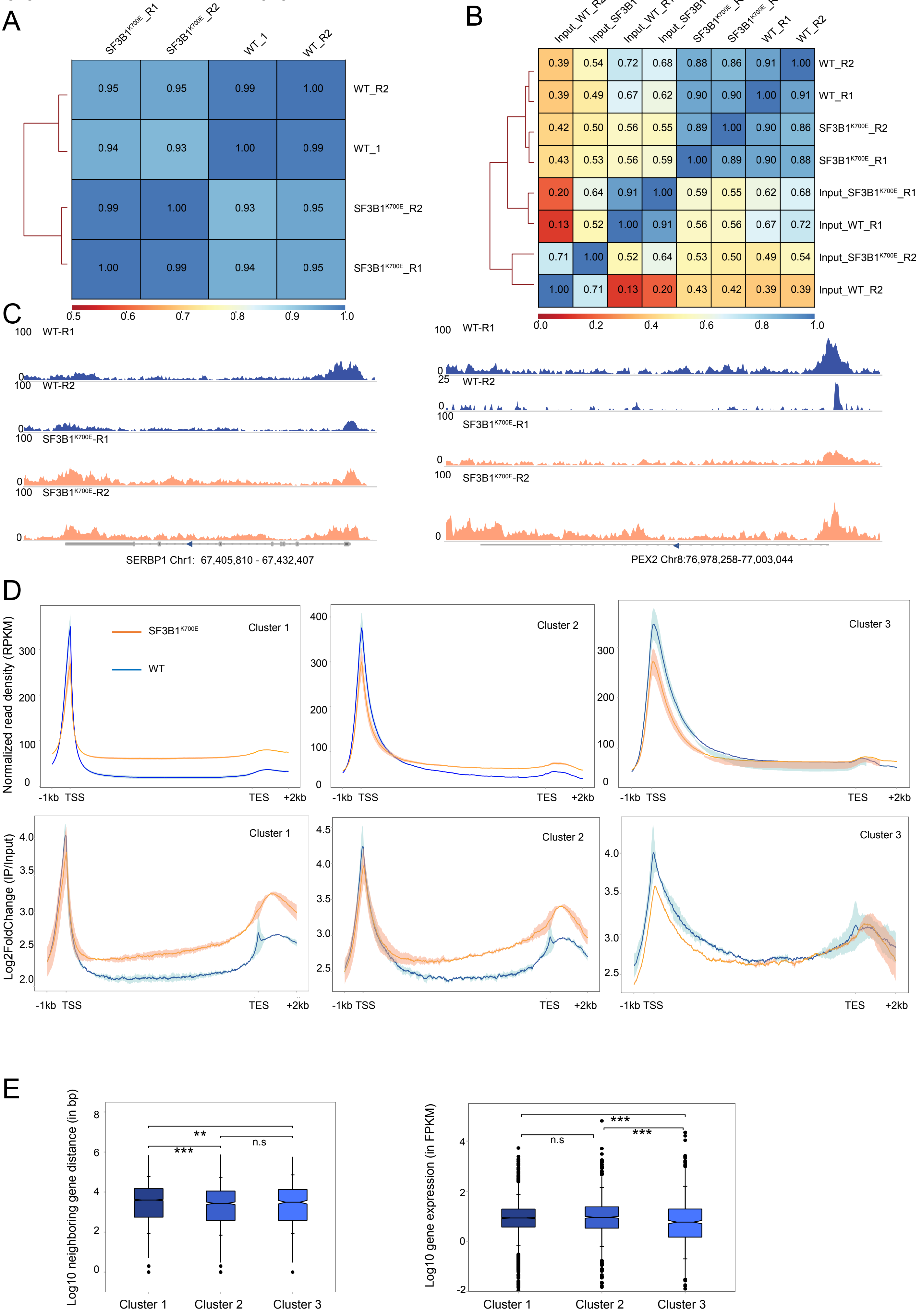


| | SF3B1K700E_R1 | SF3B1K700E_R2 | WT_1 | WT_R2 | |
|---|---|---|---|---|---|
| | 0.95 | 0.95 | 0.99 | 1.00 | WT_R2 |
| | 0.94 | 0.93 | 1.00 | 0.99 | WT_1 |
| | 0.99 | 1.00 | 0.93 | 0.95 | SF3B1K700E_R2 |
| | 1.00 | 0.99 | 0.94 | 0.95 | SF3B1K700E_R1 |

| Input_WT_R2 | Input_SF3B1K700E_R2 | Input_WT_R1 | Input_SF3B1K700E_R1 | SF3B1K700E_R1 | SF3B1K700E_R2 | WT_R1 | WT_R2 | |
|---|---|---|---|---|---|---|---|---|
| 0.39 | 0.54 | 0.72 | 0.68 | 0.88 | 0.86 | 0.91 | 1.00 | WT_R2 |
| 0.39 | 0.49 | 0.67 | 0.62 | 0.90 | 0.90 | 1.00 | 0.91 | WT_R1 |
| 0.42 | 0.50 | 0.56 | 0.55 | 0.89 | 1.00 | 0.90 | 0.86 | SF3B1K700E_R2 |
| 0.43 | 0.53 | 0.56 | 0.59 | 1.00 | 0.89 | 0.90 | 0.88 | SF3B1K700E_R1 |
| 0.20 | 0.64 | 0.91 | 1.00 | 0.59 | 0.55 | 0.62 | 0.68 | Input_SF3B1K700E_R1 |
| 0.13 | 0.52 | 1.00 | 0.91 | 0.56 | 0.56 | 0.67 | 0.72 | Input_WT_R1 |
| 0.71 | 1.00 | 0.52 | 0.64 | 0.53 | 0.50 | 0.49 | 0.54 | Input_SF3B1K700E_R2 |
| 1.00 | 0.71 | 0.13 | 0.20 | 0.43 | 0.42 | 0.39 | 0.39 | Input_WT_R2 |

### SUPPLEMENTAL FIGURE 1

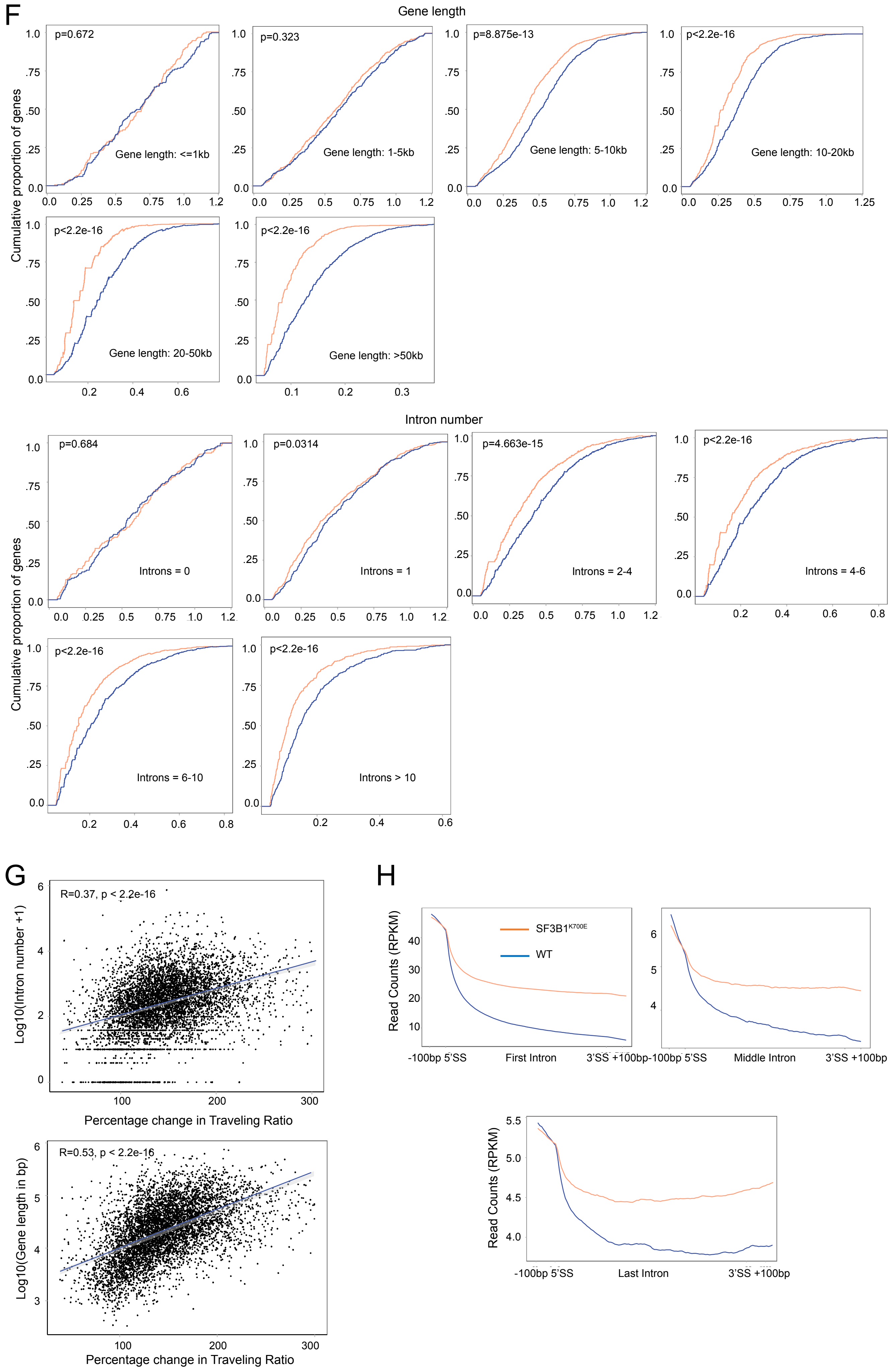

Supplemental Figure 1: **Genome-wide analysis of RNAPII redistribution due to SF3B1$^{K700E}$** (A) Correlation between RNAP II NTD ChIP-seq samples. Values in boxes represent Pearson's correlation coefficients between corresponding samples (WT and SF3B1$^{K700E}$ K562, *n* = 2 biological replicates each). (B) Values in boxes represent Pearson's correlation coefficients between corresponding samples (WT and SF3B1$^{K700E}$ K562, *n* = 2 biological replicates each). (C) Individual Ser2P RNAPII ChIP-seq gene tracks at *PEX2 and SERBP1*— in K562 cells WT versus SF3B1$^{K700E}$, at 4 days post doxycycline induction. The x-axis indicates the chromosome position, and the y-axis represents normalized read density in RPKM. (D) Meta plots comparing the RNAPII NTD ChIP signal (Top panel) $\log_2$ fold change signal of RNAPII Ser2P ChIP-seq IP/input (Bottom panel) between WT and SF3B1$^{K700E}$ K562 cells, for the 3 gene clusters (n=2 biological replicates, each). Solid lines represent the averaged signal, and the shaded area represents the upper and lower signal limits of the individual replicates (n =2, WT; n =2, SF3B1$^{K700E}$). Read densities are calculated as RPKM. (E) Box-whisker plots showing the distribution of neighboring gene distance (left) and gene expression in FPKM (right) for the three gene clusters. Dots represent data for individual genes (two-tailed Mann–Whitney *U*-test. $^{***}P = 0.0001$, $^{*}P = 0.02$, $^{ns}P > 0.05$). Neighboring gene distance and gene expression have been log-transformed for better visualization. (F) Top panel: The cumulative distribution function (CDF) plot of the TR distribution (calculated as an average of two replicates) in WT and SF3B1$^{K700E}$ cells depending on gene length. Gene lengths- <=1kb: 215 genes, 1-5kb: 599 genes, 5-10kb: 823 genes, 10-20kb: 1078 genes, 20-50kb:1638 genes, >50kb: 1480 genes. Bottom panel: The CDF plot of the TR distribution (calculated as an average of two replicates) in WT and SF3B1$^{K700E}$ cells depending on gene intron content/number. Intronless: 191 genes, single intron: 466 genes, 2-4 introns: 1503 genes, 4-6 introns:1537 genes, 6-10 introns: 1485 genes, >10 introns: 597 genes. (G) Top: Pearson correlation scatter plot comparing gene length (log(10) transformed gene length) and percentage change in traveling ratio ($WT_{TR}$-SF3B1$^{K700E}{}_{TR}$/$WT_{TR}$). Bottom: Pearson correlation scatter plot comparing gene intron number ((log(2)transformed intron number )+ 1) and percentage change in traveling ratio ($WT_{TR}$-SF3B1$^{K700E}{}_{TR}$/$WT_{TR}$). (H) Read coverage in the intronic region of interest (From left to right: first, middle, and final introns; n =5118, each) in multi-intronic genes (greater than 2 introns) from the 6694 gene dataset. Coverage is normalized to the position 100 nt upstream of each 5'SS. Orange line indicates SF3B1$^{K700E}$ and the blue line indicates WT.

### SUPPLEMENTAL FIGURE 2

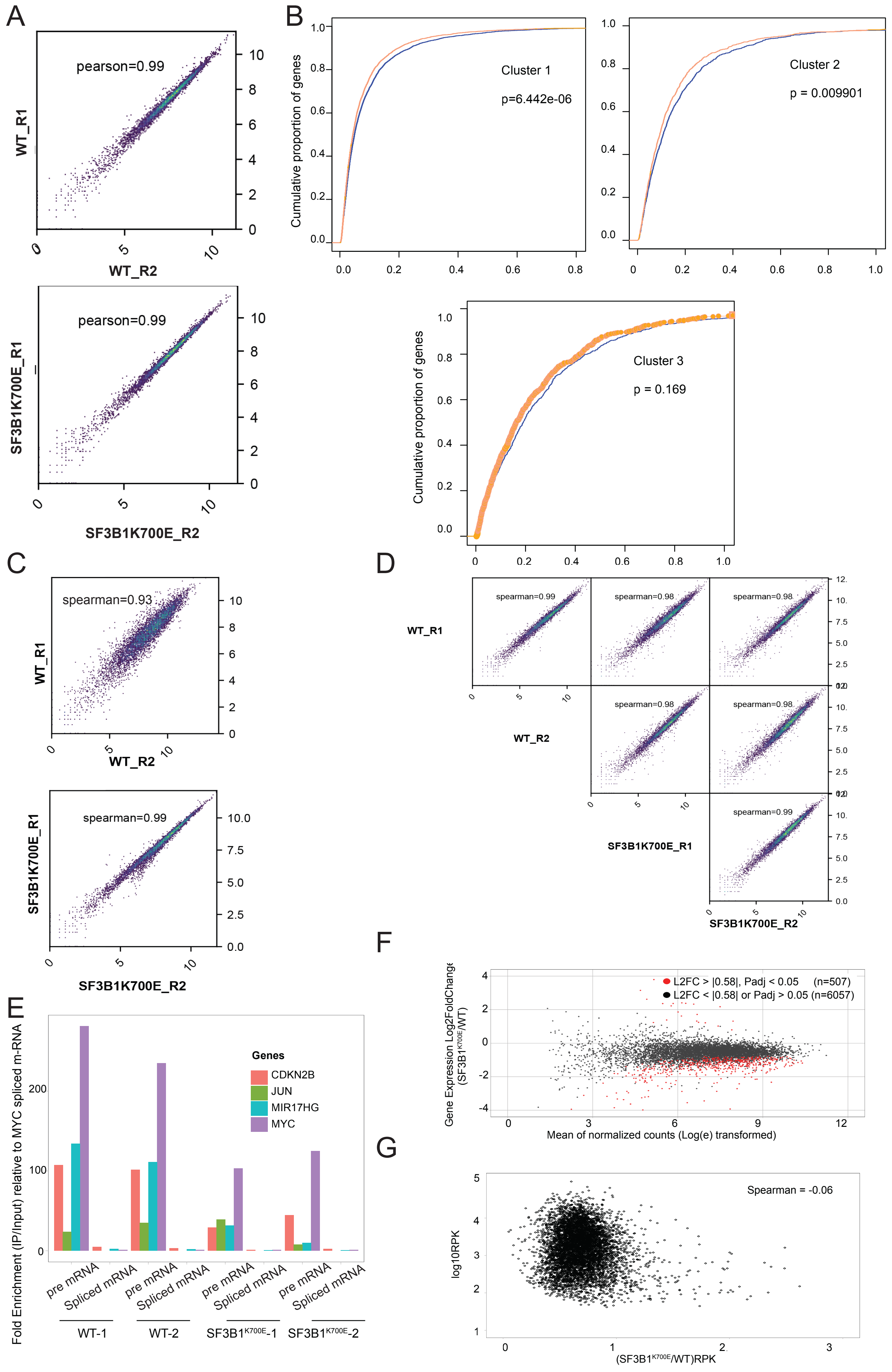

#### SUPPLEMENTAL FIGURE 2

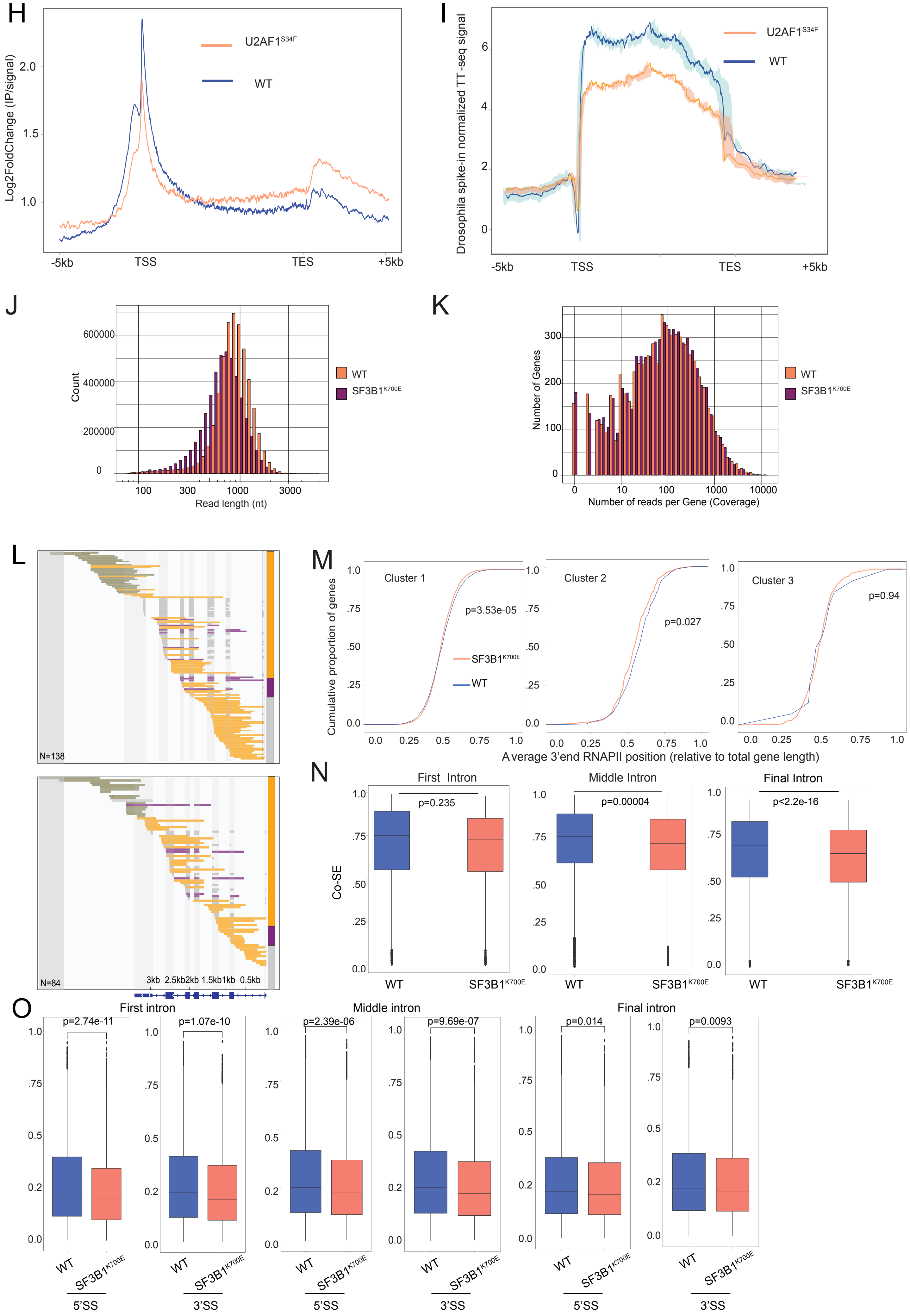

Supplemental Figure 2: **Determination SF3B1$^{K700E}$ induced changes in RNAPII transcription kinetics using nascent transcriptome assays** (A) Scatter plots depicting correlation between GRO-seq samples in WT and SF3B1$^{K700E}$ cells (n=2 replicates, each). Values in boxes represent Pearson's correlation coefficients between corresponding samples ($n$ = 2 biological replicates). (WT, r=0.986; SF3B1$^{K700E}$, r=0.991). Values expressed Log2(gene-expression + 1). (B) CDF plots of TR from GRO-seq for 'ChIP-seq-derived' maximally affected (Cluster I), moderately affected (Cluster 2) and less affected (Cluster 3) genes in WT (blue) versus SF3B1$^{K700E}$ K562 (orange) cells. Cluster I = 4289 genes, Cluster 2 = 1317 genes, and cluster 3 = 820 genes. (C) Scatter plots depicting correlation between TT-TL seq samples in WT (n=2) and SF3B1$^{K700E}$ (n=2). Values in boxes represent Spearman's correlation coefficients between corresponding samples ($n$ = 2 biological replicates). (WT, spearman=0.93; SF3B1$^{K700E}$, spearman=0.99). Values expressed Log2(gene-expression + 1). (D) Scatter plots depicting correlation between nascent short read RNA-seq samples in WT and SF3B1$^{K700E}$ cells. Values in boxes represent Spearman's correlation coefficients between corresponding samples ($n$ = 2 biological replicates). Values expressed Log2(gene-expression + 1). (E) TT-seq RT–qPCR analyses showing relative fold enrichment of pre-mRNA (intron-exon spanning primers) and spliced/mature mRNA (exon-exon primers) for *CDKN2B, MYC, MIR17HG,* and *JUN* (intronless) genes in WT and SF3B1$^{K700E}$ cells, 4 days post doxycycline induction. MYC spliced/mature mRNA (IP/input) was set as 1. WT ($n$ = 2; WT-R1 and WT-R2) and SF3B1$^{K700E}$ (n=2; SF3B1$^{K700E}$-R1 and SF3B1$^{K700E}$-R2). Data plotted are for the mean of two technical replicates per condition. (F) Minus-average plot plotting the log2 fold change (n = 6694 genes) on the y axis and log(e)-transformed normalized mean counts on the x axis. Significantly upregulated and downregulated genes (L2FC > |0.5|, $P_{adj}$<0.05) in SF3B1$^{K700E}$ (vs WT) are shown in red, above and below the zero line of the y axis, respectively. (G) Scatter plot comparing $WT_{TT\text{-}seq}$ RPK (Log-transformed; plotted on Y axis) and ratio of $SF3B1^{K700E}{}_{TT\text{-}seq}$ to $WT_{TT\text{-}seq}$ RPK (plotted on X axis). Spearman correlation of -0.06. (H) Meta plot showing the $log_2$ fold change signal of RNAPII Ser2P ChIP-seq IP data (normalized to input) across the annotated gene transcripts genes (n = 6694 genes) in WT (blue) and U2AF1$^{S34F}$ (orange) K562 cells (n =1, WT; n =1, U2AF1$^{S34F}$). Read densities are calculated as RPKM. Abbreviations used: TSS-transcription start site, TES-transcription end site (I) Metaplot showing Drosophila spike-in normalized TT-seq densities in WT (blue) and U2AF1-mutant (orange) K562 cells (TSS -5kb to TES +5kb regions) (n = 6694 genes, 2 biological replicates each). Solid lines represent the averaged signal, and the shaded area represents upper and lower signal limits among the individual replicates. (J) Read length distribution of PacBio long-reads. (K) Read depth distribution of PacBio long-reads. (L) LRS data are shown for WT (top) and SF3B1$^{K700E}$ (bottom) K562 cells for the *FTL* gene. Reads are aligned to the genome and ordered by 3′ end positions. Each horizontal row represents one read. Each transcript was categorized and colored according to its splicing status, which can be either completely spliced (grey), partially spliced (purple), all unspliced (orange), or "NA" (green) (i.e., transcripts that did not span an entire intron or a 3′SS). 'n' is the total number of reads aligning to the selected area of the genome for a particular condition. Bar graphs at the far left (for both the upper and lower plots) indicate the fraction of the aligned intron-spanning reads that are partially spliced (purple), unspliced (orange), fully spliced (grey) for that gene. (M) The CDF plot of the average 3’end RNAPII position (as determined by the LRS 3’read ends) in WT and SF3B1$^{K700E}$ in the 3 RNAPII ChIP-based gene clusters, shown in Figure 3E. Significance determined using two-sample K.S. test. (N) CoSE for introns in SF3B1$^{K700E}$ compared to the CoSE for introns in WT separated by the position of intron (first, middle, and last introns from left to right) in the selected 5118 isoforms. Analysis only included introns covered by at least 10 reads. Significance tested by Mann Whitney U test. (O) Ratio of spliced reads over total unspliced and spliced reads in WT and SF3B1$^{K700E}$ in GRO-seq at the 5’SS and 3’SS. All the introns for the first, middle, and last (from left to right) in the selected 5118 isoforms were considered. SS-splice sites. n=2 biologically independent experiments.

### SUPPLEMENTAL FIGURE 3

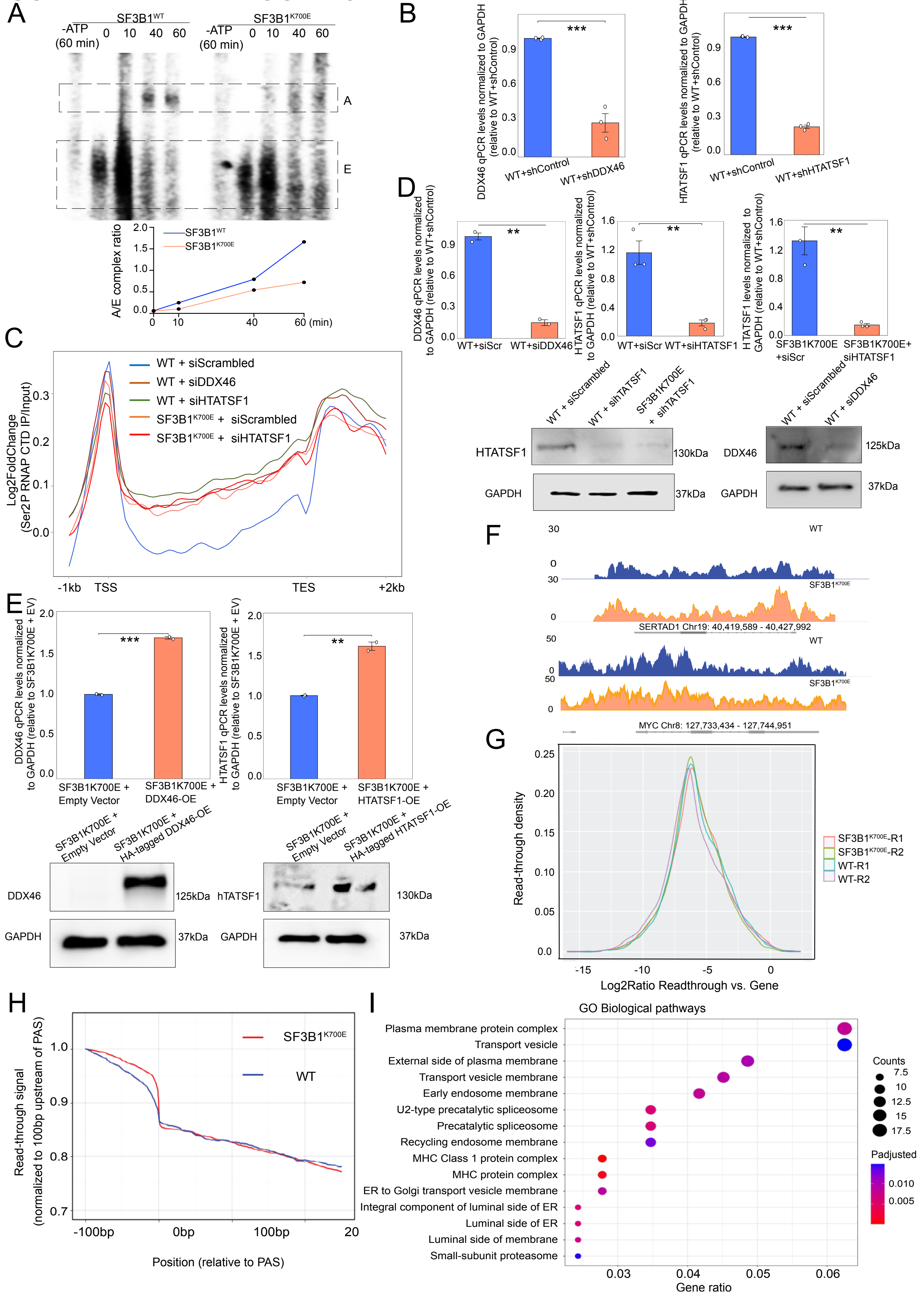

Supplemental figure 3: **Mechanisms underlying SF3B1$^{K700E}$ induced transcription elongation defect** (A) WT K562 cells transduced with inducible shControl (shScrambled), shDDX46, or shHTATSF1 were subjected to RT q-PCR to confirm knockdown of DDX46 (left) and HTATSF1 (right). qPCR levels normalized to housekeeping gene, GAPDH. WT+shControl levels set as 1. Data plotted are for the mean of three technical replicates. (B) Top panel: WT K562 cells electroporated with siRNA targeting control (siScr), DDX46 (siDDX46), or HTATSF1 (siHTATSF1), and SF3B1$^{K700E}$ K562 cells electroporated with either siRNA targeting control (siScr), or HTATSF1 (siHTATSF1). Cell lysates subjected to RT q-PCR to confirm knockdown of DDX46 and HTATSF1 in WT, and HTATSF1 in SF3B1$^{K700E}$. qPCR levels normalized to housekeeping gene, GAPDH. siScr levels set as 1. Data plotted are for the mean of two technical replicates (for DDX46) and 3 technical replicates (for HTATSF1). Bottom panel: Western blot analysis confirming knockdown of HTATSF1 (in siHTATSF1-electroporated cells) and DDX46 (in siDDX46-electroporated cells) with antibodies against DDX46, HTATSF1, and GAPDH. (C) Meta plots showing the $\log_2$ fold change signal of RNAPII Ser2P ChIP-seq IP data (normalized to input) across the annotated gene transcripts genes (n = 6694 genes) SF3B1$^{K700E}$ K562 and WT electroporated with siRNA targeting control (in WT and SF3B1$^{K700E}$), DDX46 (in WT), and HTATSF1 (in WT and SF3B1$^{K700E}$). Cells induced with doxycycline (day 0) and electroporated with siRNA on days 3 and 4, followed by ChIP on day 5. (D) Top panel: SF3B1$^{K700E}$ K562 cells transduced with doxycycline-inducible overexpression lentiviral constructs, expressing either for empty vector (EV), N-terminal HA-tagged DDX46 (DDX46-OE), or N-terminal HA-tagged HTATSF1 (HTATSF1-OE). Cell lysates subjected to RT q-PCR to confirm overexpression of DDX46 and HTATSF1 in SF3B1$^{K700E}$. qPCR levels normalized to housekeeping gene, GAPDH. EV levels set as 1. Data plotted are for the mean of two technical replicates based on two independent primer sets. Bottom panel: Western blot analysis confirming overexpression of DDX46 (left) and HTATSF1 (right) in SF3B1$^{K700E}$ +DDX46-OE and SF3B1$^{K700E}$ + HTATSF1-OE cells, respectively, using with antibodies against HA-tag and GAPDH. (E) Gene tracks examples of CDK9 occupancy in WT and SF3B1$^{K700E}$ cells for the indicated genes. The x-axis indicates the chromosome position, and the y-axis represents normalized read density in RPKM. (F) Quantification of readthrough phenotypes in WT and SF3B1$^{K700E}$ cells in TT-TL seq. Distribution of read-through levels (log2 ratio of reads in read-through region vs. gene body) for top 1000 expressed genes. (G) Long-read coverage in the region downstream of PASs is shown for long-reads, in WT and SF3B1$^{K700E}$. Coverage is normalized to the position 100 nt upstream of each PAS (H) Top 15 biological processes of gene ontology (GO) terms sorted by Pvalue < 0.01. Counts of genes and P-value related to each BP term are shown. The y-axis represents BP terms, and the x-axis shows counts of genes were annotated to the BP terms. GO enrichment analysis performed on statistically significant upregulated genes, based on 4-day RNA-seq data, in K562 SF3B1$^{K700E}$ (vs WT).

### SUPPLEMENTAL FIGURE 4

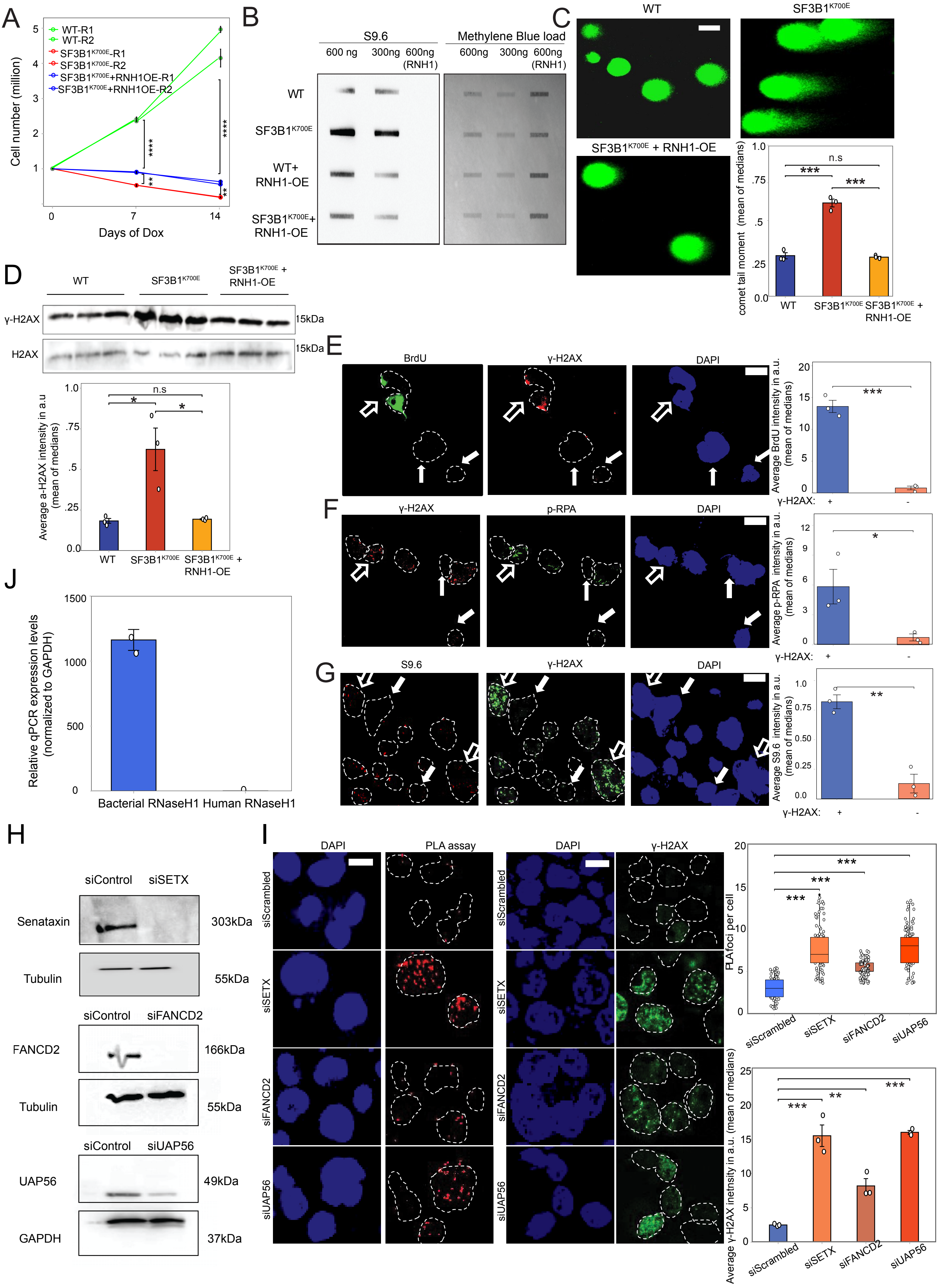

### SUPPLEMENTAL FIGURE 4

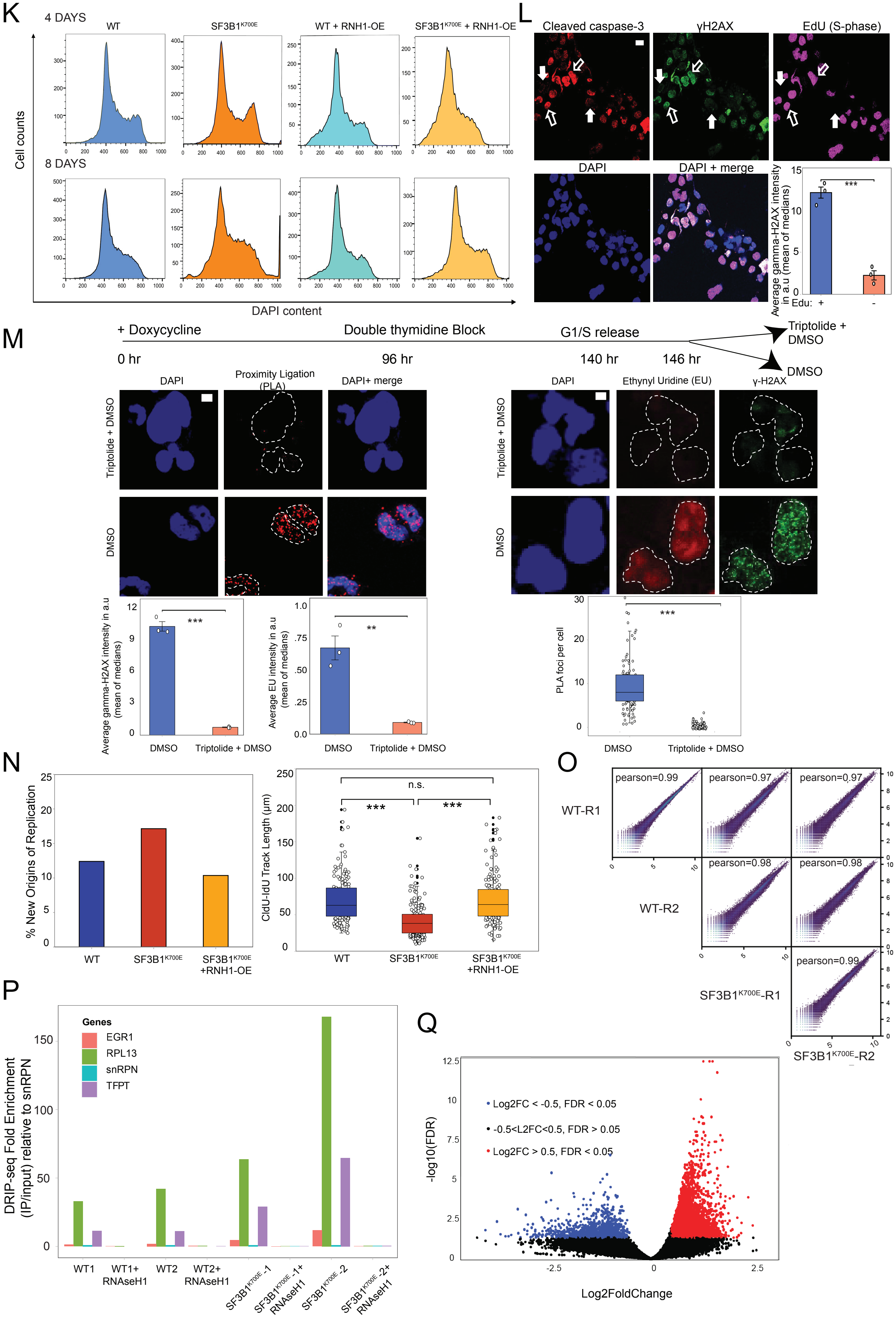

### SUPPLEMENTAL FIGURE 4

R

| Type | | WT (R-loop gain) | SF3B1$^{K700E}$ (R-loop gain) |
|---|---|---|---|
| ncRNA | | 243 (23%) | 1094 (23.9%) |
| Pseudogenes | | 38 (3.6%) | 263 (5.7%) |
| Other/unannotated | | 10 (1%) | 67 (1.4%) |
| Protein-coding | Promoter | 5 (0.4%) | 140 (3%) |
| | Gene body | 391 (30.5%) | 2152 (47%) |
| | Terminator | 10 (1%) | 296 (6.4%) |
| | Intergenic | 421 (40.3%) | 701 (15.3%) |

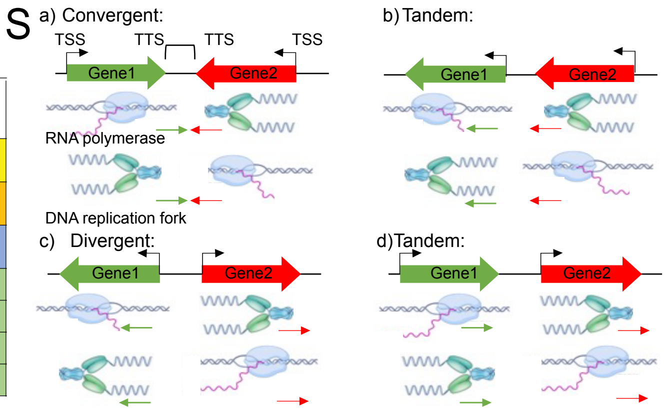


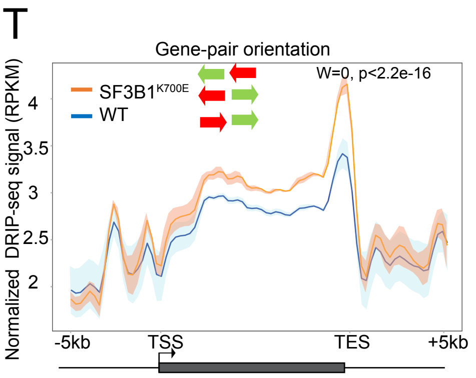


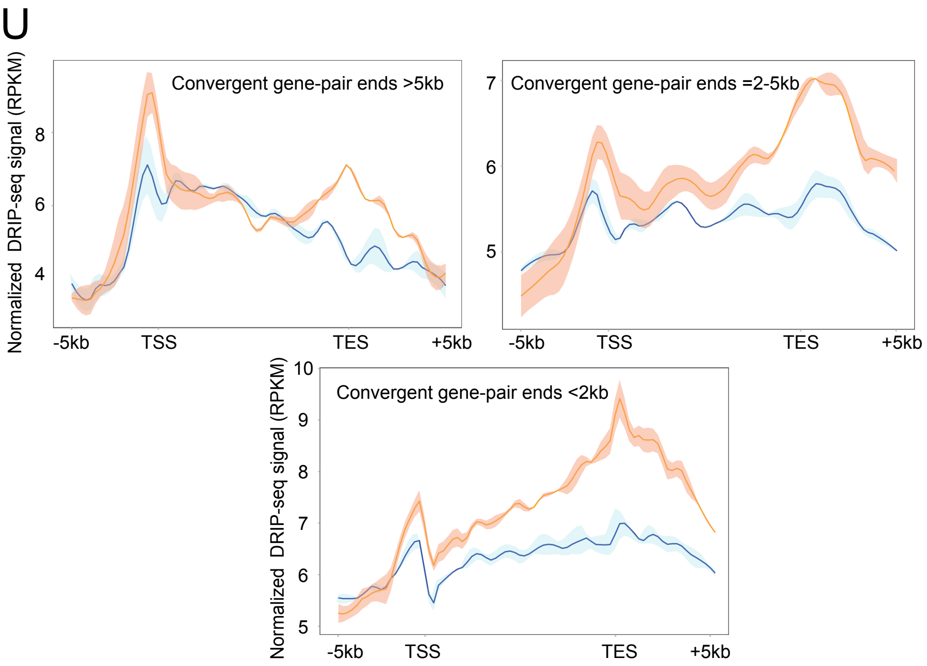


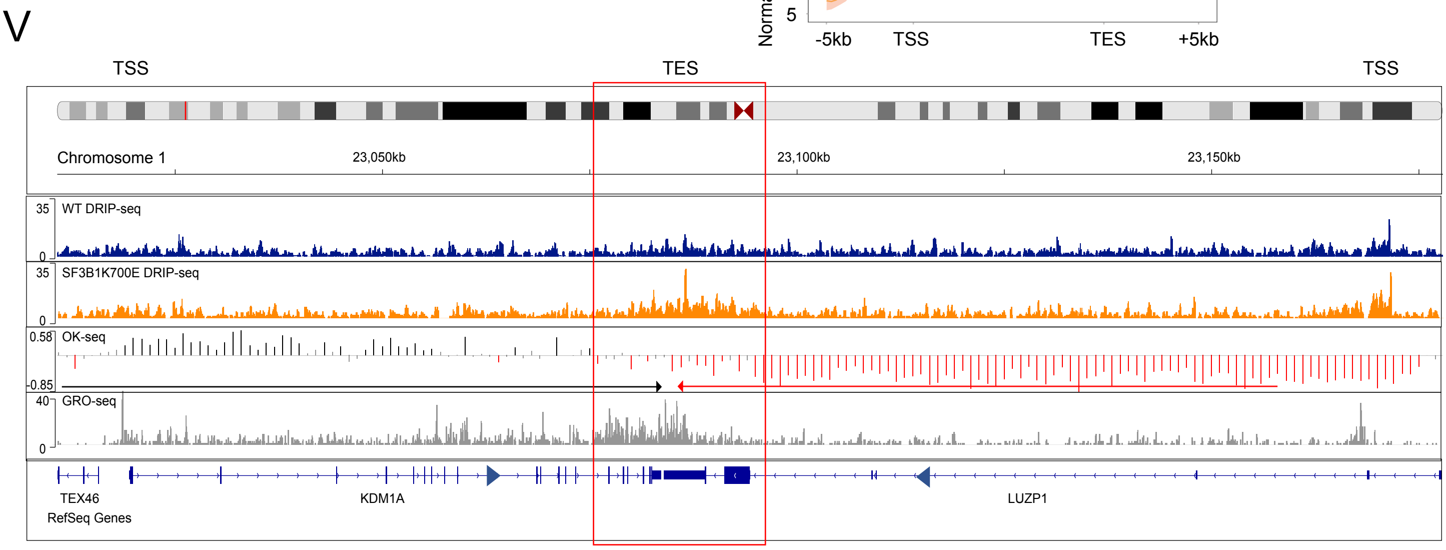


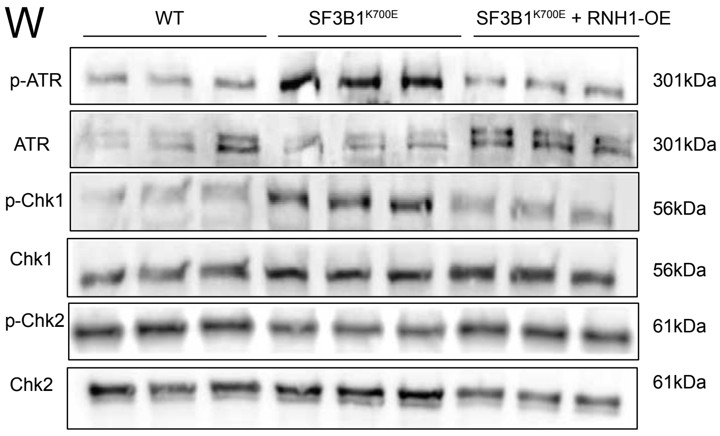

Supplemental Figure 4: **Effect of SF3B1$^{K700E}$ on R-loops, DNA replication dynamics and genome wide R-loop accumulation** (A) WT, SF3B1$^{K700E}$, SF3B1$^{K700E}$ + RNaseH1 (RNH1) overexpression (OE) cells were induced with doxycycline for 4 days (Day 0 at 96 hours post-induction time point), and their cell growth plotted over a course of 14 days ($n = 3$, error bars indicate SEM). Significance assessed using two-tailed unpaired *t* test (***$P_{7\ days(SF3B1^{K700E}\ vs\ SF3B1^{K700E}\ +\ RNH1\text{-}\ OE)} < 0.0001$, **$P_{7\ days(WT\ vs\ SF3B1^{K700E}\ with\ RNH1\text{-}OE)} = 0.0015$, ***$P_{14\ days(SF3B1^{K700E}\ vs\ SF3B1^{K700E}\ +\ RNH1\text{-}OE)} < 0.0001$, **$P_{14\ days(WT\ vs\ SF3B1^{K700E+RNH1\text{-}OE})} = 0.0018$). (B) Nucleic acid extracted from WT (with and without RNaseH1 overexpression) and SF3B1$^{K700E}$ (with and without RNaseH1 overexpression) cells were pre-treated with RNase III (to eliminate dsRNA) and the amount of RNA:DNA hybrids was analyzed using S9.6 antibody. Nucleic acids quantified using Nanodrop™ and loaded at 600 and 300 ng in each replicate. Additionally, 600 ng of RNaseH1 pre-treated nucleic acid extracts included for each replicate, serving as a negative control. Methylene blue staining of the nucleic acid extracts served as loading control. (C) Fluorescence microscopy images and quantification of neutral comet single cell gel electrophoresis assay in treated as in **B**. Data are plotted as the mean of the medians + sem. ($n = 3$ individual experiments; two-tailed, unpaired Student's *t*-test; ***$P < 0.0005$, $^{ns}P > 0.05$). Scale bar, 17 μm. (D) Top panel: Lysates from WT, SF3B1$^{K700E}$, and SF3B1$^{K700E}$ with RNH1 overexpression cells were analyzed by western blotting (WB) for γH2AX, 6 days post induction. H2AX served as loading control. Lower panel: Relative γH2AX band intensities were quantified ($n = 3$). Significance assessed using two-tailed unpaired *t* test (*$P_{\gamma H2AX(SF3B1^{K700E}\ vs\ WT)} = 0.02$, (*$P_{\gamma H2AX(SF3B1^{K700E}\ vs\ SF3B1^{K700E}\ with\ RNH1\text{-}\ OE)} = 0.03$, $^{ns}P_{\gamma H2AX(WT\ vs\ SF3B1^{K700E}\ with\ RNH1\ \text{-}OE)} = 0.57$;error bars indicate SEM). (E) IF images and quantification of BrdU and γH2AX immunofluorescence in SF3B1$^{K700E}$ cells shown as mean ± sem ($n = 3$ separate experiments). a.u: arbitrary units. Significance assessed using one-tailed unpaired *t* test (***$P < 0.0001$). Scale bar = 15 μm. (F) IF images and quantification of pRPA and γH2AX immunofluorescence in SF3B1$^{K700E}$ cells at 6 days post induction, shown as mean ± SEM ($n = 3$ separate experiments). *$P = 0.02$. a.u: arbitrary units. Significance assessed using one-tailed unpaired *t* test. Scale bar = 15 μm. (G) IF images and quantification of S9.6 and γH2AX IF in SF3B1$^{K700E}$ cells at 6 days post induction, shown as shown as mean ± sem ($n = 3$ separate experiments). a.u: arbitrary units. Significance assessed using one-tailed unpaired *t* test **$P = 0.0011$. (H) Western Blot-based confirmation of depletion of Senataxin, FANCD2, and UAP56, by siRNA knockdown experiments. siRNAs and antibodies used are indicated. (I) Epistatic study between SF3B1$^{WT}$ and known R-loop preventing factors in K562 cells. Left Panel: Representative images and quantification of RNAPII NTD +PCNA PLA in WT cells, transfected with scrambled siRNA (siC), and either siRNA against Senataxin (siSETX), siRNA against FANCD2 (siFANCD2), or siRNA against UAP56 (siUAP56). siRNA transfection by electroporation was performed 72 and 96 hours after doxycycline induction, and analysis performed at 120 hours post-doxycycline induction. Data presented as box and whisker plot (n = 3 separate experiments). Scale bar, 17 μm. (Mann-Whitney U test, one-tailed, ***$P_{siC\ vs\ siSETX} < 0.0001$, ***$P_{siC\ vs\ siFANCD2} < 0.0001$, ***$P_{siC\ vs\ siUAP56} < 0.0001$). Right Panel: Representative images and quantification of average gamma-H2AX nuclear intensity in WT cells treated as in **Left Panel.** Data are plotted as the mean of the medians + sem. ($n = 3$ individual experiments;

one-tailed, unpaired Student's $t$-test; Scale bar, 17 µm. (**$P_{siC\ vs\ siSenataxin}$ < 0.0012, ***$P_{siC\ vs\ siFANCD2}$< 0.0057, ***$P_{siC\ vs\ siUAP56}$< 0.0001). (J) SF3B1$^{K700E}$ K562 cells transduced with E. coli RNaseH1-expressing retroviral vectors were subjected to RT q-PCR to confirm bacterial RNaseH1 overexpression. qPCR levels normalized to housekeeping gene, GAPDH. Human RNaseH1 levels are set as 1. Data plotted are for the mean of two biological replicates. (K) Representative images of flow cytometry cell cycle plots in WT, WT + RNH1-OE, SF3B1$^{K700E}$, SF3B1$^{K700E}$ + RNH1-OE cells at 4 days and 8 days following doxycycline induction. Cell counts plotted on Y axis and DAPI content plotted on X axis. (L) IF images and quantification of EdU positive SF3B1$^{K700E}$ cells, 6 days post induction, that are stained for γH2AX, CC3 and EdU. Data comparing gamma-H2AX intensity in the EdU +ve versus EdU -ve cells are plotted as mean of the medians + sem. ($n$ = 3 individual experiments; two-tailed, unpaired Student's $t$-test; **$P$ = 0.0011). Scale bar, 15 µm. a.u: arbitrary units. (M) Schematic of triptolide treatment experiment in G1/S synchronized SF3B1$^{K700E}$ cells. Bottom left panel: Quantification of ethynyl uridine (EU) and gamma-H2AX nuclear intensities in SF3B1$^{K700E}$ cells treated with either Triptolide and DMSO or DMSO alone. IF images of EU and γH2AX fluorescence. Data are plotted as mean of the medians + sem. ($n$ = 3 individual experiments; two-tailed, unpaired Student's $t$-test; ***$P$ < 0.0001). Scale bar, 2 µm. a.u: arbitrary units. Bottom right panel: Quantification of RNAPII + PCNA PLA foci. Foci number per cell are presented as a box-whisker plot ($n$ = 3; two-tailed Mann–Whitney $U$-test). The median values are indicated by black lines. Scale bar, 2 µm. ***$P$ < 0.0001. (N) Left: Percentage of new origins of replication (CldU only tracks) performed in WT, SF3B1$^{K700E}$, and SF3B1$^{K700E}$ + RNH1-OE cells at 6 days post-doxycycline induction. Right panel: Quantification of DNA fiber track length (tracks with both CldU and IdU labeling) performed in WT, SF3B1$^{K700E}$, and SF3B1$^{K700E}$ + RNH1-OE cells at 6 days post-doxycycline induction. Significance assessed using one-tailed Mann-Whitney U test (WT vs SF3B1$^{K700E}$***$P$ value < 0.0001; SF3B1$^{K700E}$ vs SF3B1$^{K700E}$ + RNH1-OE***$P$ value < 0.0001; WT vs SF3B1$^{K700E}$ + RNH1-OE $^{ns}P$ value >0.05). (O) Scatter plots depicting correlation between DRIP-seq samples in WT and SF3B1$^{K700E}$ cells. Values in boxes represent Pearson's correlation coefficients between corresponding samples ($n$ = 2 biological replicates). (WT, r=0.99; SF3B1$^{K700E}$, r=0.99). Values expressed as Log2(gene-expression + 1) (P) RT–qPCR analyses of DRIP IP/Input of EGR1, snRPN, RPL13, and TFPT genes in WT and SF3B1$^{K700E}$ cells, 4 days post doxycycline induction. EGR1 and snRPN represent low R-loop signal genes while TFPT and RPL13 denote high R-loop signal genes. snRPN (IP/input) was set as 1. WT (n=2, WT-R1 and WT-R2) and SF3B1 $^{K700E}$ (n=2, SF3B1$^{K700E}$-R1 and SF3B1$^{K700E}$-R2). Data plotted are for the mean of two technical replicates. (Q) Volcano plot of DRIP-seq differential analysis between SFB1$^{K700E}$ versus WT K562 cells (n = 2 biological replicates, each). Dots represent peaks. Abbreviations used: FDR-false discovery rate, L2FC-log2 fold change. (R) Genome annotation of R-loop gain peaks in WT and SF3B1$^{K700E}$ K562 cells using HOMER software. Promoter: -100bp to +300bp of transcription start site, Gene body: TSS + 300bp to transcription end site (TES), Terminator: TES to +300bp downstream. (S) Categorization of gene pairs based on orientation of transcription between neighboring genes (Genes 1 and 2): (a) Convergent gene-pairs characterized by converging orientation of gene transcription units, (b) and (d) Tandem gene-pairs in which neighboring genes transcribe in the same orientation, and (d) Divergent gene-pairs in which neighboring genes transcribe in diverging orientations. TTSs of convergent gene-pairs represent hot-spots for head-on collisions due to the transcription associated R-loops from Gene1 blocking progression of the oppositely oriented replication fork from Gene2, and vice-versa. (T) Metaplot of the distribution of S9.6 signals (DRIP-seq mean coverage) along 5854 high expressing human non-convergent gene pairs (RPKM > 1000) and flanking regions (±5 kb) in control (blue) and SF3B1$^{K700E}$ (orange) K562 cells. Solid lines represent the averaged signal, and the shaded area represents the upper and lower density limits of the individual replicates in each (n =2, WT; n =2, SF3B1$^{K700E}$). Read densities are calculated as RPKM. Differences in signal intensity at TTS ±2 kb were calculated with the Wilcoxon rank-sum test with continuity correction. (U) Metaplot of the distribution of S9.6 signals (DRIP-seq mean coverage) along high expressing human convergent gene/gene pairs and flanking regions (±5 kb), depending on the distance between their TESs, in control (blue) and SF3B1$^{K700E}$ (orange) K562 cells. Solid lines represent the averaged signal, and the shaded area represent the upper and lower signal limits of the individual replicates in each (n =2, WT; n =2, SF3B1$^{K700E}$). Read densities are calculated as RPKM. Left Panel: >5kb, n=3456 genes; middle panel: 2-5kb, n=799 genes; right panel: <2kb, n=1250 genes. Abbreviations used: TSS-transcription start site, TES-transcription end site. (V) Distribution of DRIP-seq in SF3B1$^{K700E}$ (orange) and WT (blue) K562, Okazaki fragments (OK-seq) in WT K562, and nascent transcription (GRO-seq) signals in WT K562, at two converging genes LUZP1 and KDM1A on chromosome 1 in control K562 cells. (W) Western blot analysis of WT, SF3B1$^{K700E}$ and SF3B1$^{K700E}$ + RNH1-OE cells, 6 days post induction, for phospho-ATM, phospho-Chk1, and phospho-Chk2 levels. ATM, Chk1 and Chk2 used as loading controls, respectively.

### SUPPLEMENTAL FIGURE 5

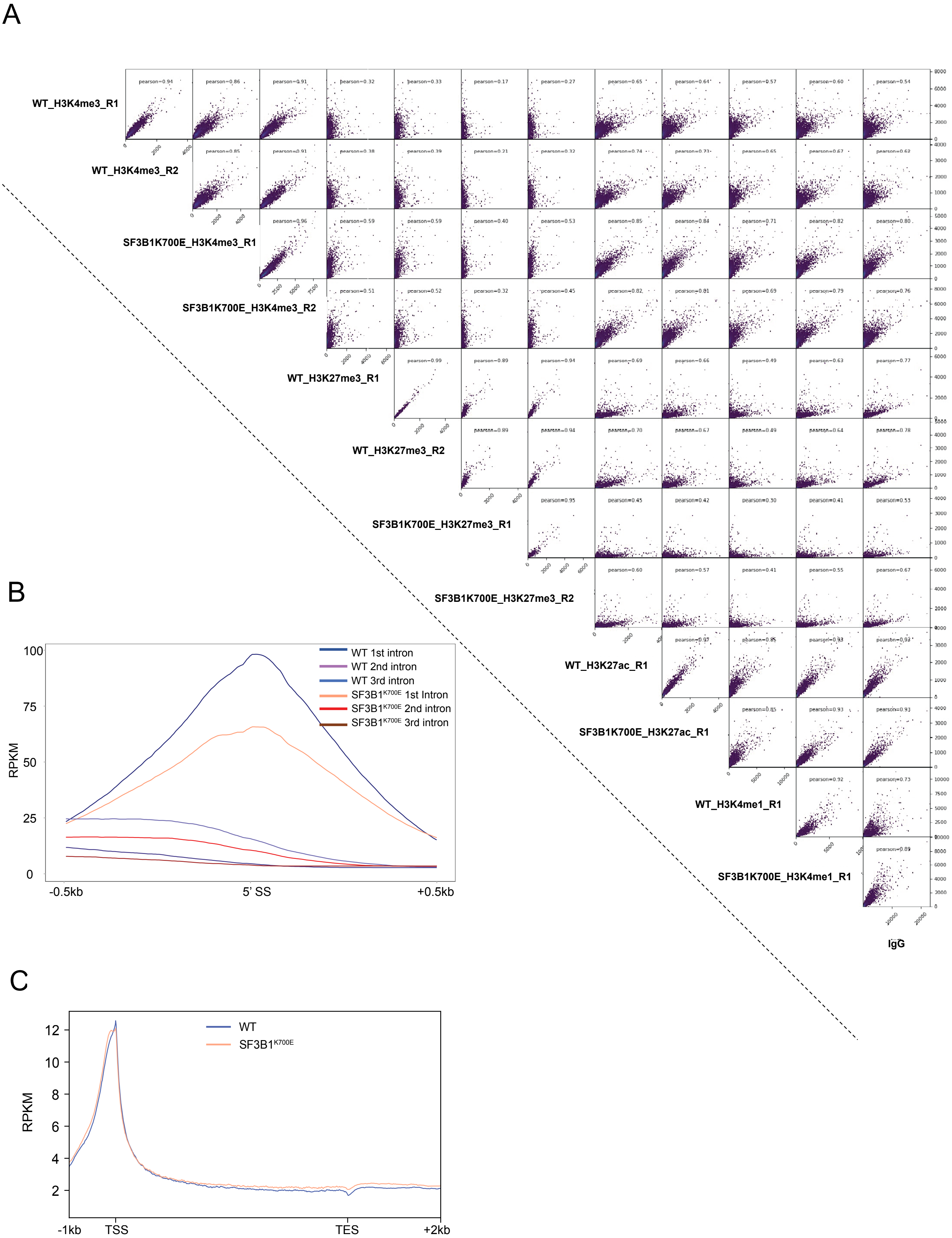

### SUPPLEMENTAL FIGURE 5

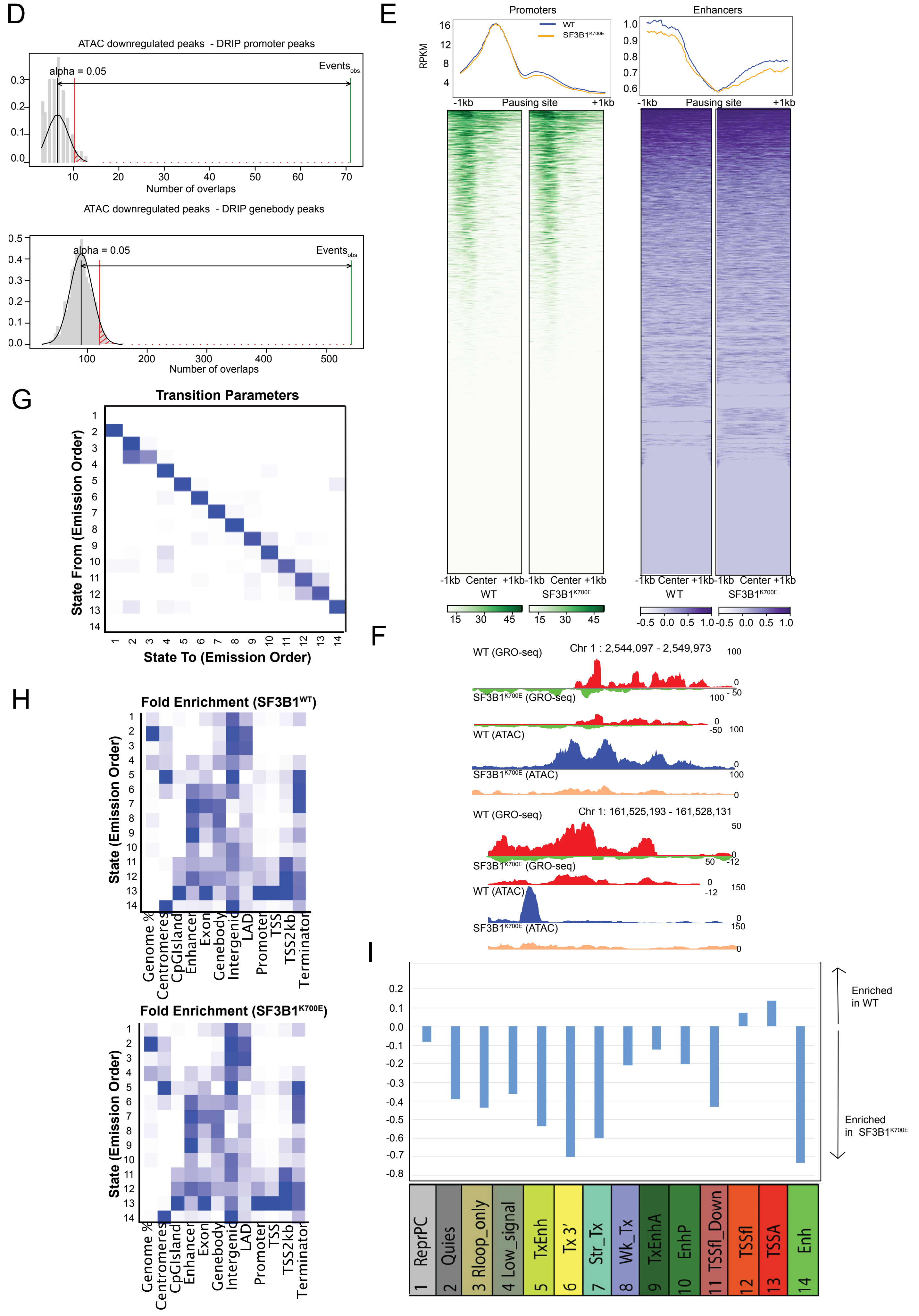

Supplemental figure 5: **Transcription-coupled changes in chromatin accessibility and histone marks due to SF3B1$^{K700E}$ (**A) Scatter plots depicting correlation between the histone mark CutnRun samples in WT and SF3B1$^{K700E}$ samples. Values in boxes represent Pearson's correlation coefficients between corresponding samples. (B) H3K4me3 CUT&RUN enrichment profiles at 5' splice sites (5'SS) of first, second, and third introns in WT and SF3B1$^{K700E}$. Y-axis shows the RPKM. (C) Metagene plot showing the normalized H3K27ac CUT&RUN levels over 6694 expressed genes in WT (blue) and SF3B1$^{K700E}$ (orange) (n =1, WT; n =1, SF3B1$^{K700E}$). Read densities are calculated as RPKM. (D) RegioneR-based colocalization analysis of downregulated ATAC peaks with upregulated R-loop regions (SF3B1$^{K700E}$/WT) at promoter and gene body regions. (E) Metaplots and heatmaps of RPKM-normalized GRO-seq signal for antisense transcription at promoters (left) and enhancers (right) ranked by decreasing occupancy. (F) Representative genome browser track examples of GRO-seq densities and ATAC-seq in WT (blue) versus SF3B1$^{K700E}$ (orange), for the indicated putative enhancers. The x-axis indicates the chromosome position, and the y-axis represents normalized RPKM. (G) Transition parameters for the 14-state model derived by ChromHMM for WT and SF3B1$^{K700E}$ cells. (H) Overlap of different genomic features (Centromeres, CpG island, Putative Enhancer, Exons, Gene body, Intergenic, laminB lads, Promoter, TSS, TSS+2kb downstream, and terminator regions with chromatin state calls in WT (left) and SF3B1$^{K700E}$ (right). The fold enrichments are calculated as the ratio between observed and expected number of genomic bins for each overlap. The color intensities are normalized within each column between its minimum value (white) and its maximum value (blue). (I) Log2 ratios between the number of total genomic bins for each chromatin state in WT and SF3B1$^{K700E}$ cells (WT/SF3B1$^{K700E}$). Log2ratio plotted on Y-axis and chromatin state plotted on X-axis. A log2ratio above zero indicates a higher genomic bin count in the WT for a particular chromatin state. Conversely, a log2ratio below zero indicates a lower genomic bin count in SF3B1$^{K700E}$ for a particular chromatin state.

### SUPPLEMENTAL FIGURE 6

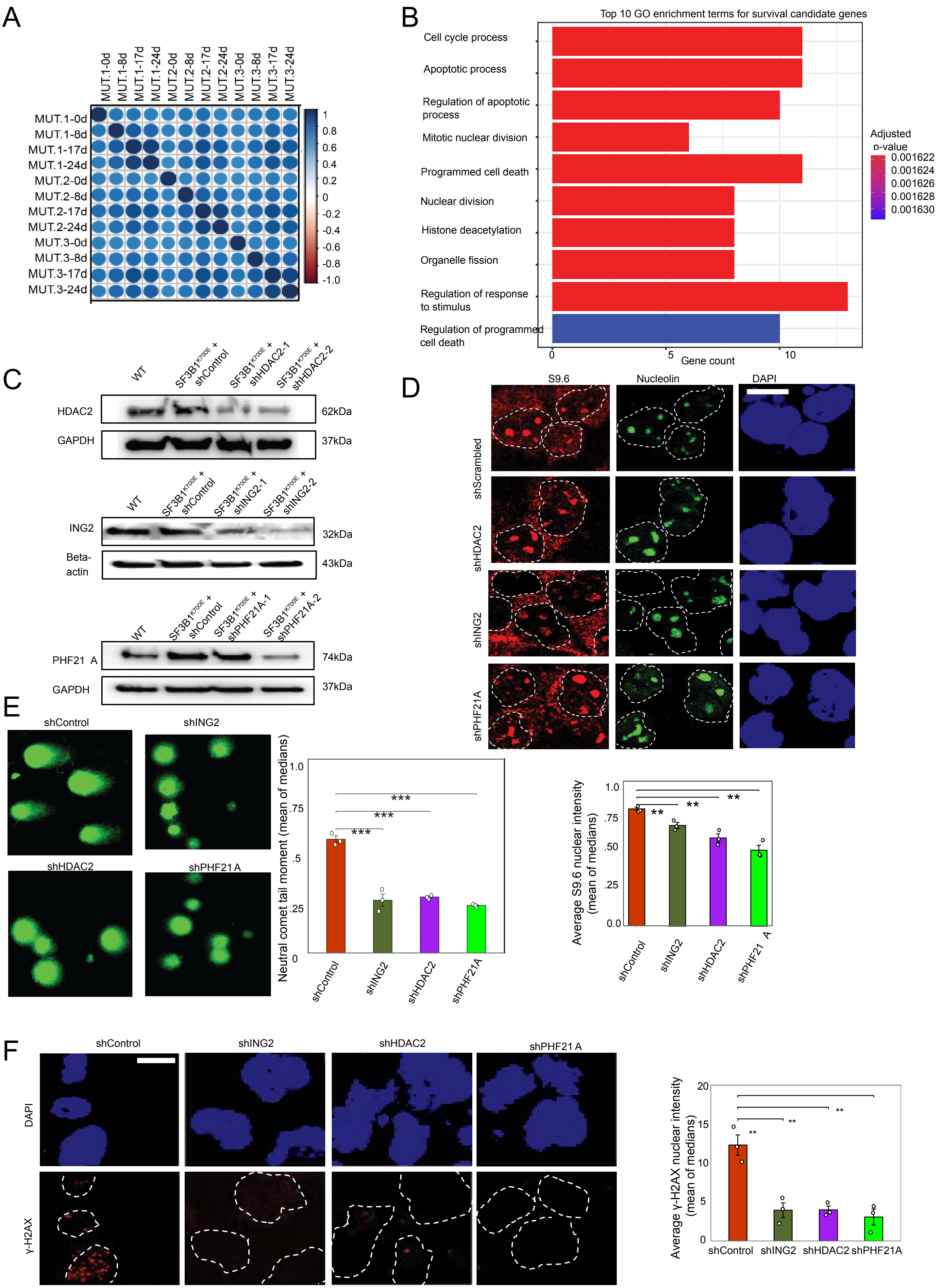

### SUPPLEMENTAL FIGURE 6

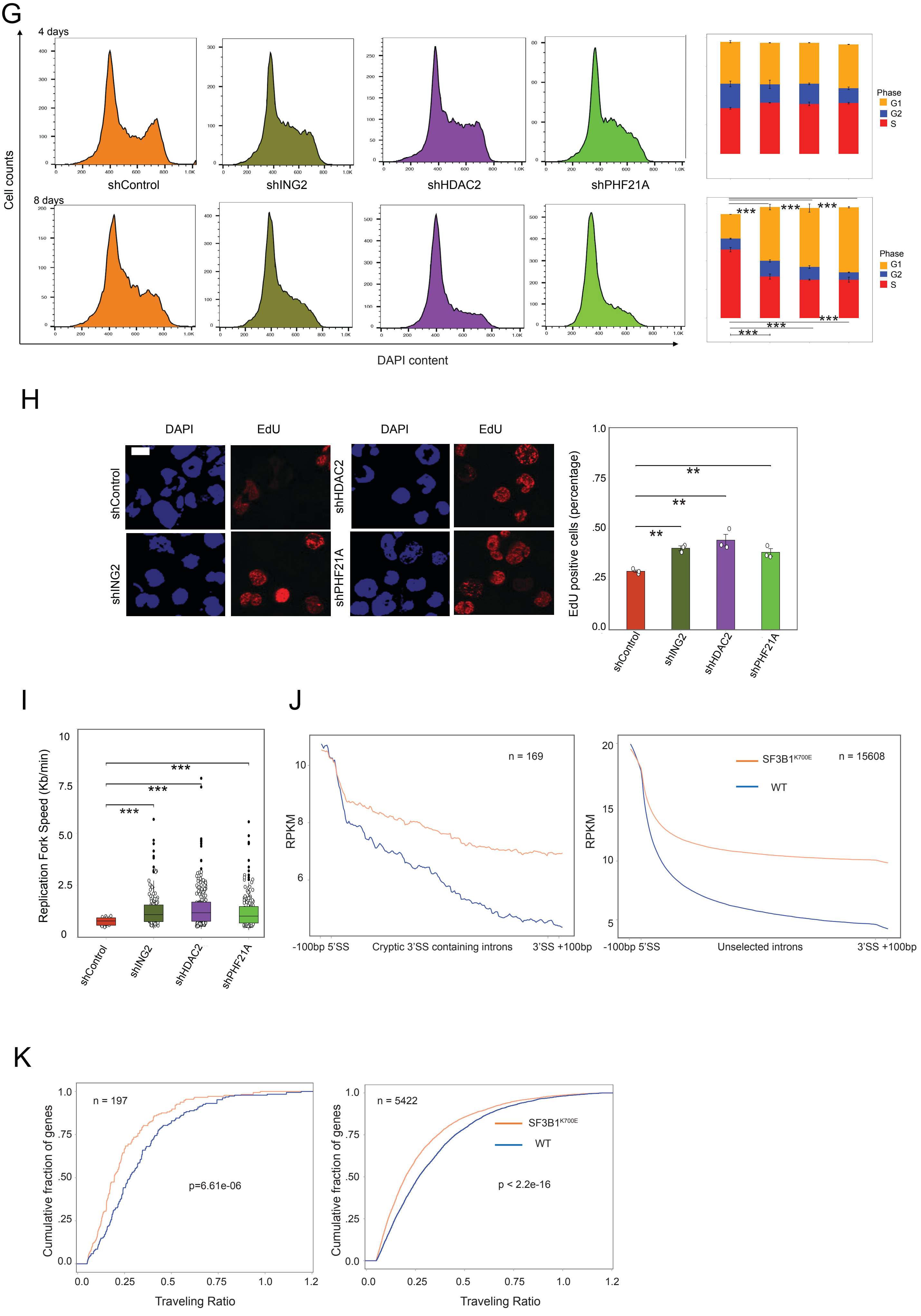

Supplemental Figure6: **An epigenetic shRNA rescue screen identifying key functional pathways important to SF3B1$^{K700E}$-related cell survival** (A) Plot correlation heatmap of SF3B1$^{K700E}$ (n=3 per genotype; labeled as MUT.1, MUT.2, MUT.3) at different time points of gDNA analysis (days 0, 8, 17, 24). (B) GO term enrichment analysis of top 20 survival candidates. Enriched GO terms (FDR <0.05) grouped by biologic process function. X axis represents the number of genes representing a particular GO term function. (C) Western blot confirmation of shRNA mediated knockdown of protein levels. Cells transduced with shScr or two independent shRNAs knocking down (KD) HDAC2 (shHDAC2-1 and shHDAC2-2), ING2 (shING2-1 and shING2-2), and PHF21A (shPHF21A-1 and shPHF21A-2) were subjected to western blotting with respective antibodies. Of note, subsequent analysis was restricted to the stronger shPHF21A-2 clone based on effectiveness in depleting PHF21A protein levels. GAPDH used as loading control for testing HDAC2 and PHF21A KDs, and beta-actin used as loading control for testing ING2 KD. (D) Representative images and quantification of nuclear S9.6 intensity in SF3B1$^{K700E}$ + shScrambled (shScr), SF3B1$^{K700E}$ + shHDAC2-1, SF3B1$^{K700E}$ + shING2-1 , SF3B1 $^{K700E}$ + shPHF21A at 6 days post induction. Data are plotted as mean of the medians + sem. (*n* = 3 individual experiments; two-tailed, unpaired Student's *t*-test; $**P_{(SF3B1^{K700E} + shHDAC2 \text{ vs } SF3B1^{K700E} + shScrambled}$ = 0.0031; $**P_{(SF3B1^{K700E} + shING2 \text{ vs } SF3B1^{K700E} + shScrambled}$ = 0.004; $**P_{(SF3B1^{K700E} + shPHF21A \text{ vs } SF3B1^{K700E} + shScrambled}$ = 0.0013). Scale bar,15 µm. a.u: arbitrary units. (E) Fluorescence microscopy images of neutral comet single cell gel electrophoresis assay in SF3B1$^{K700E}$ + shScrambled (shScr), SF3B1$^{K700E}$ + shHDAC2, SF3B1$^{K700E}$ + shING2, SF3B1$^{K700E}$ + shPHF21A cells at 6 days post induction. Data are plotted as mean of the medians + sem. (*n* = 3 individual experiments; two-tailed, unpaired Student's *t*-test; $**P_{(SF3B1^{K700E} + shScr \text{ vs } SF3B1^{K700E} + shING2)}$ = 0.001; $***P_{(SF3B1^{K700E} + shScr \text{ vs } SF3B1^{K700E} + shHDAC2)}$ <0.001; $**P_{(SF3B1^{K700E} + shScr \text{ vs } SF3B1^{K700E} + shPHF21A)}$ = 0.004). Scale bar, 17 µm. (F) IF images and quantification of γH2AX fluorescence in SF3B1$^{K700E}$ + shScrambled (shScr), SF3B1$^{K700E}$ + shHDAC2-1, SF3B1$^{K700E}$ + shING2-1, SF3B1$^{K700E}$ + shPHF21A at 5 days post induction. Data are plotted as mean of the medians + sem. (*n* = 3 individual experiments; two-tailed, unpaired Student's *t*-test; $**P_{(SF3B1^{K700E} + shHDAC2 \text{ vs } SF3B1^{K700E} + shScrambled}$ = 0.0038; $**P_{(SF3B1^{K700E} + shING2 \text{ vs } SF3B1^{K700E} + shScrambled}$ = 0.006; $**P_{(SF3B1^{K700E} + shPHF21A \text{ vs } SF3B1^{K700E} + shScrambled}$ = 0.005). Scale bar,15 µm. a.u: arbitrary units. (G) Representative images and quantification of flow cytometry cell cycle plots in SF3B1$^{K700E}$ + shScrambled (shScr), SF3B1$^{K700E}$+ shHDAC2, SF3B1$^{K700E}$ + shING2, SF3B1$^{K700E}$ + shPHF21A cells at 4 days and 8 days following doxycycline induction. The Watson pragmatic fitting algorithm was used for cell cycle analysis. Data presented as mean ± sem (n=3 separate experiments). Significance assessed using ANOVA followed by Sidak'stest: (%G2/$M_{4days:SF3B1^{K700E} + shScr \text{ vs } SF3B1^{K700E} + shPHF21A}$ =0.04; %$G1_{8days:SF3B1^{K700E} + shScr \text{ vs } SF3B1^{K700E} + shING2}$****Adjusted $P < 0.0001$; %$G1_{8days:SF3B1^{K700E} + shScr \text{ vs } SF3B1^{K700E} + shHDAC2}$****Adjusted $P < 0.0001$; %$G1_{8days:SF3B1^{K700E} + shScr \text{ vs } SF3B1^{K700E} + shPHF21A}$****Adjusted $P < 0.0001$; %$S_{8days:SF3B1^{K700E} + shScr \text{ vs } SF3B1^{K700E} + shING2}$****Adjusted $P < 0.0001$; %$S_{8days:SF3B1^{K700E} + shScr \text{ vs } SF3B1^{K700E} + shHDAC2}$****Adjusted $P < 0.0001$; %$S_{8days:SF3B1^{K700E} + shScr \text{ vs } SF3B1^{K700E} + shPHF21A}$****Adjusted $P < 0.0001$) (H) Quantification of percentage of cells incorporating EdU ($EdU^+$) in SF3B1$^{K700E}$ + shScrambled (shScr), SF3B1$^{K700E}$ + shHDAC2, SF3B1$^{K700E}$ + shING2 , SF3B1$^{K700E}$ + shPHF21A cells at 6 days following doxycycline induction. Percent EdU positive cell data are plotted as mean of the medians + sem. (*n* = 3 individual experiments; two-tailed, unpaired Student's *t*-test; $**P_{(SF3B1^{K700E} + shScr\ vs\ SF3B1^{K700E} + shING2)} = 0.0011$; $**P_{(SF3B1^{K700E} + shScr\ vs\ SF3B1^{K700E} + shHDAC2)} = 0.004$; $**P_{(SF3B1^{K700E} + shScr\ vs\ SF3B1^{K700E} + shPHF21A)} = 0.0062$). Scale bar, 15 μm. (I) Quantification of fork velocity performed in $SF3B1^{K700E}$ + shScrambled (Tangerine), $SF3B1^{K700E}$ + shING2 (olive green), $SF3B1^{K700E}$ + shHDAC2 (blue violet), $SF3B1^{K700E}$ + shPHF21A (neon green) cells at 6 days post induction. Data are shown as a box plot (*n* = 2). Significance assessed using one-tailed Mann-Whitney U test ($SF3B1^{K700E}$ + shScr vs $SF3B1^{K700E}$+ shING2, ***$P$ value < 0.001; $SF3B1^{K700E}$ + shScr vs $SF3B1^{K700E}$ + shHDAC2, ***$P$ value < 0.001;  $SF3B1^{K700E}$ + shScr vs $SF3B1^{K700E}$ + shPHF21A,**$P$ value =0.001). (J) Read coverage in the introns undergoing cryptic 3'SS splicing [determined from analysis of patient sample data] (left) versus unselected introns (right). Coverage is normalized to the position 100 nt upstream of each 5'SS. Orange line indicates $SF3B1^{K700E}$ and the blue line indicates WT. (K) The cumulative distribution function plot of the TR distribution, calculated as an average of two replicates) in WT and $SF3B1^{K700E}$ cells depending on genes with introns undergoing cryptic 3'SS splicing (determined from analysis of MDS patient sample data) versus unselected genes. Significance determined using two sample K.S. test.

### SUPPLEMENTAL FIGURE 7

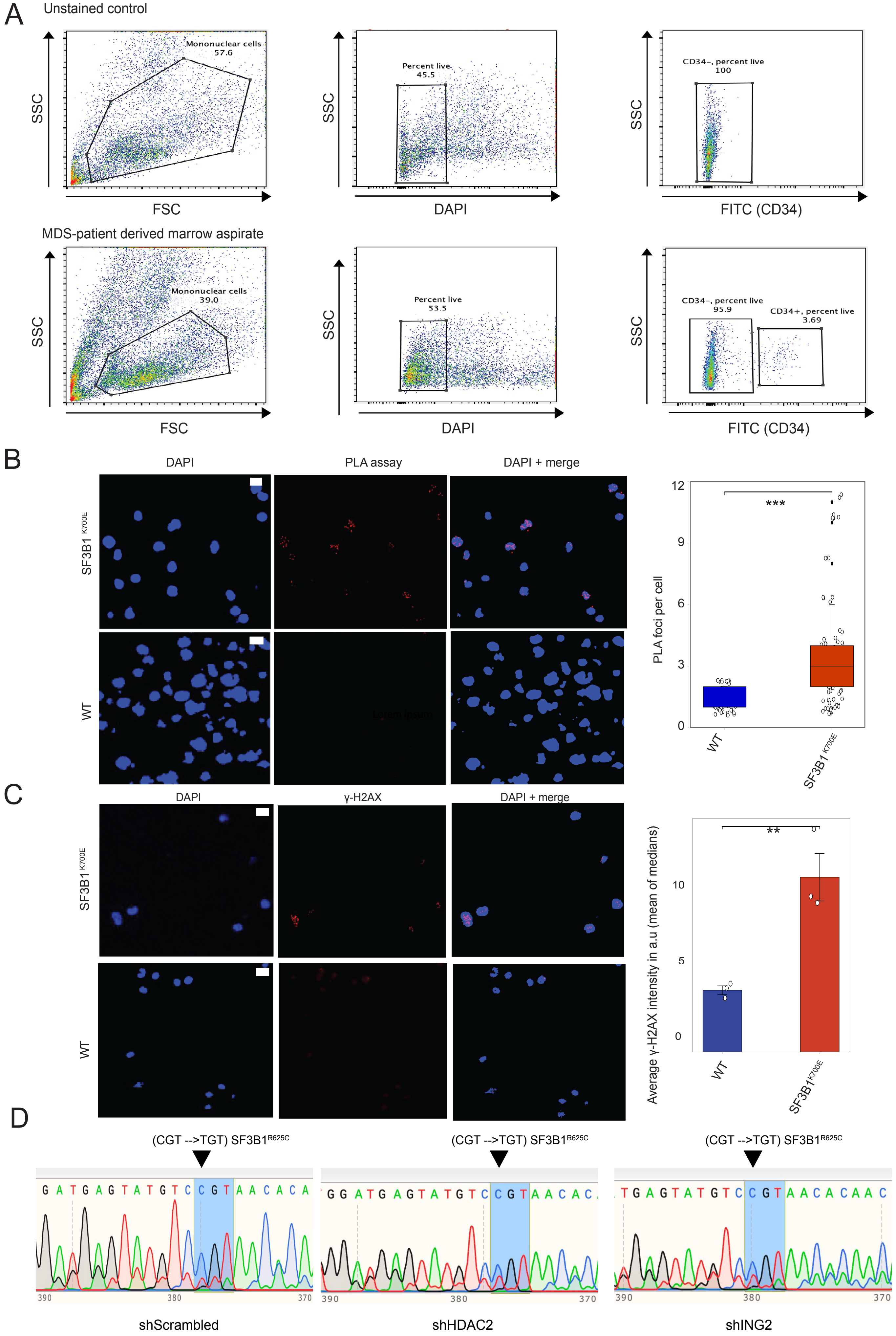

### SUPPLEMENTAL FIGURE 7

E

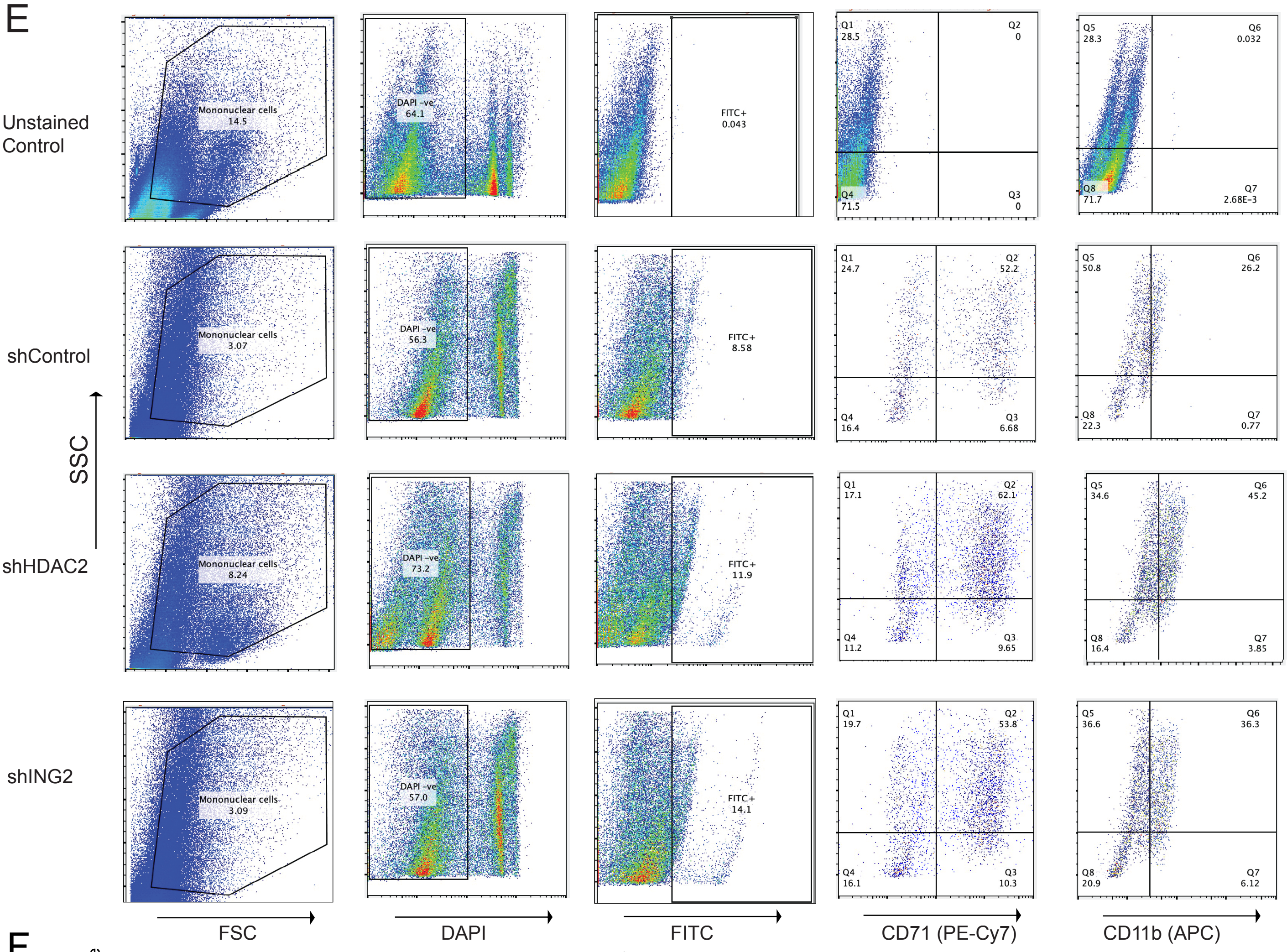


F

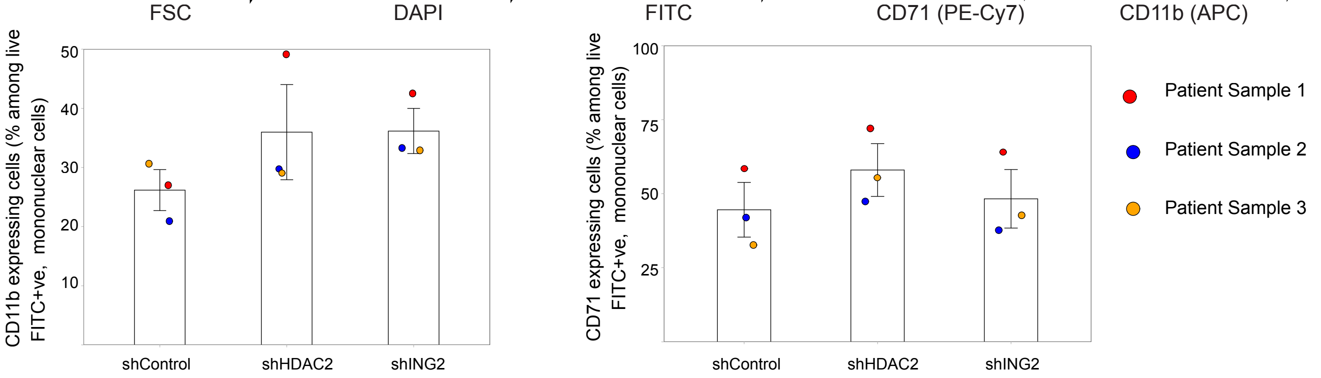


G

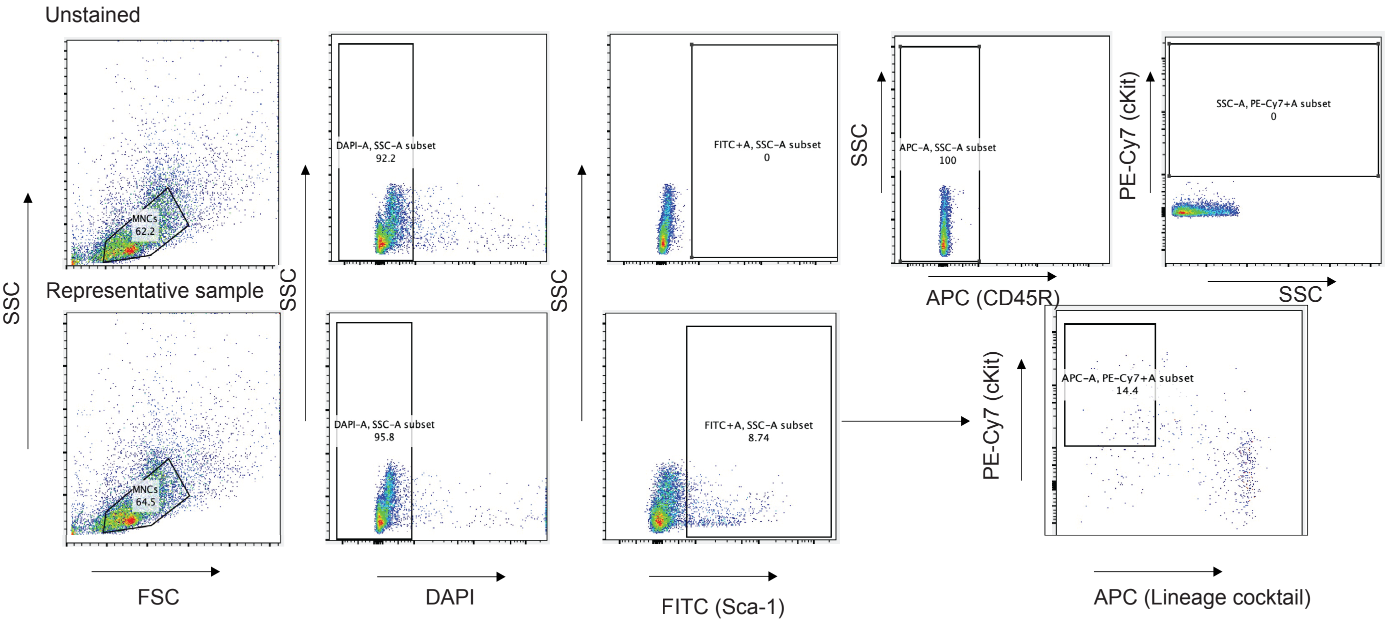

### SUPPLEMENTAL FIGURE 7

H

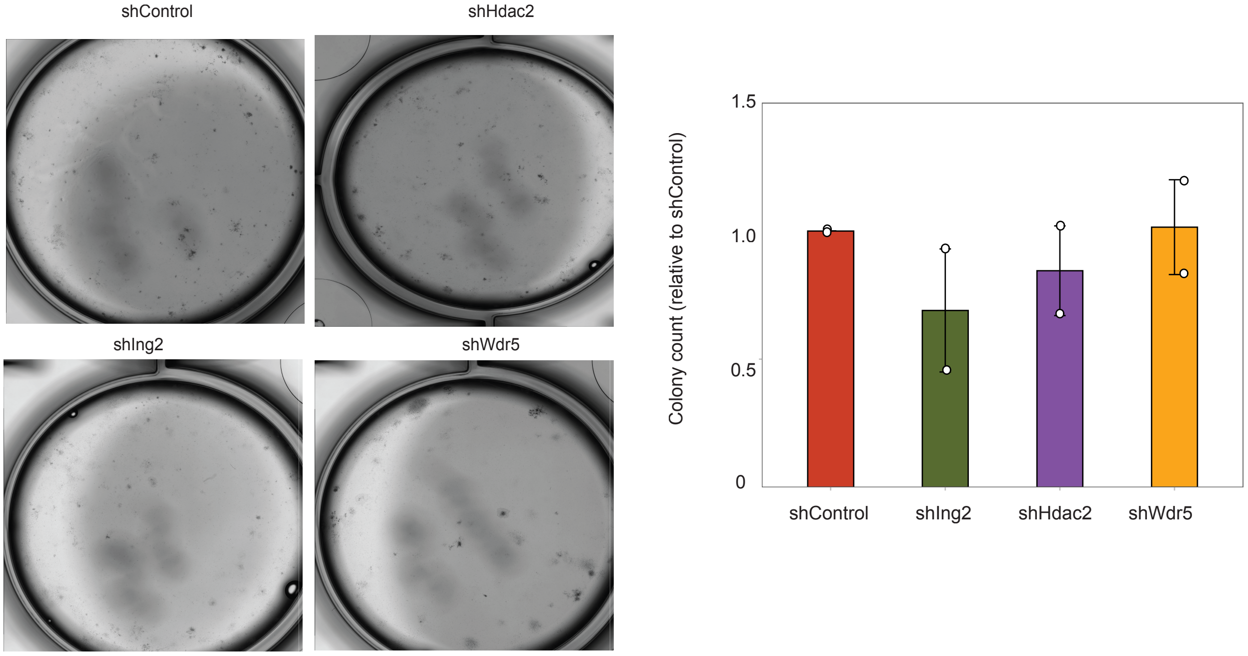


I

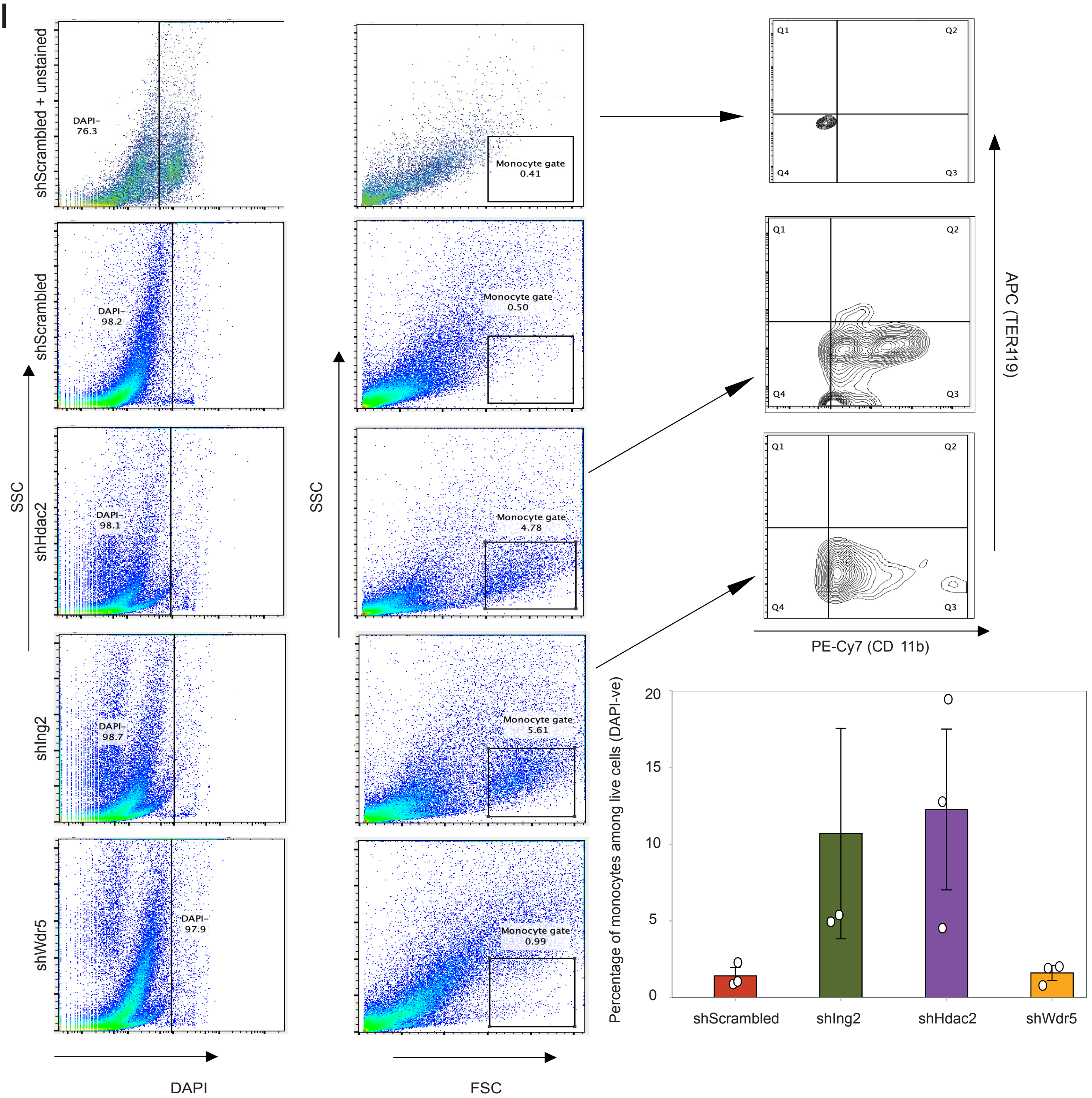

Supplemental Figure 7: **Targeting H3K4me pathway in SF3B1-mutant MDS.** (A) Gating strategy used throughout the experiments for flow cytometric analysis of CD34 surface expression markers on mononuclear cells isolated from bone marrow aspirates. Mononuclear cells were stained with DAPI and acquired in the flow cytometer. Events were gated based on forward (FSC) and side (SSC) scatter characteristics and then gated on live cells (DAPI -). Live cells were then gated on CD34 surface marker expression. Top panel: representative example of gating strategy in CD34 unstained negative control cell population; Bottom panel: representative example of gating strategy in one of the SF3B1-mutant MDS marrow aspirate derived mononuclear cells. (B) Quantification of RNAPII NTD + PCNA foci (by PLA assay) in WT and SF3B1-mutant MDS derived cells. Foci number per cell are presented as a scatter plot ($n = 3$ independent experiments each; two-tailed Mann–Whitney *U*-test). The median values are indicated by black lines. Scale bar, 2 μm. $***P < 0.0001$. (C) Quantification of gamma-H2AX intensity in WT and SF3B1-mutant MDS derived cells. Gamma-H2AX intensity presented as a bar plot. Mean + sem ($n = 3$ individual experiments). Scale bar, 2 μm. $***P < 0.0001$. (D) Representative example showing chromatograms of PCR amplicons around the SF3B1 R625 locus. Patient-derived SF3B1$^{R625C}$ MDS CD34$^{+}$ cells were isolated and grown in methylcellulose medium for 14 days after being transduced with lentiviral constructs targeting non-specific control (shScrambled), HDAC2 (shHDAC2), ING2 (shING2). DNA was isolated from the cell colonies at 14 days for the respective conditions and PCR performed on the extracted DNA to amplify the region around the SF3B1 R625 locus. Highlighted in blue is the R625 locus, showing the presence of TGT mutation (R→C). (E) Gating strategy used throughout the experiments for flow cytometric analysis of C11b and CD71 surface expression markers on nucleated cells isolated from SF3B1-mutant MDS derived colonies grown in methylcellulose medium for 14 days. Mononuclear cells were stained with DAPI and acquired in the flow cytometer. Events were first gated based on forward (FSC) and side (SSC) scatter characteristics and then gated on live cells (DAPI - ). Live cells were then gated on the FITC channel for GFP-expressing cells. Flow cytometric evaluation on the GFP+ cells was performed to evaluate for C11b (APC) and CD71 (PE-Cy7) surface expression. Panels from top to bottom in the following order: unstained control, SF3B1-mutant MDS derived cells transduced with GFP-expressing shControl, shHDAC2, and shING2 lentiviral vectors, respectively. (F) Percentage of CD11b and CD71 expressing cells (among live, mononuclear cells) in shControl, shHDAC2, and shING2 settings. Data presented as the mean of 3 individual patient samples. Each datapoint represents a patient sample. Statistical comparisons deferred due to high variability across biological replicates. (G) Gating strategy used throughout the experiments for flow cytometric analysis and isolation of cKit+Sca-1+Lin- cells isolated from bone marrow aspirates. Mononuclear cells were stained with DAPI and acquired in the flow cytometer. Events were gated based on forward (FSC) and side (SSC) scatter characteristics and then gated on live cells (DAPI - ). Live cells were then gated on Sca-1 surface marker expression (FITC). Sca-1 expressing cells were then gated on cKit (PE-Cy7) and Lineage cocktail (APC) marker expressions. Top panel: representative example of gating strategy in unstained negative control cell population; Bottom panel: representative example of gating strategy in one of the marrow mononuclear cells to isolate cKit+Sca-1+Lin- cells. (H) Gating strategy used throughout the experiments for flow cytometric analysis of C11b and CD71 surface expression markers on nucleated cells isolated from murine Sf3b1$^{K700E}$ cKit+Sca-1+Lin- cell- derived colonies grown in methylcellulose medium for 14 days. Mononuclear cells were stained with DAPI and acquired in the flow cytometer. Events were first gated on live cells (DAPI-) and then based on forward (FSC) and side (SSC) scatter characteristics. An increase in the percentage of events was noted in the 'monocyte' gate (low SSC, high FSC), for the shHdac2 and shIng2 settings. Flow cytometric evaluation on the cells, evaluating for C11b (PE-Cy7) and CD71 (APC) surface expression, showed that the events in this gate indeed corresponded to CD11b+ monocytes. Panels from top to bottom in the following order: unstained control, Sf3b1$^{K700E}$ cKit+Sca-1+Lin- cells transduced with shScrambled, shHdac2, shIng2, and shWdr5 expressing lentiviral vectors, respectively. (I) Metaplot of WT and Sf3b1$^{K700E}$ ATAC-seq signal profiling changes in the nucleosome occupancy among 1024 high-expressing genes (TPM>1000) in WT and Sf3b1$^{K700E}$ cKit+Sca-1+Lin- murine cells.

### SUPPLEMENTAL FIGURE 8

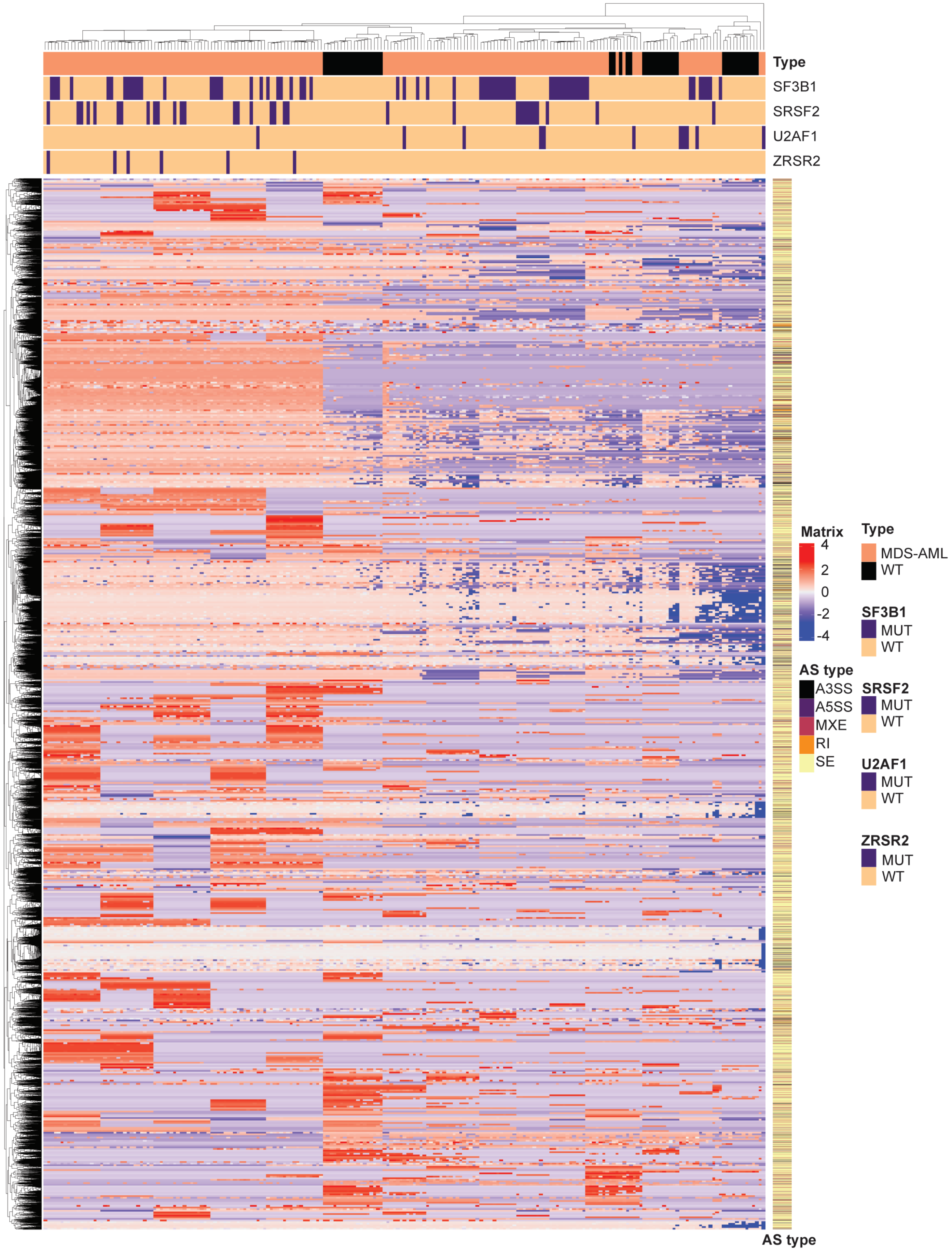

Supplemental FigS8: **Unsupervised clustering of alternative splicing events** across 172 MDS/AML patients, and 45 normal samples from 3 individual datasets[31 33 32]. Alternative splicing events were first determined by rMATS package individually for each sample. Events which were present at PSI (percentage spliced in or fraction of mRNA that corresponds to the novel isoform) value of 0.1 or higher in at least 10% of the total sample were selected (total 137,435 events) and unsupervised clustering performed by ComplexHeatmap R package. Mutational data for SF3B1, U2AF1, SRSF2 and ZRSR2 are shown on the top panel. Alternative splicing event type are show on the right-side column (A3SS = alternative 3’ Splice Site, A5SS = alternative 5’ splice site, MXE = mutually exclusive exon, SE = skipped exon, RI = retained intron)
